# Interplay between historical and current features of the cityscape in shaping the genetic structure of the house mouse (*Mus musculus domesticus*) in Dakar (Senegal, West Africa)

**DOI:** 10.1101/557066

**Authors:** Claire Stragier, Sylvain Piry, Anne Loiseau, Mamadou Kane, Aliou Sow, Youssoupha Niang, Mamoudou Diallo, Arame Ndiaye, Philippe Gauthier, Marion Borderon, Laurent Granjon, Carine Brouat, Karine Berthier

**Author notes:** These two authors contributed equally to this work (Carine Brouat, Karine Berthier).

## Abstract

Population genetic approaches may be used to investigate dispersal patterns of species living in highly urbanized environment in order to improve management strategies for biodiversity conservation or pest control. However, in such environment, population genetic structure may reflect both current features of the cityscape and urbanization history. This can be especially relevant when focusing on exotic commensal rodents that have been introduced in numerous primary colonial European settlements. Accounting for spatial and temporal cityscape heterogeneity to determine how past and recent demographic events may interplay to shape current population genetic structure of synanthropic rodents may provide useful insights to manage their populations. In this study, we addressed these issues by focusing on the house mouse, *Mus musculus domesticus*, in Dakar, Senegal, where the species may have been introduced as soon as Europeans settled in the middle of the nineteenth century. We examined genetic variation at one mitochondrial locus and 15 nuclear microsatellite markers from individuals sampled in 14 sampling sites representing different stages of urbanization history and different socio-economic environments in Dakar. We used various approaches, including model-based genetic clustering and model-free smoothing of pairwise genetic estimates. We further linked observed spatial genetic patterns to historical and current features of Dakar cityscape using random forest and Bayesian conditional autoregressive models. Results are consistent with an introduction of the house mouse at colonial time and the current genetic structure exhibits a gradient-like pattern reflecting the historical process of spatially continuous expansion of the city from the first European settlement. The genetic patterns further suggest that population dynamics of the house mouse is also driven by the spatial heterogeneity of the current cityscape, including socio-economics features, that translate in habitat quality. Our results highlight the potential importance of accounting for past demographic events to understand spatial genetic patterns of nonnative invasive commensal rodents in highly urbanized environment.

## Introduction

Urbanization is a major environmental change and represents the land use that is the fastest growing (Bloom et al., 2008), especially in developing countries (Chen et al., 2014; Cohen, 2006). Urbanization is considered as an extreme type of habitat fragmentation that can lead to spatial isolation of small subpopulations or even to local extinction (McKinney, 2006; Munshi-South et al., 2016). As a consequence, genetic drift is expected to be enhanced for numerous species, leading to a decrease of genetic diversity within local subpopulations and to an increase of genetic differentiation between them (Johnson and Munshi-South, 2017). However, the situation can be quite different for species that are not limited to remnant patches of native habitat and can easily inhabit, and disperse through, a diversity of urban infrastructures (Combs et al., 2018a,b).

Invasive rats and mice are archetypal commensals and a major concern for public health, food security and biodiversity all around the world (Angel et al., 2009; Meerburg et al., 2009; Parsons et al., 2017; Singleton et al., 2003, 2015). Rodent control actions are costly, and classical control protocols, relying mostly on poisoning, have proven to be quite inefficient in reducing urban rodent densities in the long term (Parsons et al., 2017; Richardson et al., 2017). It is now recognized that the development of more efficient management strategies requires a better understanding of urban rodent evolutionary ecology (Byers et al., 2019; Combs et al., 2018a; Feng and Himsworth, 2014; Makundi and Massawe, 2011; Parsons et al., 2017; Singleton et al., 1999) and notably of variations in gene flow (Combs et al., 2018a,b; Gardner-Santana et al., 2009; Johnson and Munshi-South, 2017; Kajdacsi et al., 2013; Parsons et al., 2017; Richardson et al., 2017). For instance, identifying genetic clusters can allow defining meaningful spatial units for control actions (Combs et al., 2018a; Kajdacsi et al., 2013; Richardson et al., 2017). As anticoagulant resistance can rapidly reach high frequency within local rodent populations (Berny et al., 2018), knowledge on spatial patterns of gene flow can also help in predicting the risk at which resistance alleles can spread across a cityscape (Haniza et al., 2015). So far, population genetic studies of urban rodents have mostly focused on the brown rat (*Rattus norvegicus*). Several of these studies showed that gene flow occurs at a sufficient rate to limit genetic drift and that spatial genetic variation tends to be organized under isolation-by-distance (IBD) resulting from both the spatially restricted dispersal and social behavior of this species (Combs et al., 2018a,b; Gardner-Santana et al., 2009). Beyond this general trend, spatially explicit analyses have also revealed sharp genetic discontinuities at relatively small spatial scales (Combs et al., 2018b; Richardson et al., 2017). Such genetic breaks have been attributed to physical barriers to dispersal (waterway in Baltimore, USA – Gardner-Santana et al., 2009; roads and topography in Pau de Lima, Salvador, Brazil – Kajdacsi et al., 2013; Richardson et al., 2017) or spatial heterogeneity in habitat quality that impacts local population size (resource availability varying with socioeconomic features of urban infrastructures, Manhattan, USA – Combs et al., 2018b). The need to integrate socioeconomic features of the urban environment in population dynamic and genetic studies of commensal rodents is advocated by previous works that have shown that areas with poor housing conditions (including building material quality, sanitation status within and outside the houses) and commercial premises are more prone to rodent infestations than high-standing residential or touristic areas (Adrichem et al., 2013; Byers et al., 2019; Combs et al., 2018a; Jassat et al., 2013; Lambert et al., 2017; Lucaccioni et al., 2016; Masi et al., 2010; Santos et al., 2017). In addition to current features, spatial genetic patterns may also reflect past demographic events related to cityscape evolution during the historical timeframe of urbanization (Harris et al., 2016; Lourenço et al., 2017). This can be especially relevant for non-native invasive commensal rodents that were introduced in numerous primary colonial European settlements, which grew into cities during and after the period of colonial expansion (Aplin et al., 2011; Jones et al., 2013; Puckett et al., 2016).

In this study, we investigate how the interplay between the current and historical features of the cityscape of Dakar, Senegal, may have impacted the spatial pattern of genetic variation of the house mouse, *Mus musculus domesticus*, a major invasive rodent pest recorded in cities of all continents. This subspecies from the *Mus musculus* complex originates from South-West Asia (Suzuki et al., 2013), and became commensal during the initial settlements of humans in the Middle East at circa 10,000 BC (Cucchi et al., 2012). From there, it has expanded its distribution to reach a cosmopolitan range (Bonhomme and Searle, 2012). Several phylogeographic studies have shown that its colonization history is strongly related to human migration patterns at different timescales (Jones et al., 2013). The house mouse may have been introduced in Dakar as soon as it was settled by Europeans in the middle of the nineteenth century (Lippens et al., 2017). The city has then been continuously growing since European settled at the southern end of the Cap-Vert peninsula. From the late nineteenth century to the First World War, urban development was driven by the will to improve commercial links between France and its West African colonies (Harris, 2011). This led to the early creation of a commercial area around the port that further expanded along the railway that connected Dakar to Saint-Louis (the capital city of former French West Africa in the North-West of Senegal) in 1885. The spatial expansion of Dakar has accelerated after World War II and, in absence of sound urban planning, has resulted in a fragmented urban matrix in terms of economic activities and housing, e.g. residential, planned, spontaneous legal and spontaneous illegal housing (Borderon et al., 2014; Diallo et al., 2012). Today, the city is the political, cultural and economic capital of Senegal (Rouhana et al., 2015). It is also one of the leading industrial, financial and service center of West Africa, and a major seaport of the western African coast.

We used two different types of genetic markers to characterize the genetic variation of the house mouse in Dakar: sequences from the mitochondrial DNA control region (D-loop), and 15 nuclear microsatellites. Data on the D-loop, which is the only molecular marker for which substantial data are available over the entire distribution of the taxon (Lippens et al., 2017), were used to further investigate the geographic origin of house mouse populations introduced in Dakar. Microsatellite data, which enable to characterize fine-scale genetic patterns, were used to determine whether spatial genetic structure is rather consistent with an isolation-by-distance process or with well-defined genetic units (in the sense of model-based genetic clustering). We further investigated to which extent considering both the current features of the cityscape and historical information on the urbanization process may help to explain the spatial pattern of genetic variation of the house mouse in Dakar.

## Materials and Methods

### Study area

Dakar is located on the Cap-Vert peninsula on the Atlantic coast (14.6927°N, 17.4466°W) (Fig. 1). It has a tropical climate with a rainy season from June to October, average annual rainfall of ca. 500 mm, and average temperatures between 17–31°C.

**Figure 1.**
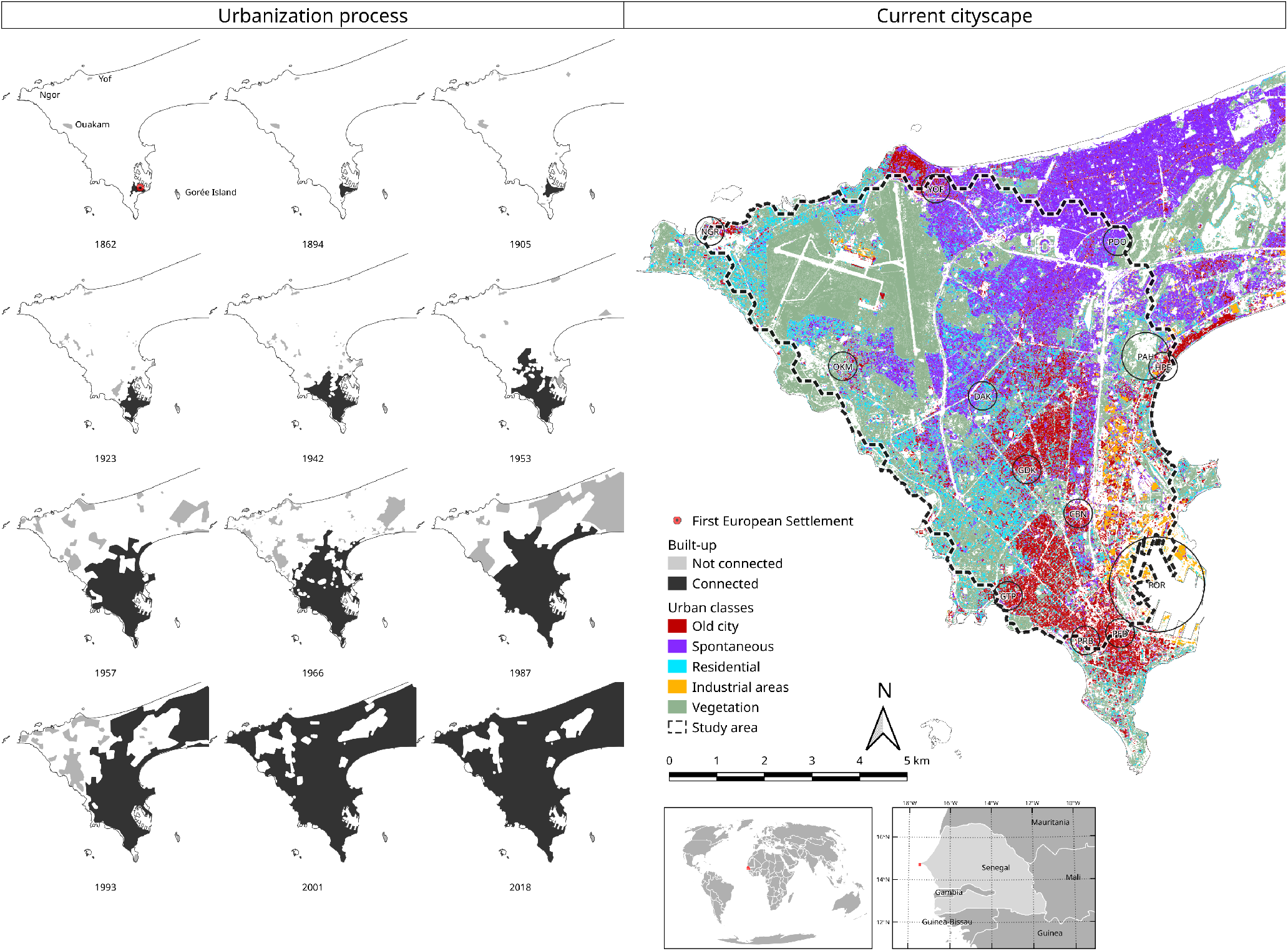
Historical and current Dakar cityscape. Historical cityscape is represented by a time series of 12 maps depicting the spatial expansion of Dakar between 1862 and 2018. The location of the first European settlement is represented with a red dot on the 1862 map while the black and grey polygons represent build-up areas, depending on whether they are connected or not, to the first European Settlement. For each map the coastline corresponds to the current one. The localization of the old fishermen villages of Ouakam, Yoff and Ngor are indicated on the first historical map. Current cityscape of Dakar is based on historical information and results of the spatial PCA performed on the 100 × 100m resolution raster of the typology of socioeconomic habitats retrieved from Borderon et al. (2014) – see Supplementary material, section 1 for details. The study area for the house mouse is delineated with a dashed line. Circles indicates sampling sites (with their code), their diameter indicating the surface area sampled. Small insets representing the location of Senegal and of Dakar were generated using data from Natural Earth (public domain).

The development of the city has gone through several stages (Fig. 1). The first colonial settlement was founded in 1621 on Gorée Island (Sinou, 1993). At this period, the Cap-Vert peninsula was settled by an ethnic group called Lebou, distributed in small villages on the coast. The south cape of the Peninsula (actual Plateau district) has been colonized by Europeans since 1857, but the urban settlement actually began with the achievement of the railway from Saint-Louis to Dakar in 1885 (Sinou, 1993). At the independence in 1960, the city covered only the southern half of the Cap-Vert peninsula (Salem, 1981; Vernière, 1973). It then extended rapidly to include the ancient fishermen villages of Ouakam, Yoff and Ngor located in the North and to eventually cover the whole peninsula. This urban expansion was related to a considerable demographic growth, like in many other cities of the developing countries (Cohen, 2006). For instance, the population of Dakar city increased from 530,000 inhabitants in 1976 (Pélissier, 1983) to 1.15 million in 2013 (ANSD, 2016).

### Spatio-temporal pattern of urbanization

The spatio-temporal dynamics of urbanization in the Cap-Vert peninsula was assessed using: (i) historical data to identify the localization of built-up areas at twelve time points over a 156 year-period, from 1862 to present, and (ii) results of a recent classification of Dakar socioeconomic habitats based on satellite images and census data from Borderon et al. (2014) to describe the current features of the cityscape.

Historical data were gathered from eleven historical maps and one recent map that were retrieved from various web sources or from paper maps provided by the Senegalese national geographical institute. All images were georeferenced as “rasters” using the QGis software (QGIS Development Team, 2018). As coordinates were not always available or reliable, easily identifiable features such as coastlines, major road crossings and outstanding buildings were used as control points for the georeferencing process (see Appendix). Once georeferenced, built-up areas from each map were hand digitized as polygons, tagged as “built-up”, and adjacent polygons were merged. We used the city map of 1862 (Fig. 1) to determine the localization of the first European settlement (spatial coordinates: 14.67210°N; 17.42923°W). For each of the 12 maps, the “built-up” polygons that overlapped the coordinates of the first European settlement were also tagged as “connected”. The tags “built-up” and “connected” attributed to the polygons were then used to define two historical variables that can be computed within any kind of spatial unit: (e.g. raster or grid cells, spatial buffers). The first variable informs on the date at which a spatial unit can be considered as urbanized. This date was determined by considering that “built-up” polygons from a given historical map must cover at least 5% of the surface of the spatial unit at hand. The second variable provides temporal information on the spatial expansion of the city from the first European settlement. Based on the historical maps, a date of connection was attributed to each spatial unit by considering that at least 5% of the surface has to be covered by polygons tagged as “connected”. We then computed the time period elapsed between the year at which most of the mice were sampled (2016) and both the year at which a spatial unit was considered as built-up and as connected to the first European Settlement. These time periods were rescaled between 0 and 1 for further analyses and thereafter referred to as the historical variables “Built-up” and “Connection”, respectively.

To describe the current features of the cityscape within the peninsula, we used six socio-economic classes defined by correlating the habitat typology proposed by the United Nations (UN-Habitat, 2008) and the results of an ascending hierarchical classification, applied on census data of population wealth, that was related to the urban morphology of Dakar based on classification analyses of SPOT 5 images recorded in September 2007 (see Borderon et al., 2014, for details). These classes, presented in Table 1, distinguish: 1) Regular planned housing of good quality (apartments and high-rise houses) mainly located in the oldest part of the city and benefiting from infrastructures such as water supply networks, sewers, waste removal; 2) the scattered village-like habitat with a high density of small low-rise houses with no clear delineations; 3) the regular and 4) irregular spontaneous habitats, which correspond to different waves of suburban development between the sixties and the nineties, 5) the more recent development of high-standing residential areas and 6) areas mainly devoted to commercial and industrial activities that have been expanding from the harbor since the first European settlement. As highly vegetated areas may constitute a very unfavorable habitat (desert resource, Combs et al., 2018b) for the house mouse, which is strictly synanthropic in Senegal (Dalecky et al., 2015), we also extracted information on vegetation occurrence to describe the cityscape of Dakar (Supplementary material, section 1, Fig. S1.1). The proportion of each land cover class was retrieved for each pixel of a 100×100m resolution raster. Due to the successive urbanization policies of Dakar since the colonial period, some features of the current cityscape are spatially highly imbricated, i.e. they strongly covary in space (see section 1 in Supplementary material for historical information on the development of the city). To account for such spatial dependencies between land cover variables, we used a multivariate spatial analysis based on Morans index and implemented in the R (R Core Team, 2019) package “adespatial” (Dray et al., 2017). This approach allows introducing a spatial constraint on a Principal Component Analysis (PCA) through the definition of a spatial weighting matrix, i.e. a connection network, used to compute Moran’s spatial autocorrelation statistic (MULTISPATI-PCA; Dray et al., 2008). MULTISPATI aims at maximizing the product between the variance and the spatial autocorrelation of the scores rather than just the variance as for classical PCA. Here, we used a minimum spanning tree to define the connection matrix between the 100 × 100m pixels. The significance of the observed structure was tested using a Monte Carlo test (10,000 permutations). Based on both the results of the spatial PCA and historical information, we merged some of the land cover classes (see Results section).

**Table 1.** Characteristics of the land cover classes describing Dakar current cityscape as determined by Borderon et al. (2014) using satellite images, UN habitat typology and census data on population wealth. Merging of the initial land cover classes based on the spatial multivariate analysis and historical information is presented in the column “Final land cover classes”

| Initial land cover classes | Characteristics | Final land cover classes |
| --- | --- | --- |
| Old city blocks | Regular planned housing with concrete roads and no sand. Good equipment (water supply networks, sewers) and good accessibility. Mainly apartments and high-rise houses, old and commercial district.<br>*Socio-economic status: mainly high* | Old city |
| Village-like | Anarchic habitat scattered along the peninsula, characterized by a high density of buildings, small plots, low-rise houses and sandy narrow roads. Typically, traditional fishing villages where inhabitants have always maintained customary law on the land.<br>*Socio-economic status: poor to medium high* |  |
| Regular spontaneous | Approved self-building. Grid area with tarred roads between subdivisions and sand roads between land plots. Predominance of multi-storey, solid houses, often concrete.<br>*Socio-economic status: poor to medium high* | Spontaneous |
| Irregular spontaneous | Unplanned housing, often occurring without property title, resulting from important rural migration and moves from downtown during the rapid city sprawl in the 1970s and 1980s. Vehicle accessibility to these areas, often called "floating neighborhoods", is difficult due to narrow sandy streets and flooding. No equipment.<br>*Socio-economic status: mainly poor* |  |
| High standing residential | Individual two-storey houses, villas with gardens or secured residences. High quality equipment and facilities.<br>*Socio-economic status: mainly medium high* | Residential |
| Industrial areas | Mainly non-residential areas with large warehouses, factories, or official administrative buildings.<br>*Socio-economic status: poor and medium* | Industrial |
| Vegetation | Grass, bush or tree cover mixed or not with various type of bare ground such as concrete or sand. | Vegetation |

### Sampling

Within Dakar, we selected 14 sampling sites separated from each other by a median distance of 5850 m (min: 660 m; max: 15,690 m) (Table 2; Fig. 1). They were chosen to represent different stages of the urban history and different habitats (Table 1 and Supplementary material, section 1). Three sites were located in the first urban areas developed by Europeans in Gorée Island (IDG) or in the extreme South of the Cap-Vert peninsula (PRB and PFD). POR corresponds to the commercial harbor of Dakar. GTP corresponds to a first main extension of the city that was developed in 1914 for the indigenous population during a plague epidemic (M’bokolo, 1982). GDK, CBN, and PDO are within districts that were built since 1940. DAK corresponds to the district of Sacré-Cœur founded in 1960 at the heart of the Cap-Vert peninsula. YOF, OKM, NGR and HPE are inside traditional fisherman villages of Lebou that have been successively absorbed by the urban development after 1960. The sampling site PAH is located within the public and zoological park of Hann. Fieldwork was carried out under the framework agreements established between the Institut de Recherche pour le Développement and the Republic of Senegal, as well as with the Senegalese Head Office of Waters and Forests, and the Ministry of Health and of the Social Action. Handling procedures were performed under our lab agreement for experiments on wild animals (no. D-34-169-1), and follow the official guidelines of the American Society of Mammalogists (Sikes and Gannon, 2011). Trapping campaigns within district were systematically performed with prior explicit agreement from relevant local authorities. Sampling was performed in 2016, except the Sacré Coeur district (DAK), which was sampled in 2014 for a previous study (Lippens et al., 2017), according to a standardized protocol described in Dalecky et al. (2015). Each sampling site was the focus of one live trapping session of 2 to 5 consecutive days, in order to reach the target sample size of a minimum of 20 captured house mice on a median surface of 0.04 km^2^ (min: 0.01, max: 1.51; the maximum value concerned POR). In general, two traps (one wire-meshed trap and one Sherman trap) were set per room or courtyard in buildings corresponding to dwelling houses, boutiques, workshops, offices or warehouses, and whose location was precisely recorded with a GPS device. In POR, trapping was done inside and around 19 offices and warehouses (between 5 and 20 pairs of traps per warehouse). Rodents were brought back to our laboratory, euthanized by cervical dislocation, weighted, sexed and measured the same day. House mouse (easily distinguishable morphologically from other rodent species) was the dominant rodent species in all sampling site. A piece of tissue (hind foot) for each adult mouse was stored in 95% ethanol for molecular analyses.

**Table 2.** Characteristics of the sampling sites of *Mus musculus domesticus* in Dakar, Senegal. For each site, are indicated: the latitude and longitude (decimal degrees), the number of mice genotyped at microsatellite markers (N) and sequenced for the mitochondrial D-loop (*n*_*mt*_). For microsatellite data, the table includes an estimation of the median pairwise kinship coefficients (*ρ*, Loiselle et al., 1995), allelic richness (*a*_*r*_), Nei’s unbiased genetic diversity (*H*_*s*_) and inbreeding coefficient (*F*_*IS*_). For mitochondrial data, the table includes D-loop haplotypes names, according to Lippens et al. (2017) (*H*_*D*_ − *loop*).

| Site | code | latitude | longitude | N | $\rho$ | $a_r$ | $H_s$ | $F_{IS}$ | $n_{mt}$ | $H_D - loop$ |
| --- | --- | --- | --- | --- | --- | --- | --- | --- | --- | --- |
| Hann pêcheur | HPE | 14.724035 | -17.429230 | 47 | 0.043 | $4.60 \pm 1.21$ | $0.64 \pm 0.15$ | 0.16 | 3 | H1 |
| Ouakam | OKM | 14.723088 | -17.490068 | 46 | 0.108 | $4.65 \pm 1.52$ | $0.58 \pm 0.18$ | 0.20 | 4 | H1 |
| Ngor | NGR | 14.748939 | -17.517352 | 31 | 0.129 | $4.42 \pm 1.13$ | $0.57 \pm 0.16$ | 0.23 | 4 | H1 |
| Park of Hann | PAH | 14.726967 | -17.434613 | 19 | -0.021 | $4.67 \pm 1.68$ | $0.64 \pm 0.17$ | 0.14 | — | — |
| Yoff | YOF | 14.757200 | -17.473321 | 31 | 0.054 | $4.86 \pm 1.86$ | $0.60 \pm 0.22$ | 0.23 | 4 | H1 |
| Plateau-Rebeuss | PRB | 14.672396 | -17.442259 | 54 | 0.030 | $5.71 \pm 1.55$ | $0.64 \pm 0.13$ | 0.26 | — | — |
| Plateau-Faidherbe | PFD | 14.674027 | -17.436326 | 39 | 0.032 | $5.52 \pm 1.73$ | $0.66 \pm 0.14$ | 0.11 | 4 | H2/H16 |
| Port of Dakar | POR | 14.683414 | -17.430436 | 29 | 0.001 | $6.11 \pm 2.21$ | $0.67 \pm 0.16$ | 0.16 | 8 | H1/H16 |
| Grand Dakar | GDK | 14.704387 | -17.455356 | 34 | 0.056 | $5.09 \pm 1.44$ | $0.59 \pm 0.18$ | 0.24 | — | — |
| Patte d'Oie | PDO | 14.748332 | -17.437953 | 39 | 0.036 | $5.08 \pm 1.77$ | $0.68 \pm 0.11$ | 0.30 | — | — |
| Colobane | CBN | 14.695814 | -17.444924 | 45 | 0.040 | $4.84 \pm 1.46$ | $0.65 \pm 0.12$ | 0.16 | — | — |
| Gueule Tapée | GTP | 14.680193 | -17.458608 | 43 | 0.046 | $4.86 \pm 1.36$ | $0.61 \pm 0.15$ | 0.30 | 4 | H1/H2/H17 |
| Gorée Island | IDG | 14.667140 | -17.398320 | 72 | 0.201 | $3.16 \pm 0.91$ | $0.44 \pm 0.19$ | 0.26 | 9 | H1/H18 |
| Sacré Cœur 3 (2014) | DAK | 14.717635 | -17.463809 | 28 | 0.077 | $4.77 \pm 1.61$ | $0.58 \pm 0.19$ | 0.31 | 5 | H1/H2 |

### Laboratory analyses

For the 529 mice sampled, genomic DNA was extracted from tissues using the DNeasy Blood & Tissue Kit (QIAGEN) according to the manufacturers protocol.

The 876 bp the complete D-loop sequence was amplified for 40 mice sampled within eight of the sampled sites (from three to nine mice from distant houses per site, Table 2). We used PCR primers and conditions described in Rajabi-Maham et al. (2008). PCR products were sequenced in both directions by Eurofins MWG (Ebersberg, Germany).

Genotyping was performed for the 529 mice sampled using 15 nuclear microsatellites loci (D1Mit291, D2Mit456, D3Mit246, D4Mit17, D4Mit241, D6Mit373, D7Mit176, D9Mit51, D10Mit186, D11Mit236, D14Mit66, D16Mit8, D17Mit101, D18Mit8 and D19-Mit30: available from the MMDBJ database: http://www.shigen.nig.ac.jp/mouse/mmdbj/top.jsp), according to Lippens et al. (2017).

### Mitochondrial sequence analysis

All 40 D-loop sequences (731 bp) from this study were aligned with 119 sequences from Senegal (including five sequences from DAK), published by Lippens et al. (2017), and with 1958 sequences retrieved from GenBank: 1672 whose references were compiled in Lippens et al. (2017) and 168 published in other studies (Castiglia et al., 2005; Förster et al., 2009; Gabriel et al., 2011; García-Rodríguez et al., 2018; Jones et al., 2011, 2012; King, 2016; MacKay et al., 2013; Solano et al., 2013; Storz et al., 2007). Sequences alignment was performed using the Clustal W multiple alignment in Bioedit v7.2.6 software (Hall, 1999). Haplotypes were identified with DNA Sequence Polymorphism v6.10.01 software (Rozas et al., 2017). Those found in Dakar were classified following the haplogroup nomenclature defined by Bonhomme et al. (2010). The haplogroups found in this study have already been found from house mice sampled in Senegal and included in the phylogenetic analysis conducted by Lippens et al. (2017). A phenogram was constructed using median joining network calculations implemented in NETWORK v5.0.0.1. (http://www.fluxus-engineering.com) on haplo-types from Senegal, in order to illustrate their relative frequency and relatedness.

### Microsatellite analysis

#### Genetic diversity and differentiation

Population genetic analyses were done on a dataset comprising 557 house mice (Table 2: 529 mice sampled for this study; 28 mice from DAK genotyped in Lippens et al. (2017) using the same markers). Deviations from Hardy-Weinberg Equilibrium (HWE) and genotypic linkage disequilibrium (LD) were tested by locus and by sampling site using GENEPOP v4 (Rousset, 2008). Corrections for multiple tests were performed using the false discovery rate (FDR) approach (Benjamini and Hochberg, 1995) implemented in the Bioconductors qvalue R package (Storey, 2002). Heterozygote deficiencies are often found in house mouse populations and are classically attributed to their social system (Ihle et al., 2006). We calculated the kinship coefficient (*ρ*, Loiselle et al., 1995) between all pairs of individuals within sampling sites, using SPAGeDI v1.4 (Hardy and Vekemans, 2002), in order to evaluate whether heterozygote deficiencies in the dataset may be due to the oversampling of closely related individuals or to the pool of individuals sampled from different social groups (Parreira and Chikhi, 2015). Genotype data from all localities were used as the dataset of reference for allelic frequencies. We also examined whether null alleles may alternatively explain heterozygote deficiencies, by estimating null allele frequencies for each sampling site at each locus using the software FreeNA (Chapuis and Estoup, 2006).

For each sampling site, genetic diversity was estimated using the expected heterozygosity *H*_*s*_ (Nei, 1987), the inbreeding coefficient (*F*_*IS*_), and the allelic richness (*a*_*r*_) calculated using the rarefaction procedure implemented in FSTAT v. 2.9.3 (Goudet, 2001) for a minimum sample of 19 diploid individuals. Genotypic differentiation among sampling sites was tested using Markov chain methods in GENEPOP (using the FDR approach to correct for multiple tests). Genetic differentiation between sampling sites was summarized by calculating pairwise *F*_*ST*_ estimates (Weir and Cockerham, 1984) using FSTAT.

Under a model of isolation by distance (IBD), genetic distance between populations is expected to increase with geographical distance (Rousset, 1997). IBD was analyzed by regressing pairwise estimates of *F*_*ST*_ /(1 − *F*_*ST*_) against between-sites log-transformed geographical distances. Mantel tests were performed to test the correlation between genetic differentiation and Euclidean geographical distance using GENEPOP (10,000 permutations). We used the model-based clustering approach implemented in STRUCTURE v. 2.3.4 (Pritchard et al., 2000), in order to estimate the potential number of homogeneous genetic groups (K) in the data set. The analyses were performed with a model including admixture and correlated allele frequencies (Falush et al., 2003). We performed 20 independent runs for each K values (from K=1 to 10). Each run included 500,000 burn-in iterations followed by 1,000,000 iterations. The number of genetic groups was inferred by the deltaK method applied to the log probabilities of data (Evanno et al., 2005). We checked that a single mode was obtained in the results of the 20 runs for all K-values explored, using the Greedy Algorithm implemented in CLUMPP v. 1.2.2 (Jakobsson and Rosenberg, 2007). Bar plots were finally generated with DISTRUCT v. 1.1 (Rosenberg, 2004). IBD and STRUCTURE analyses were performed with and without the IDG samples, in order to analyze the spatial pattern of genetic variation apart from the effect of the geographical isolation of the island.

#### Spatial patterns of genetic variation and cityscape heterogeneity

The relationship between the spatial patterns of genetic diversity and differentiation and the spatio-temporal urbanization process within the Cap-Vert peninsula (i.e. Gorée Island, locality IDG, was excluded from the dataset) was analyzed using both a punctual (i.e. population-based genetic estimates) and pairwise (i.e. pairwise genetic estimate) approach. For the punctual approach, we considered three genetic estimates for each sampling site: the allelic richness, the expected heterozygosity and a local-*F*_*ST*_ value. The latter was computed using the program GESTE (Foll and Gaggiotti, 2006) and represents the average level of differentiation of one site from all the others. To relate these genetic estimates to the characteristics of the cityscape in the surrounding of each sampling site we used nested circular buffers, of radius equal to 300m, 600m, 1000m and 1500m, centered on the barycenter of the individual coordinates at each site. The value of each cityscape variable was computed as detailed in the section “Spatio-temporal pattern of urbanization”. In order to account for collinearity between nested variables, values within any buffer “*i*” corresponded to the difference between the values of the variables computed within “*i*” and within the buffer nested within “*i*” (Marston et al., 2014). For each buffer size, we computed pairwise Spearman correlation coefficients between all variables. The relationships between nested cityscape features and punctual genetic diversity and differentiation indices (*a*_*r*_, *H*_*s*_ and local-*F*_*ST*_) were then explored using the Random Forest (RF – Breiman, 2001) algorithm “cforest” implemented in the R packages “party” (Hothorn et al., 2005). This approach allows computing conditional permutation importance values that limit the bias toward the systematic selection of variables that are highly correlated to really significant predictors while being actually independent to the response variable (Strobl et al., 2008, 2009). Analyses were done considering different value (2 to 13) for the “mtry” parameter, which controls the number of variables to possibly split at in each node. For each value of “mtry”, 20 replicates were performed using different seed values. The shape of the relationship between genetic estimates and each cityscape feature were determined from partial dependence plots. We also used the R package “ranger” (Wright, Ziegler, et al., 2017) to estimate variable importance p-values based on unconditional RFs (see Supplementary material, Section 3 for details on RFs analyses).

For the pairwise approach, we used the MAPI method, which has a low sensitivity to IBD and is not based on a predefined population genetic model (Piry et al., 2016, implemented in the R package “mapi”). MAPI is a smoothing-like procedure applied on pairwise genetic measures which are attributed to a network of ellipses linking the sampling points. The method produces geographical maps representing the spatial variation in the average level of genetic differentiation between sampled localities. MAPI was run using *F*_*ST*_ values computed between all pairs of localities to build the spatial genetic network of ellipses. To account for the dispersion of individuals within each of the 13 sampled sites, an error circle radius of 300m was applied to the spatial coordinates of all sites, except for the harbor “POR” (radius = 1000m) and the Hann park “PAH” (radius = 500m). A grid of 243 hexagonal cells with a half-width of 310m was superimposed on the study area. This resolution was determined by setting the “beta” parameter to 0.25 in the MAPI analysis in respect of the Nyquist-Shannon sampling theorem under a situation of random sampling (see Piry et al., 2016, for further details). Each cell received the weighted arithmetic mean of the *F*_*ST*_ values attributed to the ellipses intercepting its geographical extent. The mean was weighted using the inverse of the ellipse areas to limit long distance effects (i.e., IBD). The eccentricity value of the ellipses, which controls the smoothing intensity, was set to the default value of 0.975 (see Piry et al., 2016, for further details on the effects of this parameter). We used the permutation procedure (10,000 permutations), along with the FDR approach as described in Piry et al. (2016), to identify major areas of high genetic continuity and discontinuity. The MAPI grid was then superimposed on the layers describing the historical and current cityscape and the value of each variable within each cell of the grid was computed as detailed in the section “Spatio-temporal pattern of urbanization”. As for the punctual approach, Spearman correlation coefficient was calculated for all variable pairs associated to the cells of the MAPI grid. To identify which cityscape features can better explain variations in MAPI smoothed-*F*_*ST*_ values we used conditional RFs, considering different seed and “mtry” (2 to 5) values, to analyze 100 independent training datasets built by randomly subsampling 50% of the MAPI grid cells. Result from each training dataset was used to predict smoothed-*F*_*ST*_ within the remaining MAPI cells (i.e. not included in the training dataset). Model accuracy was assessed by computing the Pearson correlation coefficient (r) between predicted and observed MAPI smoothed-*F*_*ST*_ as well as the R^2^ and RMSE (root mean square error).

Finally, we used a conditional autoregressive model (CAR) to further analyze the relationships between MAPI smoothed-*F*_*ST*_ values and the cityscape variables computed within each cell. To account for spatial autocorrelation, the error term of the model was assumed to have a conditional intrinsic Gaussian autoregressive (CAR) distribution (Besag et al., 1991) with an inverse-distance neighborhood weight matrix. The model was fitted in a Bayesian framework with an INLA approach (Blangiardo and Cameletti, 2015) using the R-INLA package (Martins et al., 2013) (specification of the model for R-INLA is provided in Supplementary material, Section 3). Bayesian inference resulted in posterior densities for all predictors from which significance was assessed with a threshold of 0.05. To evaluate the fit of the model to the data we computed the Pearson correlation coefficient (r) between predicted and observed values as well as the Conditional Predictive Ordinate (CPO) and Probability Integral Transformation (PIT) values (see Held et al., 2010). We checked for the occurrence of extremely small values of CPO indicating unlikely observations under the fitted model. We used a Kolmogorov-Smirnov test to confirm that PIT values did not significantly depart from a uniform distribution as expected if model assumptions are correct. As numerical problem can occur when computing CPO and PIT, the “inla” function of the R-INLA package provides a measure of the reliability of each value which is scaled between 0 and 1 (values far from 0 indicate that for a given observation, the corresponding predictive measure is not reliable). Finally, we also checked that model residuals were not overly structured in space. MAPI is a smoothing procedure that performs better when the sampling maximizes the spatial coverage of the study area (Piry et al., 2016). To evaluate whether the observed genetic structure can be biased, due to the smoothing of a relatively small network (i.e. clumped sampling of 13 localities in the Cap-Vert peninsula), we applied the approach combining the MAPI analysis and Bayesian CAR model on: 1) individual microsatellites genotypes simulated under a landscape configuration corresponding to a spatial gradient from a favorable to an unfavorable habitat as fully described in Piry et al. (2016), and 2) subsampling of the initial dataset, with a number of sampling sites (N) varying from 6 to 12 (see Supplementary Material, section 4, for details on the simulation and subsampling framework).

## Results

### Spatio-temporal pattern of urbanization

The Monte-Carlo permutation test of the MULTISPATI-PCA performed on the seven initial land cover variables describing the current cityscape was highly significant (p<0.001). The first two axes of the spatial PCA explained about 50% of the variance. The first axis strongly reflected the opposition between the occurrence of highly vegetated areas in the western part of the city and very dense urbanized areas in the eastern part. The second axis showed the strong spatial correlation between the land cover classes “Old city blocks” and “Village-like” on one side and between the classes representing the “Regular spontaneous” and “Irregular spontaneous” habitats on the other side (Supplementary material, Section 1, Figs. S1.1, S1.2 and S1.3, Tables S1.1 and S1.2). Based on the results of both the spatial PCA and historical information (see Supplementary material, section 1), we merged the spatially imbricated land cover classes “Old city blocks” and “Village-like” into a single class called “Old city” and the classes “Regular spontaneous” and “Irregular spontaneous” into a same class called “Spontaneous”. Five variables (“Old city”, “Spontaneous”, “Residential”, “Industrial” and “vegetation”) were then retained to characterize the current cityscape in further analyses (see Table 1 and Fig. 1).

### Mitochondrial sequence analyses

There were nine variable sites defining five haplotypes (mean haplotype diversity: 0.264 ± 0.313; mean nucleotide diversity: 0.003 ± 0.005; Fig. 2; Table 2) among the 45 D-loop sequences of house mice from Dakar (40 sequences from this study, five sequences from Lippens et al., 2017). Two aplotypes corresponded to haplotypes H1 (80% of the individuals) and H2 (9% of individuals) previously found by Lippens et al. (2017) to be major D-loop haplotypes in Senegal: they belonged to the haplogroups HG11 and HG4, respectively (Bonhomme and Searle, 2012). H1 was found in all sampling sites except PFD, and H2 in three sampling sites (DAK, GTP and PFD). Both haplotypes had been reported at relatively high frequencies in different part of the world, notably Western Europe (Supplementary material, section 2, Table S2.1). The three other haplotypes (H16, H17 and H18) were not previously found in Senegal. H17 (2.2% of individuals, only found in GTP) and H18 (2.2% of individuals; only found in IDG) were separated from H1 by only a few mutational steps (3 and 1, respectively) and belonged also to HG11 (Fig. 2A). H16 (6.7% of the individuals) was found in POR and PFD and belonged to haplogroup HG1 that was already found in one individual of Saint-Louis, North Senegal (Lippens et al., 2017), and in some localities from Western Europe and Canaries Islands (Supplementary material, section 2, Table S2.1). Haplotype diversity was greater in sampling sites of the southern part of the Cap-Vert peninsula (Table 2; Fig. 2A).

**Figure 2.**
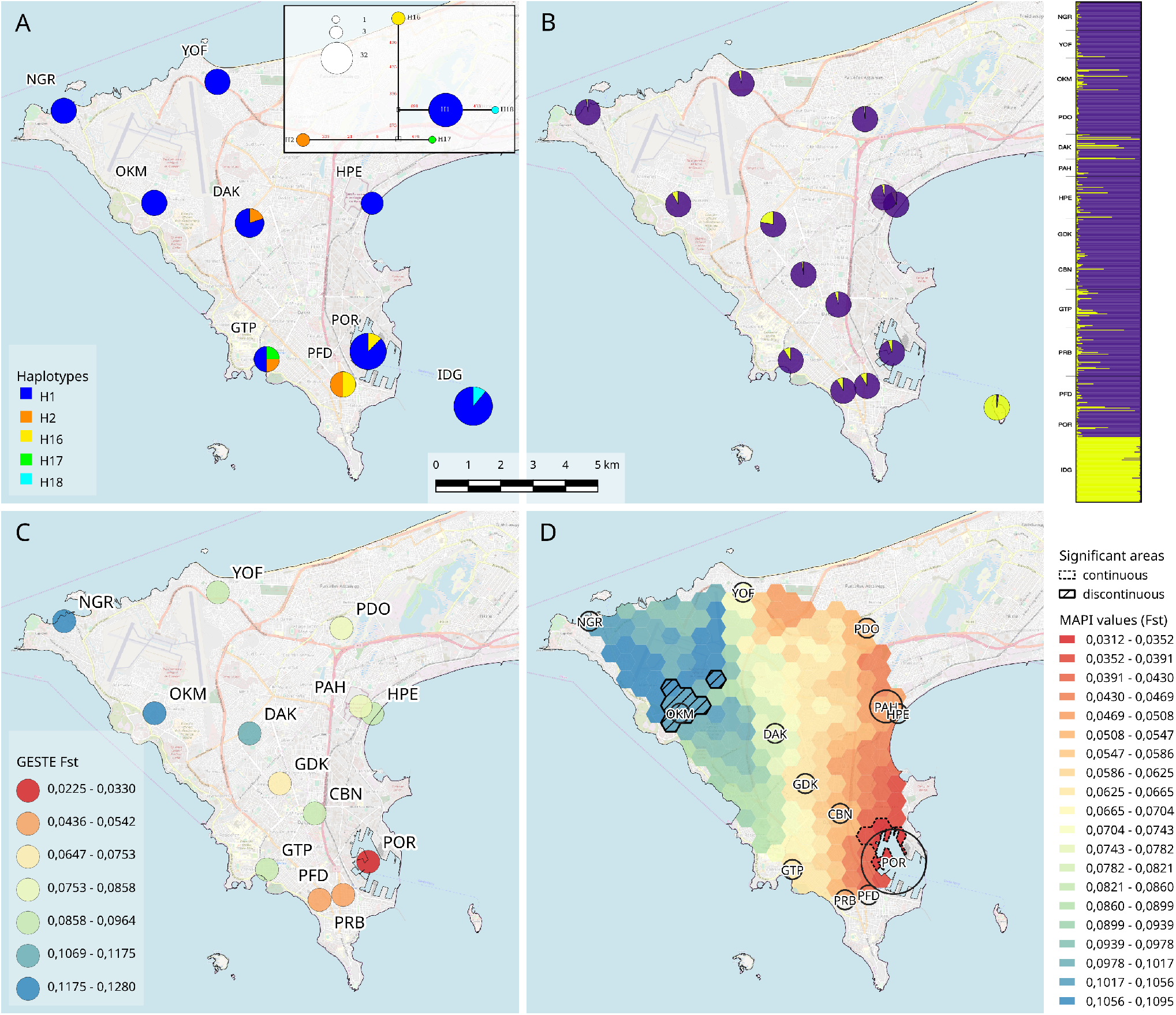
Spatial genetic structure of *Mus musculus domesticus* in Dakar assessed using: **(A)** the five mitochondrial haplotypes found in Dakar The median joining network is presented with white squares corresponding to network nodes, and blue, orange and yellow symbols corresponding to haplotypes from the three haplogroups (HG4, HG1 and HG11, respectively) identified (Bonhomme and Searle, 2012); **(B)** STRUCTURE: results are presented for K = 2 with colors in pie charts indicating the proportion of house mice assigned to each genetic group within each site; the barplot indicates individual ancestry estimates (each vertical line represents an individual) in each genetic group; **(C)** GESTE with a color scale representing the variation in local-*F*_*ST*_ from lowest values in red to highest values in blue; **(D)** MAPI with a color scale representing the spatial variation of the average level of genetic differentiation (*F*_*ST*_), from lowest values in red to highest values in blue; significant areas of high genetic continuity and discontinuity are delineated by dotted and full black lines, respectively. Sampling sites are represented by a circle with a size depending on the spatial distribution of individuals sampled within each site.

### Microsatellite analyses

#### Genetic diversity and differentiation

Linkage disequilibrium was significant for 10 of the 1470 tests performed (0.7%) and the significant values involved different pairs of loci. The 15 loci were thus considered to be genetically independent. None of the loci were at Hardy-Weinberg equilibrium after FDR correction, and positive *F*_*IS*_ values were obtained at all sites (Table 2; Supplementary material, section 2, Table S2.2). The locus D4m17 displayed 11 null genotypes, and its mean *F*_*IS*_ (0.31) was significantly higher than those of the other loci (mean *F*_*IS*_ = 0.22, 95%CI = [0.20;0.24]). This suggested the occurrence of null alleles at this locus. The mean null allele frequency estimated with FreeNA for D4m17 was, however, sufficiently low (0.12) to not significantly affect estimates of genetic diversity and differentiation (Chapuis et al., 2008; Chapuis and Estoup, 2006). Hence, this locus was conserved for further analyses. Null alleles were unlikely to explain heterozygote deficiencies at other loci, since only a small number of null genotypes were observed (0-2 per locus). Within sampling sites, the median kinship Loiselle coefficient *ρ* calculated between all pairs of individuals ranged from −0.021 to 0.201, the higher value being found for IDG (Table 1). Distributions of *ρ* within sites were largely unimodal (Supplementary material, section 2, Fig. S2.1) and globally zero-centered. This suggests that the observed deviations from Hardy-Weinberg equilibrium result from the presence, within each site, of several social groups from which a few individuals have been sampled, rather than a strong family effect due to the oversampling of highly related individuals from only one or a few of these groups (Berthier et al., 2016; Dobson et al., 2007; Parreira and Chikhi, 2015). Allelic richness (*a*_*r*_) ranged from 3.16 to 5.71 (mean 4.88 ± 0.68), *H*_*s*_ from 0.44 to 0.68 (mean 0.61 ± 0.06), and *F*_*IS*_ from 0.11 to 0.31 (mean 0.21 ± 0.01) per sampling site (Table 1). Genotypic differentiation was significant between all sampling sites 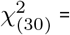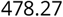 *P* = 0.0001. Pairwise *F*_*ST*_ values ranged from 0.01 to 0.24 with a global mean *F*_*ST*_ value of 0.09 (95% CI = [0.08;0.10]) (Supplementary material, section 2, Table S2.3). As expected, substantial genetic structure was observed between Gorée Island (IDG) and all other sites sampled in the Cap-Vert peninsula (mean *F*_*ST*_ = 0.18 ± 0.03). IBD was significant between sampling sites with IDG (Mantel test: *P* = 0.007; slope *b* = 0.046, 95% CI = [0.034;0.064]) and without IDG (Mantel test: *P* = 0.024; slope *b* = 0.018, 95% CI = [0.011;0.028]).

The program STRUCTURE was used to determine whether house mice were spatially structured as well-defined genetic units in Dakar. The highest deltaK value was obtained for K = 2 (deltaK = 1172 at K = 2, and below 3.8 otherwise). At K = 2, IDG was assigned to a first group, and all individuals from the sampling sites of the Cap-Vert peninsula were largely assigned to a second group (Fig. 2B). At K = 3, the two genetic groups that emerged in the Cap-Vert peninsula were not stable among runs, and many individuals have mixed membership in both groups. Results were similar when IDG was excluded from the analysis (i.e. no clear pattern of spatial structure within the Cap-Vert peninsula).

#### Spatial patterns of genetic variation and cityscape heterogeneity

##### Punctual approach

Posterior mode of local-*F*_*ST*_ estimates, computed for each sampling site with GESTE, ranged from 0.023 (HDPI95% = [0.011; 0.038]) to 0.130 (HDPI95% = [0.097; 0.171]) in POR and OKM, respectively (Supplementary material, Section 3, Table S3.1). The spatial pattern of local genetic differentiation exhibited a gradient-like structure, from the lowest local-*F*_*ST*_ values estimated for the sampling sites located in the South part (POR, PFD, PRB) of the Cap-Vert peninsula to the highest local-*F*_*ST*_ values estimated for the sampling sites located in the North-Western part (OKM, NGR) (Fig. 2C). RF ranking of cityscape features to explain *a*_*r*_, *H*_*s*_ and local-*F*_*ST*_ values was very consistent whatever the setting of the analysis. Moreover, there was no evidence for a bias toward the selection of highly correlated predictor variables during tree building (Supplementary material, section 3, Fig. S3.1 and Table S3.2). Among the seven significant nested variables explaining the variation of *a*_*r*_, four corresponded to the variable “Connection” computed within the different buffers, two to the variable “Vegetation” computed within the 600m- and 1000m-radius buffers and the last one to “Built-up” from the 1000m-radius buffer (Supplementary material, section 3, Fig. S3.1A). The same seven variables were also ranked as the most important to explain the variation in local-*F*_*ST*_ values (Supplementary material, section 3, Fig. S3.1B). As could be expected considering that *a*_*r*_ and local-*F*_*ST*_ were strongly negatively correlated (Supplementary material, section 3, Table S3.2), partial dependence plots showed an opposite effect of “Connection” on *a*_*r*_ and local-*F*_*ST*_, regardless of the buffer size, i.e. increase of *a*_*r*_ and decrease of local-*F*_*ST*_ (Supplementary material, section 3, Fig. S3.2). In the same way, the variable “Vegetation” at 600m and 1000m had a negative and positive effect on *a*_*r*_ and local-*F*_*ST*_, respectively. Finally, “Built-up” at 1000m had an opposite effect, i.e. increase and decrease of *a*_*r*_ and local-*F*_*ST*_, respectively. Although *H*_*s*_ and *a*_*r*_ were positively correlated (Supplementary material, section 3, Table S3.2), a different set of significant variables was identified to explain the variation of *H*_*s*_: “Industrial” computed within the 600m-, 1000m- and 1500m-radius buffers, “Vegetation” from the 1000m- and 1500m-radius buffers and “Residential” from the 1000m-radius buffer (Supplementary material, section 3, Fig. S3.1C). Partial dependence plots suggested a positive effect of “Industrial” and a negative effect of “Vegetation” and “Residential” on *H*_*s*_ (Supplementary material, section 3, Fig. S3.2).

##### Pairwise approach

Similarly to GESTE, MAPI clearly showed a gradient-like pattern from lowest smoothed-*F*_*ST*_ values in the eastern part of the study area to highest values in the western part (Fig. 2D). A significant area of genetic discontinuity encompassed the northern part of the Ouakam district (around the sampling site OKM), between the two main runways of the Léopold-Sédar-Senghor Airport, while a significant area of high genetic continuity was found in the harbor area close to the first European settlement. Spearman correlation tests showed that MAPI smoothed-*F*_*ST*_ values were positively correlated to the variable “Vegetation” and “Residential” (*R*_*s*_ = 0.65 and 0.16, respectively) and negatively correlated to all other variables (*R*_*s*_ varied from −0.32, for the variable “Spontaneous”, to −0.65 for the variable “Connection”; Supplementary material, section 3, Table S3.3). The ranking of the conditional importance of the cityscape features from the RFs trained on 100 randomly resampled datasets (50% of the MAPI grid cells) was very consistent whatever the setting of the analysis: 1) “Connection”, 2) “Vegetation”, 3) “Industrial”, 4) “Spontaneous”, 5) “Residential”, 6) “Old_City” and 7) “Built-up” (Supplementary material, section 3, Fig S3.3 and S3.4). The predictive accuracy of the model was relatively high and slightly varied when the number of variables used for node split increased from 2 to 5. Pearson correlation coefficient (*r*), *R*^2^ and RMSE, averaged over the 100 replicates, ranged: from *r* = 0.81 ± 0.029 to *r* = 0.83 ± 0.030; from *R*^2^ = 0.59 ± 0.034 to *R*^2^ = 0.68 ± 0.046; from RMSE = 0.014 ± 0.001 to RMSE = 0.012 ± 0.001 for “mtry” ranging from 2 to 5.

The reliability measure for the CPO and PIT values of the CAR model was equal to 0 in all cases, indicating that CPO and PIT could be used to assess the calibration of the model. There was no evidence for a lack of fit of the model from the CPO values and the distribution of the PIT values did not significantly depart from a uniform distribution (Kolmogorov-Smirnov test, p-value > 0.05), which confirmed that the model was well calibrated. Pearson correlation coefficient between observed and predicted values was 0.99 and model residuals were not overly structured in space (Supplementary information, section 3, Fig S3.5). Results of the CAR model (Table 3) showed that the posterior probability was significant (at a 5% threshold) for all cityscape variables but “Old_city”. The most important variables to explain the variation in MAPI smoothed-*F*_*ST*_ values were “Connection” and “Industrial”, which both had a negative effect (regression coefficient = −0.081±0.007 and −0.066±0.020, respectively). The two other important variables, “Residential” and “Vegetation” had a positive effect (regression coefficient = 0.059 ± 0.010 and 0.028 ± 0.007, respectively). Lastly, the variables “Builtup” and “Spontaneous” had a slight positive and negative effect, respectively (regression coefficient = 0.017 ± 0.005 and −0.017 ± 0.007).

**Table 3.**
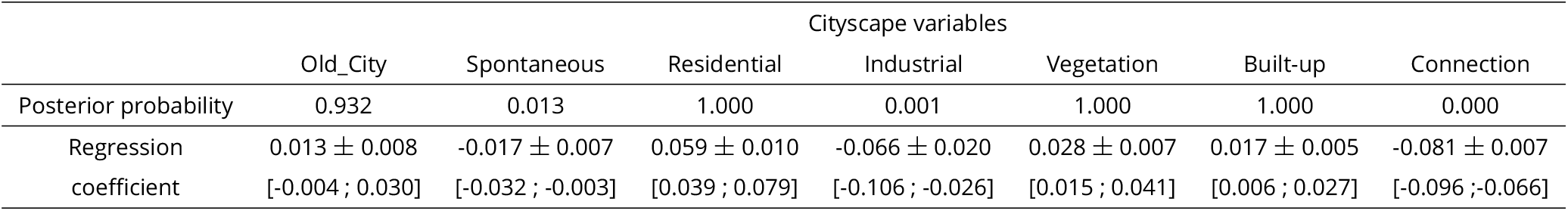
Result of the Bayesian CAR model: posterior probability and regression coefficient associated to each cityscape variable. Considering a threshold of 0.05, a posterior probability > 0.95 indicates a significant positive effect of the corresponding variable while a posterior probability < 0.05 indicates a significant negative effect. Variable regression coefficient is presented with its standard deviation and 95% confidence interval.

##### Simulations

When combining MAPI and a CAR model to analyze 200 datasets of individual genotypes simulated under a landscape configuration corresponding to a spatial gradient of habitat quality, we found the expected significant positive effect of the unfavorable habitat on the MAPI smoothed-*F*_*ST*_ for 95.5% of the simulated datasets. None of the remaining datasets showed an unexpected significant negative effect (Supplementary material, Section 4, Fig S4.1).

##### Subsampling

The analysis of subsampled datasets (with a number of sampling sites N varying from 6 to 12) showed that the gradient-like pattern in smoothed-*F*_*ST*_ and the main effects of cityscape variables were observable even with N=6 sampling sites: high posterior probability for a negative effect of “Connection” and “Industrial” and positive effect of “Vegetation”, “Residential” and “Built-up” (Supplementary material, Section 4, Fig S4.2). For the variables “Old_city” and “Spontaneous”, the number of sampling sites was clearly crucial to obtain consistent posterior probabilities across the replicates.

## Discussion

In this study, we aimed to investigate population genetic structure of *M. m. domesticus* in the city of Dakar, with both mitochondrial sequences and microsatellite markers. By linking genetic patterns with spatial and temporal data describing the urbanization process, we wanted to evaluate whether the genetic structure of this synanthropic species may be reflecting both the heterogeneity of the current cityscape and the historical dynamics of urbanization. We used various approaches, including population genetics model-based clustering, model-free smoothing of genetic differentiation as well as random forests that allow investigating non-linear relationships and taking into account multicollinearity between predictor variables (i.e. conditional permutation importance) and a Bayesian conditional autoregressive model, which allows accounting for spatial autocorrelation. We also performed a simulation study and an analysis of subsampled datasets to assess whether the observed genetic structure could be biased due to the suboptimal spatial coverage of the study area (i.e. clumped sampling of 13 localities in the Cap-Vert peninsula).

Large populations of mice in colonial coastal cities of Senegal have been reported since the middle of the 19^th^ century (Temminck, 1853), which corresponds to the foundation of the first European settlement in the south of the Cap-Vert peninsula. Consistently with these historical inventories, mitochondrial genetic variation observed in house mouse populations in Dakar strongly suggests that the species was introduced during colonial time. Indeed, all haplogroups found in Dakar are typical from Western Europe (Supplementary material, section 2, Table S2.1; Bonhomme and Searle, 2012) with the highest level of diversity observed in sampling sites at the south of the Cap-Vert peninsula (Table 2, Fig. 11). When considering microsatellite markers, beyond the well-marked genetic differentiation of Gorée Island from other sampling sites, there was no evidence that house mice were structured as delineated genetic units in Dakar. Within the Cap-Vert peninsula, the spatial genetic structure corresponded to a gradient-like pattern, from lowest differentiation levels in the south-eastern part of the study area to highest differentiation in the north-western part.

Spatial expansion of introduced populations may be characterized by sequential founder events, leading to spatial decrease of genetic diversity and increase of genetic differentiation along colonization axes (Clegg et al., 2002). For the house mouse within the Cap-Vert peninsula, the analyses of the relationships between genetic estimates (punctual and pairwise) and cityscape variables showed that the duration of the period elapsed since the connection of a built-up area to the first European settlement, was the most important factor to explain the spatial genetic patterns: the shorter this period, the lower the level of allelic richness and the higher the level of genetic differentiation. These results strongly suggest that the observed gradient in genetic variation mainly reflects the historical process of colonization by the house mouse, which followed the continuous expansion of the city from the first European settlement (Gardner-Santana et al., 2009; Harris et al., 2016; Lourenço et al., 2017). This is consistent with the ecology of the house mouse, which is known to be strictly commensal in Senegal, i.e. confined to human buildings (Dalecky et al., 2015). In line with such a strict commensalism, we also found strong negative and positive relationships between the proportion of vegetation and the levels of genetic diversity (*a*_*r*_, *H*_*s*_) and genetic differentiation (local-*F*_*ST*_ and smoothed-*F*_*ST*_), respectively, suggesting that highly vegetated areas (i.e. no or a few buildings used for housing, commercial activities…) act as desert resource habitats and/or dispersal barriers for this species.

Deviations from the main genetic pattern resulting from historical processes can be attributed to the current heterogeneity of the cityscape in terms of habitat quality for the house mouse (Adrichem et al., 2013; Byers et al., 2019; Combs et al., 2018a,b; Lambert et al., 2017; Lucaccioni et al., 2016; Masi et al., 2010; Santos et al., 2017). Spatial variations in the level of genetic differentiation are often interpreted as reflecting the presence of physical barriers to individual dispersal, while they may actually result from variations in effective population size, i.e. demographic barrier (Berthier et al., 2005; Piry et al., 2016; Richardson et al., 2016). Such a demographic process can be especially relevant for commensal rodents as natural dispersal distances (i.e. not human-mediated) of these species are generally short ranged (< 1 km) and mainly prompted by the lack of feeding and harborage sites (Byers et al., 2019; Pocock et al., 2005). In a very large and spatially continuous population, gene flow will occur through step-by-step migration over generations (IBD). Conversely, for an equivalent migration rate in a spatially discontinuous population, important gaps in densities can generate genetic discontinuities (Berthier et al., 2006, 2005). The first situation could correspond to house mouse dynamics in industrial areas that were characterized by high levels of genetic diversity (*H*_*s*_) and low levels of differentiation. In these areas, the presence of numerous warehouses to stock different food products such as groundnuts, rice, sugar and flour (Ba et al., 2013) would allow to sustain large and continuous population of mice. The second situation could explain why high-standing residential areas, which are already known to be less prone to commensal rodent infestations (Adrichem et al., 2013; Byers et al., 2019; Combs et al., 2018a,b; Lambert et al., 2017; Masi et al., 2010), are characterized by low levels of genetic diversity (*H*_*s*_) and high levels of differentiation.

Interestingly, local variations in the two estimates of genetic diversity (*a*_*r*_, *H*_*s*_) are reflecting different processes. Variations of *a*_*r*_ mostly reflect the historical process of colonization (i.e. “Connection”) while variations of *H*_*s*_ are more linked to the current features of the cityscape (i.e. “Industrial” and “Residential”). Changes in *a*_*r*_ depends on the presence/absence of the different alleles sampled through the whole metapopulation while *H*_*s*_ is more dependent on allele encounter probability (mating between individuals carrying different alleles) and, then on the evenness of common allele frequencies (Nei et al., 1975). Historical isolation and consecutive founder events during the expansion process probably had a strong impact on *a*_*r*_ due to the rapid loss of rare alleles that already seldom contributed to *H*_*s*_, which is more dependent on current variations in local density of house mouse due to difference in habitat quality (Comps et al., 2001; Greenbaum et al., 2014).

Two of the cityscape features had a less straightforward effect on the pattern of genetic differentiation (i.e. smoothed-*F*_*ST*_) of the house mouse in the Cap-Vert peninsula: the urban habitat “Spontaneous” and the historical variable “Built-up”. In our study area, the “Spontaneous” habitat mainly corresponds to unplanned housing areas (i.e. irregular spontaneous habitat, see Table 1). These areas often lack basic urban infrastructures (e.g. waste removal) and houses are built with precarious material (e.g. metal sheet, wood, mud brick), which are known to be highly favorable to rodents as they can easily circulate, burrow and reach food storages (Bonwitt et al., 2017; Jassat et al., 2013; Santos et al., 2017). High density of mice in the “Spontaneous” habitat can explain why it is characterized by a low level of genetic differentiation as retrieved by both the RF and CAR models. However, as suggested by the resampling analysis, this effect is highly localized in space and detecting a clear trend strongly depends on the sampling scheme (Supplementary material, Section 4, Fig. S4.2). Local effects, due to the presence of the Lebou villages Ngor, Yoff and Ouakam (NGR, YOF, OKM), can also explain the slight but significant positive effect of the variable “Built-up” in the CAR model. These villages were clearly identified as built-up areas in early maps while they have been among the last areas to be connected to the first European settlement and then, still associated to high levels of differentiation (Figs. 1 and 2). Even if house mice may have been introduced in these villages quite early, they have probably remained isolated until the early 2000s.

Sampling rodents in urban environments may be a challenging task, notably because it requires negotiating access to premises from private owners or city agencies (Parsons et al., 2017). For such a reason, sampling size and coverage may be limited which make it difficult to precisely analyse the relationships between genetic patterns and urban features. In this study, despite the suboptimal coverage of the sampling within the Cap-Vert peninsula, the analysis of simulated and subsampled datasets gave confidence that the MAPI method allowed retrieving the main spatial genetic structure of the house mouse and to relate it to important historical and current features of the cityscape. However, we do believe that, in this particular case, the approach was facilitated by the strength and simplicity of the main structure (i.e. a gradient-like pattern with a high level of genetic differentiation). More complicated or fine-scale genetic structure would be hard to detect without further increasing the sampling size and spatial coverage through a more intensive individual-based sampling design within and across urban habitats (Combs et al., 2018b; Richardson et al., 2017).

In term of rodent management, our results suggest that defining spatial units for house mouse control might not be straightforward in Dakar and, as already pointed out for rats in other cities, classical scales such as individual property or city block are likely to be irrelevant in many local contexts (Gardner-Santana et al., 2009; Kajdacsi et al., 2013). Relevant spatial scale for management will vary across the city due to habitat quality heterogeneity. For example, areas such as the harbor and the large contiguous industrial/commercial area running north are likely to sustain high densities of mice and should be considered as a highly connected network when implementing control strategies (Byers et al., 2019). Areas where mice are likely to be more scattered (residential areas and highly vegetated environment) could constitute spatial targets for localized control as the probability of recolonization through natural dispersal could be relatively low following treatments.

## Conclusion

In this study, we showed that the spatial genetic structure of an invasive and strictly commensal species such as the house mouse likely result from the interplay between the spatio-temporal process of urbanization and the current heterogeneity of the cityscape. First, spatial genetic patterns are consistent with an introduction of the house mouse in the Cap-Vert peninsula during colonial time. Second, the genetic structure within the peninsula mainly corresponds to a gradient-like pattern matching the historical process of spatially continuous expansion of the city from the first European settlement. Deviations from this average pattern reflect the spatial heterogeneity of the current cityscape features in terms of habitat quality for the house mouse, with the residential and highly vegetated areas being the less favorable habitats while the industrial areas and, to some extent, the unplanned housing would be highly favorable to mouse infestation, as already observed for rats. This study highlights the need to investigate for potential effects of past demographic events when conducting genetic studies on non-native invasive commensal rodents in order to assess gene flow to further inform control actions in highly urbanized environment.

## Data accessibility

Data and R pipeline are available online from INRAE repository: https://data.inrae.fr/dataset.xhtml?persistentId=doi:10.15454/NBNSHH

## Supplementary material

Supplementary material and appendix are available from bioRxiv: https://www.biorxiv.org/content/10.1101/557066v3.supplementary-material

## Acknowledgements

We thank all the people in Senegal who welcomed us into their homes for the purposes of small mammal trapping, as well as the Senegalese Head Office of Waters and Forests, the Ministry of Health and Social Action in Senegal, the National hygiene service, and the management director of the Dakar harbour for allowing us to do fieldwork and to export samples. Samples are preserved in the CBGP-Small mammal collection (DOI: 10.15454/WWNUPO) and referenced in the Small Mammal Database associated. Molecular data were generated at the molecular biology platform of CBGP and through the GenSeq technical facilities, with the support of the LabEx CeMEB. We are grateful to Nathalie Sarr for data capture, Caroline Tatard for helping with molecular laboratory work, Julien Papaïx and Thomas Opitz for helpful comments on the Bayesian CAR model, Jean-Marc Duplantier for fruitful discussions on urbanization history and rodent colonization of Dakar. We thank Tuomas Aivelo, Torsti Schulz and an anonymous reviewer as well as the recommender, Michelle DiLeo, for their constructive comments on this work. Funding was provided by the Institut de Recherche pour le Developpement (IRD).

Version 4 of this preprint has been peer-reviewed and recommended by Peer Community In Ecology (Recommendation DOI: 10.24072/pci.ecology.100044).

