## Supplementary material for "Interplay between historical and current features of the cityscape in shaping the genetic structure of the house mouse (*Mus musculus domesticus*) in Dakar (Senegal, West Africa)"

Claire Stragier[1], Sylvain Piry[2], Anne Loiseau[2], Mamadou Kane[1], Aliou Sow[1], Youssoupha Niang[1], Mamoudou Diallo[1], Arame Ndiaye[1], Philippe Gauthier[3], Marion Borderon[4], Laurent Granjon[3], Carine Brouat*[3], and Karine Berthier*[5]

[1] *BIOPASS (IRD-CBGP, ISRA, UCAD), Campus de Bel-Air, BP 1386, CP 18524 Dakar, Senegal*
[2] *CBGP, INRAE, CIRAD, IRD, Montpellier SupAgro, Univ. Montpellier, Montpellier, France*
[3] *CBGP, IRD, CIRAD, INRAE, Montpellier SupAgro, Univ. Montpellier, Montpellier, France*
[4] *Department of Geography and Regional Research, University of Vienna, Austria*
[5] *Pathologie Végétale, INRAE, 84140 Montfavet, France*

**References** **27**

*These two authors contributed equally to this work

### 1 Analysis of historical and current features of Dakar cityscape

#### Historical information on urbanization policies of Dakar

During the colonial period, the French urbanization policies for Dakar were to eliminate traditional, precarious housing from the core of the colonial city. This led to the first wave of expansion of Dakar due to important displacements of indigenous people in the northern periphery of the colonial city. These displacements culminated in 1914 during a major plague epidemic when the French administration stated that rats were highly abundant in traditional housing. The spatial segregation was, however, never really achieved as, first, lot of Lebou people refused to move from the center of the city and, second, the new dwellings where rapidly absorbed with the growing of the city along with its commercial and economic development. The continuous population growth due notably to the high level of immigration from rural areas and neighborhood countries resulted in the densification of the poor dwellings located within more regular and planned city blocks in the oldest central areas of Dakar (e.g. area between GDK, CBN and GTP; see figure S1.2, on page 4).

In the land cover typology retrieved from Borderon et al., 2014, the classes "Old city blocks" and "Village-like" corresponded to the two oldest identifiable types of urban habitats (Table 1). At the end of the 1950s, spontaneous urban areas started to emerge, especially between YOF and PDO. In the 60-70, these spontaneous urban habitats expanded drastically without any kind of urban planning due to the still ongoing expropriations from the center of the city. In the 90s they slightly grew again with the addition of blocks of approved self-buildings. These types of urban habitat correspond to the classes "Irregular and Regular spontaneous habitats" in our land cover typology. The two other classes of the land cover typology, "Residential" and "Industrial", correspond, respectively, to the more recent development of high-standing residential housing conjointly with the increase of international migrations and to the areas with a high density of commercial and industrial premises, which have been expanding from the harbor along the railway since the colonial period.

Sources: Ndiaye, 2015 ; Sinou, 1985 ; Vernière and Gondard, 1974 ; Vernière, 1977.

#### Figure S1.1 - Maps of the current cityscape of Dakar retrieved from Borderon et al. (2014)

Current cityscape of Dakar mapped as a 100x100m resolution raster using the typology of socioeconomic habitats retrieved from Borderon et al., 2014. See Table 1 for details on these classes. The large vegetated area in the north-east of the peninsula corresponds to the Léopold-Sédar-Senghor airport. The study area for the house mouse is delineated with a dashed line and circles represent the spatial dispersion of sampled mice within each study site (see Table 2 for details).

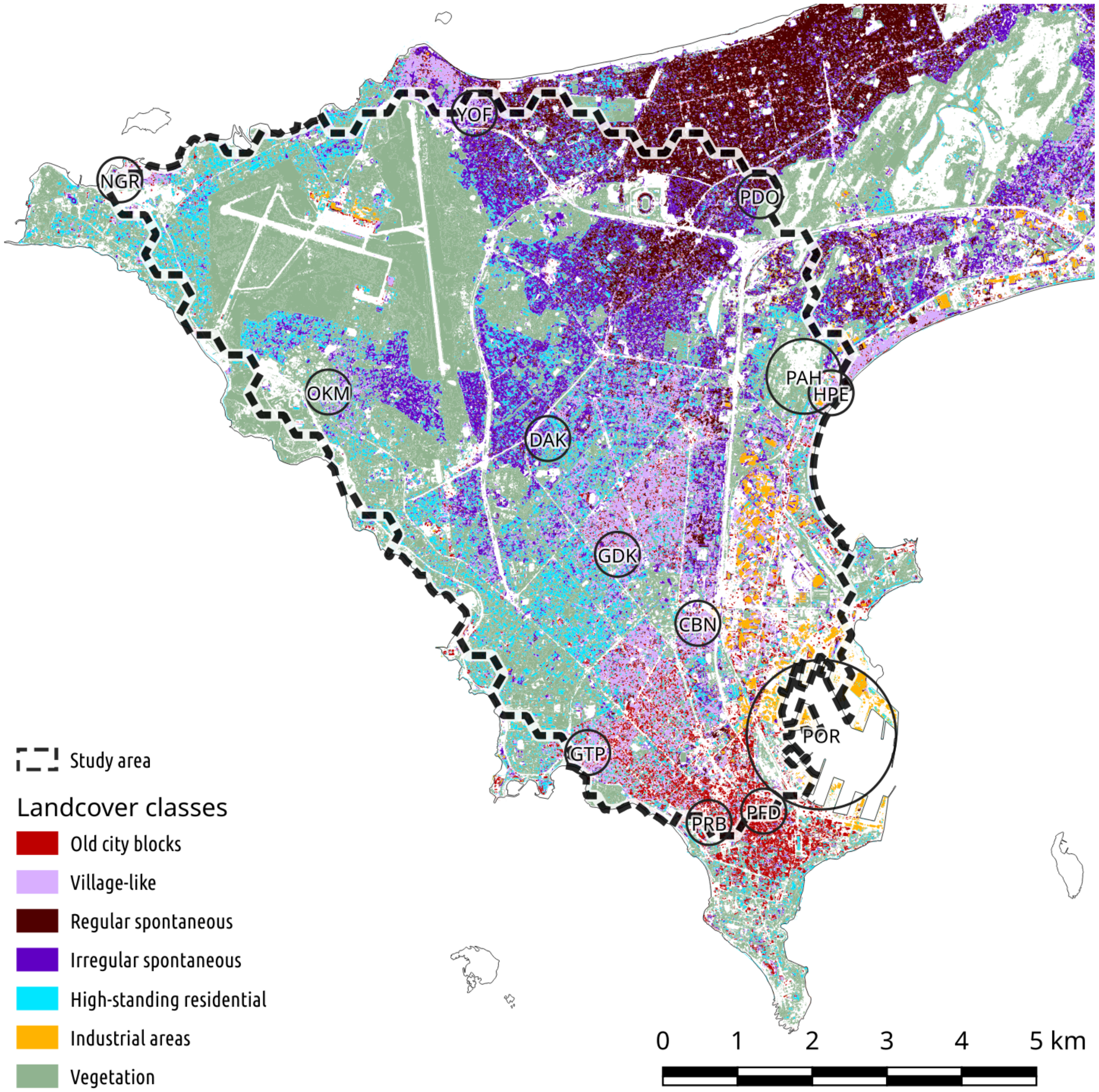

#### Figure S1.2 - Correlation circle of the spatial PCA on current cityscape variables

Correlation circle from the two first axes of the spatial PCA (MULTISPATI; Dray et al., 2008) performed on the seven land cover variables describing the current cityscape of Dakar.

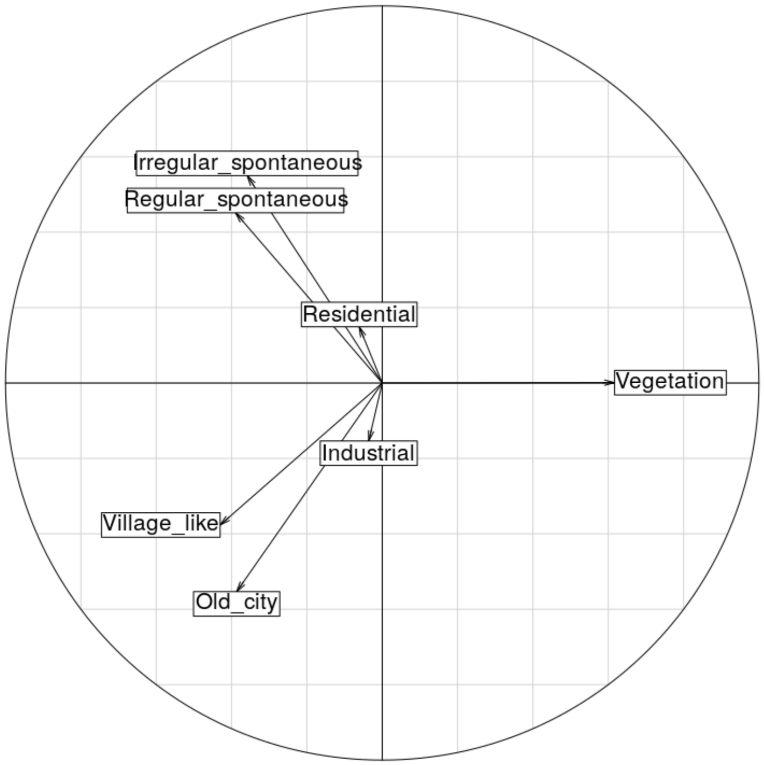

#### Tables S1.1 & S1.2 - Characteristics of the axes of the spatial PCA

**Table S1.1 -** Eigenvalue, variance and Moran spatial autocorrelation coefficient for the first and second axis of the spatial PCA (results of the non-spatial PCA are presented for comparison). The gain in spatial information (Moran) on the two axes was relatively small compared to the classical PCA, which supports that land cover variables are strongly structured in space.

| Axis | PCA | | | | MULTISPATI | | |
|---|---|---|---|---|---|---|---|
| | Variance | Cumulative | Ratio | Moran | Eigenvalue | Variance | Moran |
| 1 | 2.019 | 2.019 | 0.29 | 0.838 | 1.695 | 2.017 | 0.840 |
| 2 | 1.451 | 3.470 | 0.49 | 0.839 | 1.228 | 1.442 | 0.852 |

**Table S1.2 -** Contribution of each variable to the first and second axis of the spatial PCA.

| Variable | Axis 1 | Axis 2 |
|---|---|---|
| Regular spontaneous | -0.389 | 0.451 |
| Old city blocks | -0.386 | -0.553 |
| Irregular spontaneous | -0.358 | 0.550 |
| Residential | -0.061 | 0.149 |
| Industrial | -0.036 | -0.154 |
| Village-like | -0.430 | -0.377 |
| Vegetation | 0.617 | 0.001 |

#### Figure S1.3 - Maps of the scores and lagged scores of the spatial PCA

Mapping of MULTISPATI scores (left) and lagged scores (right) of the 100x100m cells of the raster describing the current cityscape on the first (top) and second axis (bottom). Lagged scores are the averages of the neighboring values weighted by the spatial connection matrix; they provide information on local spatial similarity that can be used to spatially classify the sites based on their characteristics (Dray et al., 2008).

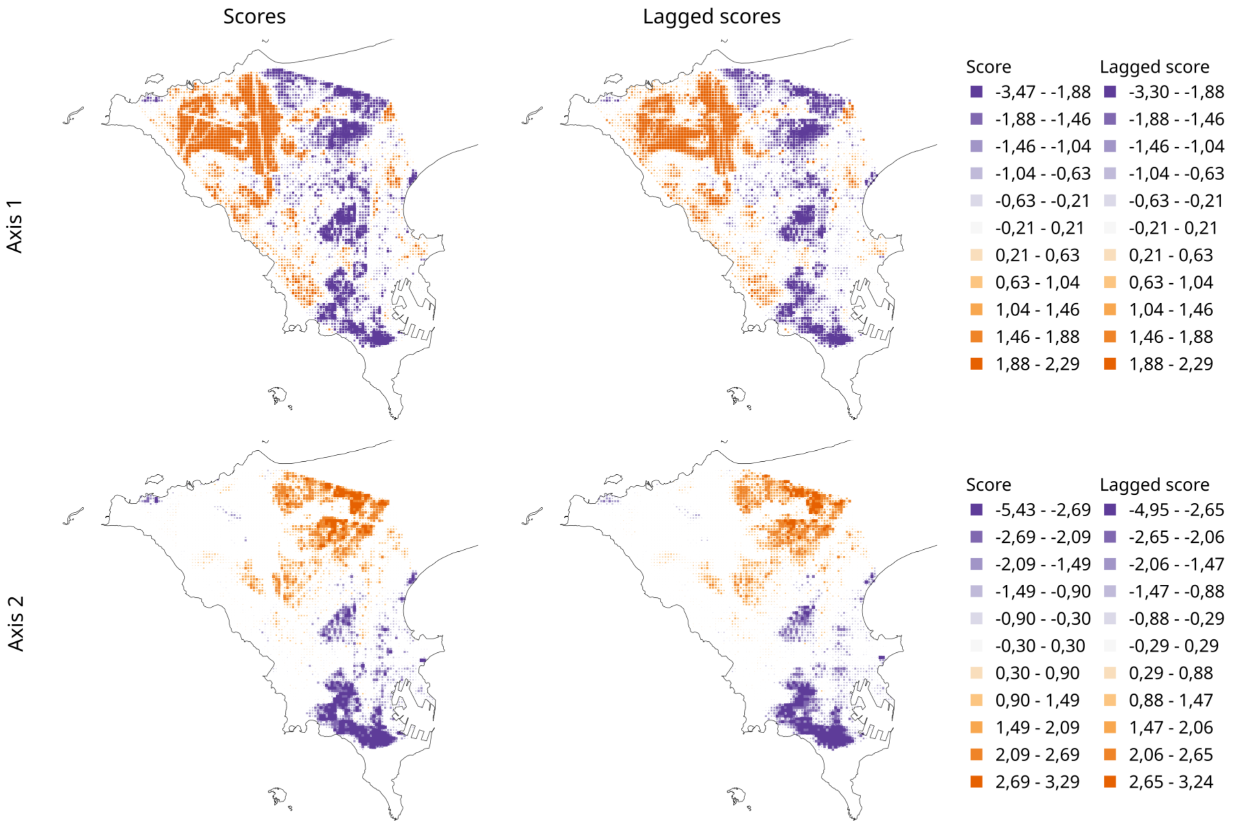

### 2 Mitochondrial and microsatellite supplementary data

#### Table S2.1 - Potential origin of mitochondrial haplotypes

Major locations from where the D-loop haplotypes of house mice sampled in Dakar have been reported. Most of these sequences were compiled in Lippens et al., 2017, and 168 sequences were obtained from other studies (Castiglia et al., 2005; Storz et al., 2007; Förster et al., 2009; EP Jones, Jóhannesdóttir, et al., 2011; Gabriel et al., 2011; EP Jones, Skirnisson, et al., 2012; MacKay et al., 2013; Solano et al., 2013; King, 2016; García-Rodríguez et al., 2018). The percentage of sequences was given for each haplotype, as well as the total number (total nb) of sequences that are in the dataset for each location. Haplotypes H17 and H18 had not been found outside Dakar.

| Location | H1 (%) | H2 (%) | H16 (%) | Total nb |
|---|---|---|---|---|
| Australia | 15 | 15 | | 13 |
| Cameroon | 15 | | | 27 |
| Canaries Islands | | | 32 | 140 |
| France | 24 | 2 | 16 | 164 |
| Germany | 15 | | 13 | 165 |
| Great Britain | 23 | 24 | | 139 |
| Greece | 19 | | | 16 |
| Ireland | | 23 | | 13 |
| Kenya | 0 | 9 | | 11 |
| Lebanon | 2 | | | 59 |
| Madeira | | 2 | | 158 |
| Morocco | 37 | | | 35 |
| New Zealand | 30 | 15 | | 20 |
| Norway | 8 | 92 | | 12 |
| Portugal | 1 | | 3 | 92 |
| Spain | | | 8 | 25 |

#### Table S2.2 - Population polymorphism at the 15 microsatellite loci

Population polymorphism at 15 microsatellite loci over the 557 *Mus musculus domesticus* sampled in 14 sites in Dakar. See Table 2 for codes of sampling sites. $N$ is the number of individuals genotyped per population, $n$ the number of alleles, $a_r$ the allelic richness and $H_s$ the expected heterozygosity. * indicates loci deviating from Hardy-Weinberg expectations (after FDR correction for multiple comparisons, $p < 0.05$)

| | | HPE | OKM | NGR | PAH | YOF | PRB | PFD | POR | GDK | PDO | CBN | GTP | IDG | DAK |
|---|---|---|---|---|---|---|---|---|---|---|---|---|---|---|---|
| | $N$ | 47 | 46 | 31 | 19 | 31 | 54 | 39 | 29 | 34 | 39 | 45 | 43 | 72 | 28 |
| D1Mit291 * | $n$ | 7 | 9 | 6 | 5 | 6 | 10 | 8 | 8 | 7 | 6 | 5 | 7 | 4 | 7 |
| | $a_r$ | 5.8 | 7.5 | 5.1 | 5.0 | 5.6 | 6.6 | 7.0 | 7.0 | 6.1 | 5.7 | 4.9 | 6.0 | 3.2 | 6.5 |
| | $H_s$ | 0.74 | 0.84 | 0.53 | 0.77 | 0.72 | 0.66 | 0.77 | 0.82 | 0.75 | 0.77 | 0.68 | 0.73 | 0.35 | 0.74 |
| D2Mit456 * | $n$ | 5 | 5 | 4 | 5 | 6 | 9 | 8 | 5 | 8 | 7 | 7 | 7 | 4 | 8 |
| | $a_r$ | 4.4 | 4.7 | 3.6 | 5.0 | 5.8 | 6.3 | 6.6 | 4.5 | 6.6 | 6.2 | 6.0 | 5.6 | 3.0 | 6.6 |
| | $H_s$ | 0.74 | 0.52 | 0.50 | 0.75 | 0.69 | 0.68 | 0.71 | 0.64 | 0.73 | 0.77 | 0.71 | 0.69 | 0.50 | 0.54 |
| D3Mit246 * | $n$ | 8 | 10 | 8 | 7 | 10 | 11 | 11 | 12 | 8 | 9 | 10 | 10 | 7 | 7 |
| | $a_r$ | 6.2 | 7.2 | 6.6 | 7.0 | 8.9 | 8.5 | 9.6 | 10.8 | 6.9 | 8.3 | 7.9 | 8.1 | 4.8 | 6.3 |
| | $H_s$ | 0.75 | 0.79 | 0.58 | 0.75 | 0.85 | 0.85 | 0.87 | 0.84 | 0.60 | 0.85 | 0.79 | 0.81 | 0.67 | 0.73 |
| D4Mit17 * | $n$ | 3 | 3 | 3 | 4 | 4 | 7 | 4 | 5 | 4 | 3 | 6 | 5 | 3 | 4 |
| | $a_r$ | 3.0 | 2.7 | 3.0 | 4.0 | 4.0 | 5.8 | 3.9 | 4.3 | 3.9 | 3.0 | 5.9 | 4.4 | 2.8 | 4.0 |
| | $H_s$ | 0.62 | 0.30 | 0.58 | 0.64 | 0.64 | 0.74 | 0.66 | 0.69 | 0.62 | 0.64 | 0.75 | 0.71 | 0.54 | 0.59 |
| D4Mit241 * | $n$ | 6 | 4 | 3 | 3 | 4 | 6 | 6 | 6 | 4 | 3 | 3 | 6 | 4 | 4 |
| | $a_r$ | 5.2 | 3.9 | 2.9 | 3.0 | 3.8 | 5.5 | 4.9 | 5.3 | 4.0 | 3.0 | 3.0 | 5.2 | 2.9 | 3.7 |
| | $H_s$ | 0.67 | 0.60 | 0.24 | 0.64 | 0.60 | 0.56 | 0.67 | 0.68 | 0.73 | 0.64 | 0.62 | 0.67 | 0.23 | 0.66 |
| D6Mit373 * | $n$ | 5 | 5 | 6 | 5 | 5 | 9 | 7 | 10 | 5 | 6 | 5 | 7 | 3 | 5 |
| | $a_r$ | 4.1 | 4.0 | 5.6 | 5.0 | 5.0 | 6.7 | 6.2 | 8.9 | 4.3 | 4.8 | 4.8 | 5.3 | 2.5 | 4.9 |
| | $H_s$ | 0.60 | 0.56 | 0.75 | 0.64 | 0.76 | 0.75 | 0.76 | 0.74 | 0.45 | 0.68 | 0.68 | 0.51 | 0.51 | 0.67 |
| D7Mit176 * | $n$ | 6 | 7 | 5 | 7 | 6 | 7 | 7 | 8 | 8 | 7 | 7 | 6 | 5 | 8 |
| | $a_r$ | 5.9 | 6.0 | 4.9 | 7.0 | 5.9 | 5.9 | 6.6 | 7.6 | 6.7 | 6.7 | 6.6 | 6.0 | 3.6 | 7.2 |
| | $H_s$ | 0.78 | 0.63 | 0.77 | 0.83 | 0.79 | 0.71 | 0.76 | 0.82 | 0.79 | 0.79 | 0.81 | 0.80 | 0.33 | 0.81 |
| D9Mit51 * | $n$ | 5 | 5 | 5 | 8 | 10 | 8 | 8 | 9 | 7 | 9 | 7 | 6 | 3 | 5 |
| | $a_r$ | 4.8 | 4.3 | 4.9 | 8.0 | 7.9 | 6.3 | 6.1 | 8.3 | 6.2 | 6.7 | 5.2 | 5.6 | 2.7 | 4.9 |
| | $H_s$ | 0.69 | 0.66 | 0.59 | 0.66 | 0.76 | 0.54 | 0.64 | 0.77 | 0.69 | 0.72 | 0.57 | 0.68 | 0.41 | 0.75 |
| D10Mit186 * | $n$ | 6 | 5 | 5 | 4 | 3 | 6 | 5 | 5 | 4 | 5 | 5 | 5 | 4 | 5 |
| | $a_r$ | 5.0 | 4.4 | 3.8 | 4.0 | 3.0 | 4.6 | 4.5 | 4.6 | 3.5 | 4.7 | 4.4 | 4.4 | 3.6 | 4.6 |
| | $H_s$ | 0.70 | 0.66 | 0.24 | 0.60 | 0.40 | 0.49 | 0.64 | 0.58 | 0.41 | 0.65 | 0.58 | 0.58 | 0.55 | 0.50 |
| D11Mit236 * | $n$ | 2 | 2 | 3 | 2 | 3 | 2 | 2 | 3 | 3 | 2 | 3 | 2 | 2 | 3 |
| | $a_r$ | 2.0 | 2.0 | 2.6 | 2.0 | 2.6 | 2.0 | 2.0 | 3.0 | 2.8 | 2.0 | 2.9 | 2.0 | 2.0 | 2.4 |
| | $H_s$ | 0.19 | 0.20 | 0.48 | 0.19 | 0.23 | 0.36 | 0.35 | 0.28 | 0.28 | 0.41 | 0.33 | 0.39 | 0.33 | 0.07 |
| D14Mit66 * | $n$ | 5 | 5 | 4 | 5 | 5 | 7 | 6 | 5 | 5 | 5 | 4 | 5 | 2 | 5 |
| | $a_r$ | 4.9 | 4.6 | 3.9 | 5.0 | 4.6 | 6.2 | 4.7 | 4.6 | 4.6 | 4.9 | 3.4 | 4.2 | 1.7 | 4.6 |
| | $H_s$ | 0.74 | 0.70 | 0.60 | 0.66 | 0.41 | 0.60 | 0.51 | 0.50 | 0.34 | 0.69 | 0.57 | 0.61 | 0.05 | 0.55 |
| D16Mit8 * | $n$ | 5 | 5 | 6 | 3 | 4 | 6 | 5 | 7 | 6 | 6 | 4 | 5 | 4 | 3 |
| | $a_r$ | 4.3 | 4.9 | 5.5 | 3.0 | 4.0 | 4.1 | 4.8 | 6.4 | 5.4 | 5.5 | 3.8 | 4.1 | 4.0 | 3.0 |
| | $H_s$ | 0.64 | 0.60 | 0.58 | 0.66 | 0.71 | 0.62 | 0.72 | 0.73 | 0.69 | 0.76 | 0.69 | 0.60 | 0.71 | 0.55 |
| D17Mit101 * | $n$ | 7 | 7 | 5 | 5 | 6 | 9 | 7 | 9 | 8 | 8 | 7 | 6 | 4 | 7 |
| | $a_r$ | 6.2 | 6.1 | 5.0 | 5.0 | 5.4 | 6.8 | 6.5 | 7.8 | 7.3 | 7.2 | 6.0 | 4.7 | 3.6 | 6.7 |
| | $H_s$ | 0.74 | 0.74 | 0.77 | 0.75 | 0.65 | 0.78 | 0.72 | 0.78 | 0.75 | 0.75 | 0.69 | 0.65 | 0.49 | 0.72 |
| D18Mit8 * | $n$ | 4 | 5 | 4 | 3 | 2 | 5 | 7 | 4 | 4 | 4 | 3 | 5 | 3 | 3 |
| | $a_r$ | 3.9 | 4.1 | 4.0 | 3.0 | 1.9 | 3.9 | 4.4 | 3.7 | 3.5 | 3.9 | 3.0 | 4.1 | 2.3 | 3.0 |
| | $H_s$ | 0.47 | 0.42 | 0.61 | 0.32 | 0.06 | 0.49 | 0.42 | 0.45 | 0.33 | 0.58 | 0.58 | 0.29 | 0.20 | 0.37 |
| D19Mit30 * | $n$ | 4 | 4 | 5 | 4 | 5 | 7 | 6 | 5 | 5 | 4 | 5 | 4 | 6 | 3 |
| | $a_r$ | 3.4 | 3.5 | 4.8 | 4.0 | 4.6 | 6.6 | 5.0 | 4.9 | 4.5 | 3.5 | 4.9 | 3.5 | 4.7 | 3.0 |
| | $H_s$ | 0.59 | 0.51 | 0.69 | 0.70 | 0.68 | 0.77 | 0.66 | 0.73 | 0.70 | 0.53 | 0.73 | 0.47 | 0.67 | 0.49 |
| Across all loci | $n$ | 5 | 5 | 5 | 5 | 5 | 7 | 6 | 7 | 6 | 6 | 5 | 6 | 4 | 5 |
| | $a_r$ | 4.6 | 4.7 | 4.4 | 4.7 | 4.9 | 5.7 | 5.5 | 6.1 | 5.1 | 5.1 | 4.8 | 4.9 | 3.2 | 4.8 |
| | $H_s$ | 0.64 | 0.58 | 0.57 | 0.64 | 0.60 | 0.64 | 0.66 | 0.67 | 0.59 | 0.68 | 0.65 | 0.61 | 0.44 | 0.58 |

##### Table S2.3 - Genetic differentiation between localities

Pairwise $F_{ST}$ values between 14 sampling sites of *Mus musculus domesticus* in Dakar, Senegal. See Table 2 for codes of sampling sites.

| | CBN | DAK | GDK | GTP | HPE | IDG | NGR | OKM | PAH | PDO | PFD | POR | PRB |
|---|---|---|---|---|---|---|---|---|---|---|---|---|---|
| DAK | 0.0879 | | | | | | | | | | | | |
| GDK | 0.0577 | 0.0637 | | | | | | | | | | | |
| GTP | 0.0615 | 0.0834 | 0.0547 | | | | | | | | | | |
| HPE | 0.0437 | 0.0687 | 0.0501 | 0.0474 | | | | | | | | | |
| IDG | 0.1595 | 0.2085 | 0.1931 | 0.1800 | 0.1824 | | | | | | | | |
| NGR | 0.0922 | 0.1223 | 0.1078 | 0.1140 | 0.0921 | 0.2468 | | | | | | | |
| OKM | 0.1061 | 0.1159 | 0.1043 | 0.0774 | 0.0859 | 0.1918 | 0.1178 | | | | | | |
| PAH | 0.0465 | 0.0702 | 0.0498 | 0.0631 | 0.0344 | 0.1753 | 0.0903 | 0.0599 | | | | | |
| PDO | 0.0466 | 0.0725 | 0.0524 | 0.0729 | 0.0380 | 0.1928 | 0.1000 | 0.1088 | 0.0493 | | | | |
| PFD | 0.0412 | 0.0660 | 0.0491 | 0.0542 | 0.0312 | 0.1619 | 0.0814 | 0.1035 | 0.0424 | 0.0354 | | | |
| POR | 0.0289 | 0.0413 | 0.0233 | 0.0360 | 0.0264 | 0.1583 | 0.0814 | 0.0740 | 0.0268 | 0.0327 | 0.0155 | | |
| PRB | 0.0394 | 0.0708 | 0.0729 | 0.0705 | 0.0584 | 0.1446 | 0.0771 | 0.1019 | 0.0491 | 0.0524 | 0.0370 | 0.0245 | |
| YOF | 0.0526 | 0.0878 | 0.0423 | 0.0738 | 0.0475 | 0.1816 | 0.0985 | 0.1181 | 0.0323 | 0.0481 | 0.0448 | 0.0376 | 0.0502 |

#### Figure S2.1 - Pairwise kinship coefficient

Distribution of pairwise kinship coefficients (Loiselle et al., 1995) computed between all pairs of house mice from the same site. See Table 2 for codes of sampling sites.

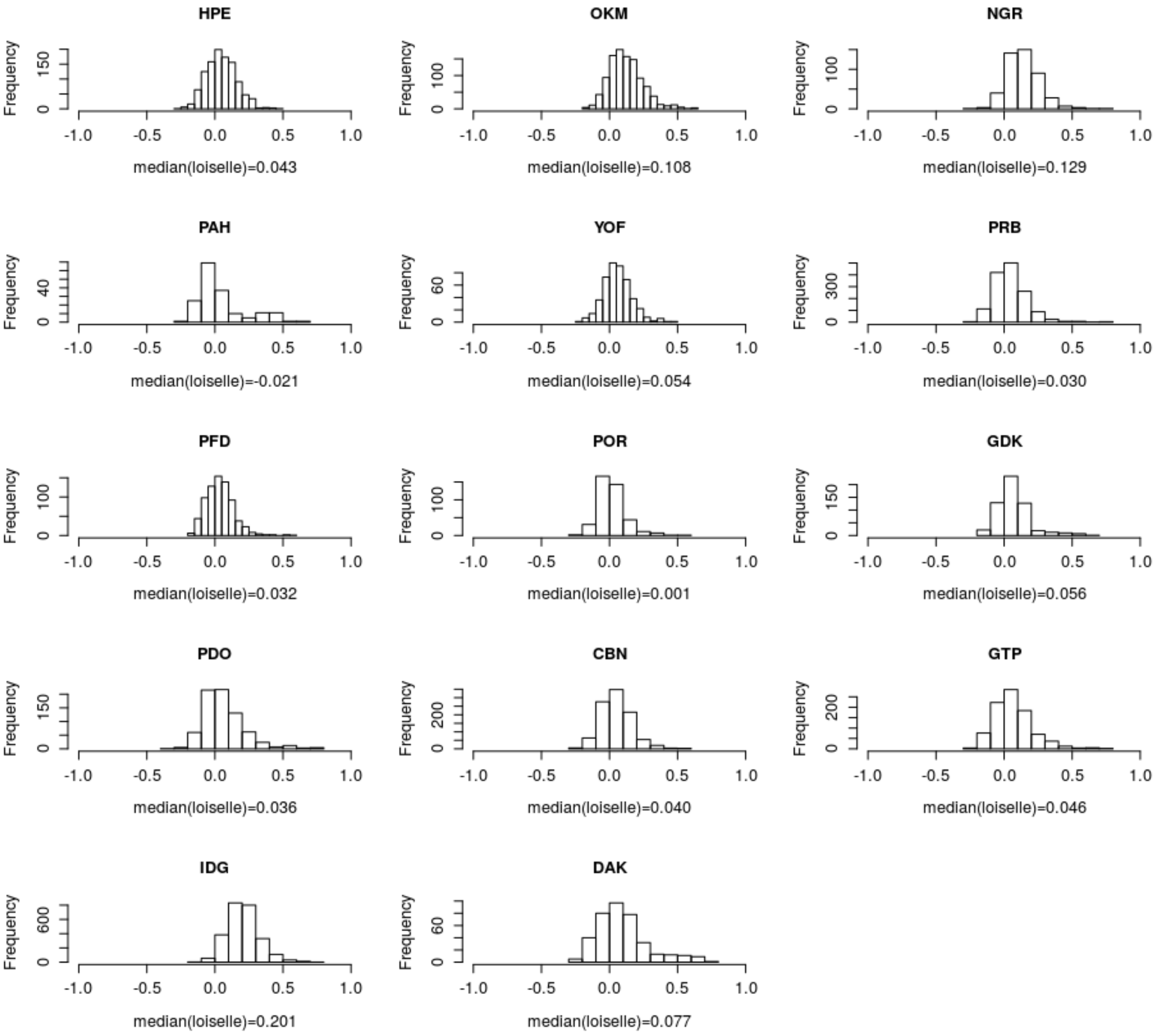

### 3 Spatial patterns of genetic variation and cityscape heterogeneity

#### Table S3.1 - Local - $F_{ST}$ estimates for each locality within the Cap-Vert peninsula

Mode and 95% highest probability density intervals (HPDI) of local - $F_{ST}$ estimates computed with the program GESTE (Foll et al., 2006) for each population located within the Cap-Vert peninsula.

| Locality | Mode local - $F_{ST}$ | 95% HDPI |
|---|---|---|
| CBN | 0.093 | [0.068 ; 0.129] |
| DAK | 0.110 | [0.077 ; 0.149] |
| GDK | 0.071 | [0.047 ; 0.099] |
| GTP | 0.092 | [0.065 ; 0.123] |
| HPE | 0.095 | [0.070 ; 0.134] |
| NGR | 0.124 | [0.090 ; 0.171] |
| OKM | 0.130 | [0.097 ; 0.171] |
| PAH | 0.083 | [0.055 ; 0.126] |
| PDO | 0.076 | [0.054 ; 0.107] |
| PFD | 0.049 | [0.033 ; 0.070] |
| POR | 0.023 | [0.011 ; 0.038] |
| PRB | 0.045 | [0.033 ; 0.062] |
| YOF | 0.088 | [0.062 ; 0.126] |

#### Table S3.2 - Spearman correlations for the punctual approach

Spearman correlation coefficient ($R_s$) and p-values estimated for all pairs of variables computed within the nested buffers. Correlation coefficients between genetic estimates computed within sites are presented in the column corresponding to the 300m buffer size. Significant values for $R_s$ are indicated in bold.

| | | 300m | | 600m | | 1000m | | 1500m | |
|---|---|---|---|---|---|---|---|---|---|
| Variable 1 | Variable 2 | $R_s$ | p-value | $R_s$ | p-value | $R_s$ | p-value | $R_s$ | p-value |
| $a_r$ | $H_s$ | **0.58** | 0.038 | | | | | | |
| $a_r$ | local - $F_{ST}$ | **-0.91** | 0.000 | | | | | | |
| $H_s$ | local - $F_{ST}$ | **-0.67** | 0.011 | | | | | | |
| $a_r$ | Vegetation | **-0.66** | 0.014 | **-0.81** | 0.001 | **-0.76** | 0.002 | -0.40 | 0.177 |
| $H_s$ | Vegetation | -0.29 | 0.341 | **-0.69** | 0.010 | **-0.79** | 0.001 | **-0.89** | 0.000 |
| local - $F_{ST}$ | Vegetation | **0.56** | 0.044 | **0.85** | 0.000 | **0.82** | 0.001 | **0.54** | 0.055 |
| $a_r$ | Spontaneous | -0.18 | 0.565 | -0.22 | 0.464 | -0.10 | 0.734 | -0.16 | 0.596 |
| $H_s$ | Spontaneous | -0.24 | 0.428 | -0.14 | 0.639 | 0.14 | 0.659 | 0.26 | 0.396 |
| local - $F_{ST}$ | Spontaneous | 0.34 | 0.255 | 0.21 | 0.493 | **0.03** | 0.929 | **0.03** | 0.915 |
| $a_r$ | Old_city | 0.29 | 0.338 | 0.59 | 0.035 | **0.67** | 0.013 | **0.58** | 0.036 |
| $H_s$ | Old_city | -0.06 | 0.850 | 0.25 | 0.407 | 0.38 | 0.198 | 0.41 | 0.165 |
| local - $F_{ST}$ | Old_city | -0.16 | 0.590 | **-0.63** | 0.022 | **-0.60** | 0.029 | -0.51 | 0.074 |
| $a_r$ | Residential | -0.39 | 0.190 | -0.54 | 0.057 | -0.50 | 0.079 | -0.04 | 0.908 |
| $H_s$ | Residential | **-0.70** | 0.008 | **-0.77** | 0.002 | **-0.78** | 0.002 | **-0.58** | 0.037 |
| local - $F_{ST}$ | Residential | 0.50 | 0.078 | 0.53 | 0.064 | **0.58** | 0.037 | 0.18 | 0.553 |
| $a_r$ | Industrial | 0.04 | 0.884 | 0.13 | 0.663 | 0.27 | 0.370 | 0.29 | 0.343 |
| $H_s$ | Industrial | 0.52 | 0.070 | **0.72** | 0.005 | **0.82** | 0.001 | **0.77** | 0.002 |
| local - $F_{ST}$ | Industrial | -0.23 | 0.455 | -0.28 | 0.352 | -0.46 | 0.115 | -0.45 | 0.122 |
| $a_r$ | Builtup | 0.20 | 0.514 | 0.29 | 0.341 | **0.67** | 0.013 | **0.56** | 0.048 |
| $H_s$ | Builtup | -0.10 | 0.735 | -0.07 | 0.819 | 0.30 | 0.317 | 0.26 | 0.386 |
| local - $F_{ST}$ | Builtup | -0.08 | 0.780 | -0.15 | 0.616 | **-0.59** | 0.034 | **-0.45** | 0.119 |
| $a_r$ | Connection | **0.70** | 0.008 | **0.72** | 0.006 | **0.72** | 0.006 | **0.70** | 0.007 |
| $H_s$ | Connection | **0.60** | 0.029 | **0.58** | 0.040 | **0.56** | 0.048 | 0.50 | 0.085 |
| local - $F_{ST}$ | Connection | **-0.71** | 0.006 | **-0.70** | 0.008 | **-0.70** | 0.007 | **-0.65** | 0.015 |
| Vegetation | Spontaneous | 0.13 | 0.681 | 0.03 | 0.915 | -0.12 | 0.707 | -0.31 | 0.297 |
| Vegetation | Old_city | **-0.59** | 0.033 | -0.46 | 0.117 | -0.54 | 0.058 | -0.51 | 0.078 |
| Spontaneous | Old_city | -0.03 | 0.915 | -0.28 | 0.354 | -0.34 | 0.255 | -0.20 | 0.517 |
| Vegetation | Residential | 0.54 | 0.058 | **0.69** | 0.010 | **0.65** | 0.017 | 0.41 | 0.162 |
| Spontaneous | Residential | 0.25 | 0.405 | 0.07 | 0.831 | -0.27 | 0.374 | -0.51 | 0.078 |
| Old_city | Residential | 0.07 | 0.817 | -0.03 | 0.929 | -0.13 | 0.681 | 0.30 | 0.316 |
| Vegetation | Industrial | -0.27 | 0.375 | -0.44 | 0.132 | **-0.66** | 0.015 | **-0.81** | 0.001 |
| Spontaneous | Industrial | -0.29 | 0.342 | -0.03 | 0.921 | 0.09 | 0.760 | 0.16 | 0.590 |
| Old_city | Industrial | -0.16 | 0.612 | 0.02 | 0.942 | 0.30 | 0.322 | 0.48 | 0.094 |
| Residential | Industrial | **-0.65** | 0.017 | -0.45 | 0.122 | **-0.57** | 0.042 | -0.42 | 0.156 |
| Vegetation | Builtup | -0.51 | 0.072 | -0.25 | 0.407 | -0.44 | 0.136 | -0.31 | 0.296 |
| Spontaneous | Builtup | -0.27 | 0.381 | **-0.58** | 0.037 | **-0.68** | 0.010 | **-0.59** | 0.035 |
| Old_city | Builtup | 0.27 | 0.366 | 0.17 | 0.590 | **0.82** | 0.001 | **0.80** | 0.001 |
| Residential | Builtup | -0.42 | 0.150 | -0.28 | 0.360 | -0.10 | 0.755 | 0.33 | 0.265 |
| Industrial | Builtup | 0.11 | 0.718 | -0.13 | 0.671 | 0.30 | 0.317 | 0.47 | 0.106 |
| Vegetation | Connection | **-0.58** | 0.038 | **-0.57** | 0.044 | -0.66 | 0.014 | -0.52 | 0.071 |
| Spontaneous | Connection | **-0.60** | 0.029 | **-0.60** | 0.032 | -0.45 | 0.120 | -0.34 | 0.258 |
| Old_city | Connection | 0.48 | 0.099 | **0.73** | 0.004 | **0.83** | 0.000 | **0.87** | 0.000 |
| Residential | Connection | -0.39 | 0.193 | -0.23 | 0.459 | -0.19 | 0.540 | 0.17 | 0.587 |
| Industrial | Connection | 0.34 | 0.250 | 0.37 | 0.217 | **0.55** | 0.049 | **0.62** | 0.023 |
| Builtup | Connection | 0.14 | 0.643 | 0.25 | 0.416 | **0.90** | 0.000 | **0.94** | 0.000 |

#### Setting of random forests on punctual genetic estimates ($a_r$, $H_s$ and local - $F_{ST}$)

We used two different RF algorithms: "cforest" and "ranger" implemented in the R packages "party" (Hothorn et al., 2005) and "ranger" (Wright et al., 2017), respectively. Both approaches allow computing importance value (increase in the mean squared error) for each predictor variable using a permutation procedure. The package "party" allows computing conditional permutation importance, i.e. measure the importance of a variable given the other variables in the model (Strobl, Boulesteix, et al., 2008; Strobl, Hothorn, et al., 2009). The package "ranger" computes unconditional permutation importance and provides options for statistical testing of the estimated values. Here, computation of variable importance and associated p-values with "ranger" were done using the permutation (N=10,000) approach of Altmann et al., 2010. For both approaches, RFs were performed on the complete dataset with a total of 10,000 trees. Analyses were done considering values ranging from 2 to 13 for the "mtry" parameter, which determine the number of variables to possibly split at in each node. For each value of "mtry", twenty independent replicates of the analyses were done using different values for the initial seed (generated using the random number generator of the R package "random" v. 0.2.6). Considering the size of the datasets (13 values for each genetic estimates), the "cforest" analysis was run by setting the "minsplit" (minimal number of observations for splitting) and "minbucket" (minimal number of observation in terminal nodes) parameters to 6 and 3, respectively. The shape of the relationships between cityscape features and punctual genetic estimates was assessed using the "partial_dependence" function from the R package "edarf" (ZM Jones et al., 2017) and the accumulated local effects (ALE) plot (R package "DALEX" – Biecek, 2018) ), when using "cforest" and "ranger", respectively. ALE plots use differences in predictions instead of averages which is more efficient with highly correlated predictors (Apley, 2018). Both methods gave the same results: the variables ranked by "cforest" as the most important were the ones found as significant when using "ranger" (seven variables for $a_r$ and local - $F_{ST}$ and six variables for $H_s$).

#### Figure S3.1 - Importance values of cityscape variables for punctual genetic estimates

Violin-plots of the permutation importance values of each nested cityscape feature according to the "mtry" parameter value (from 2 to 13) for the conditional (cforest) and unconditional (ranger) RF methods applied on A) $a_r$, B) local - $F_{ST}$ and C) $H_s$. For each "mtry" value, 20 independent replicates based on different seeds were analyzed. Each row of the graphs corresponds to a buffer size from 300m to 1500m.

**A)**

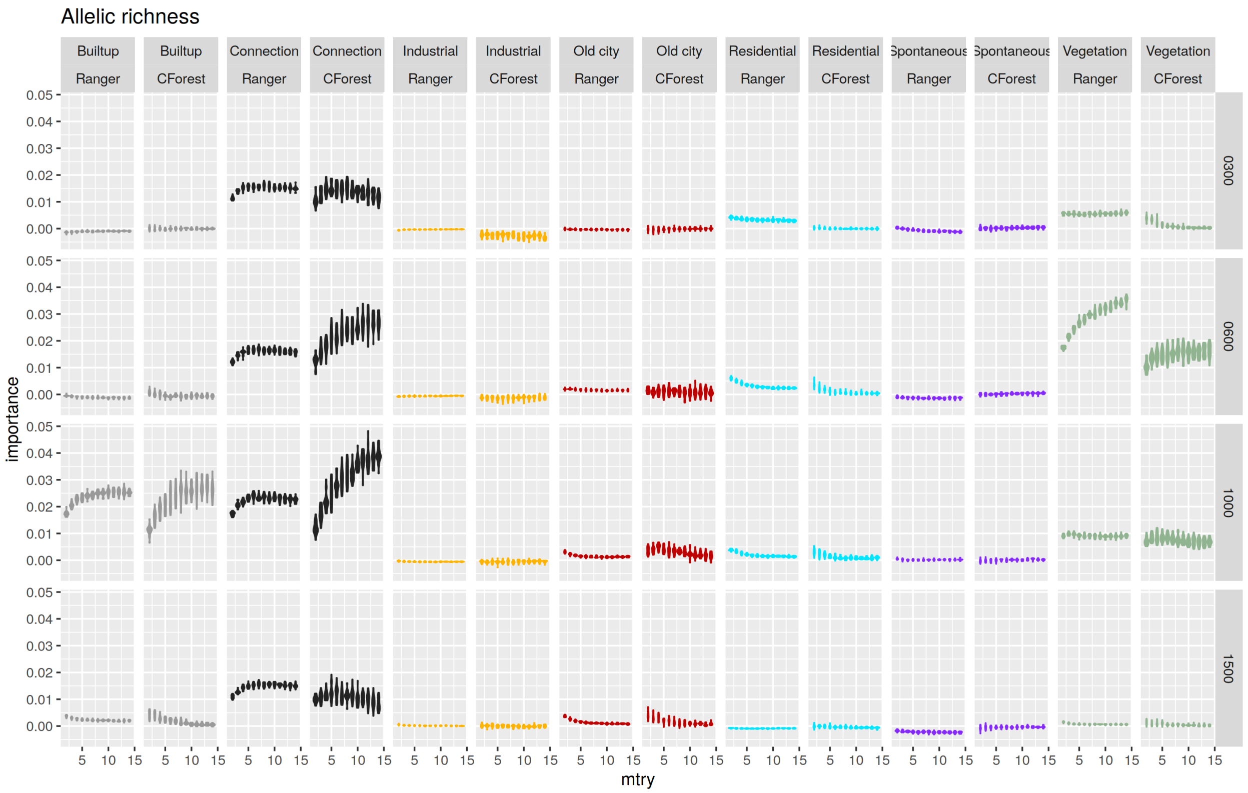

B)

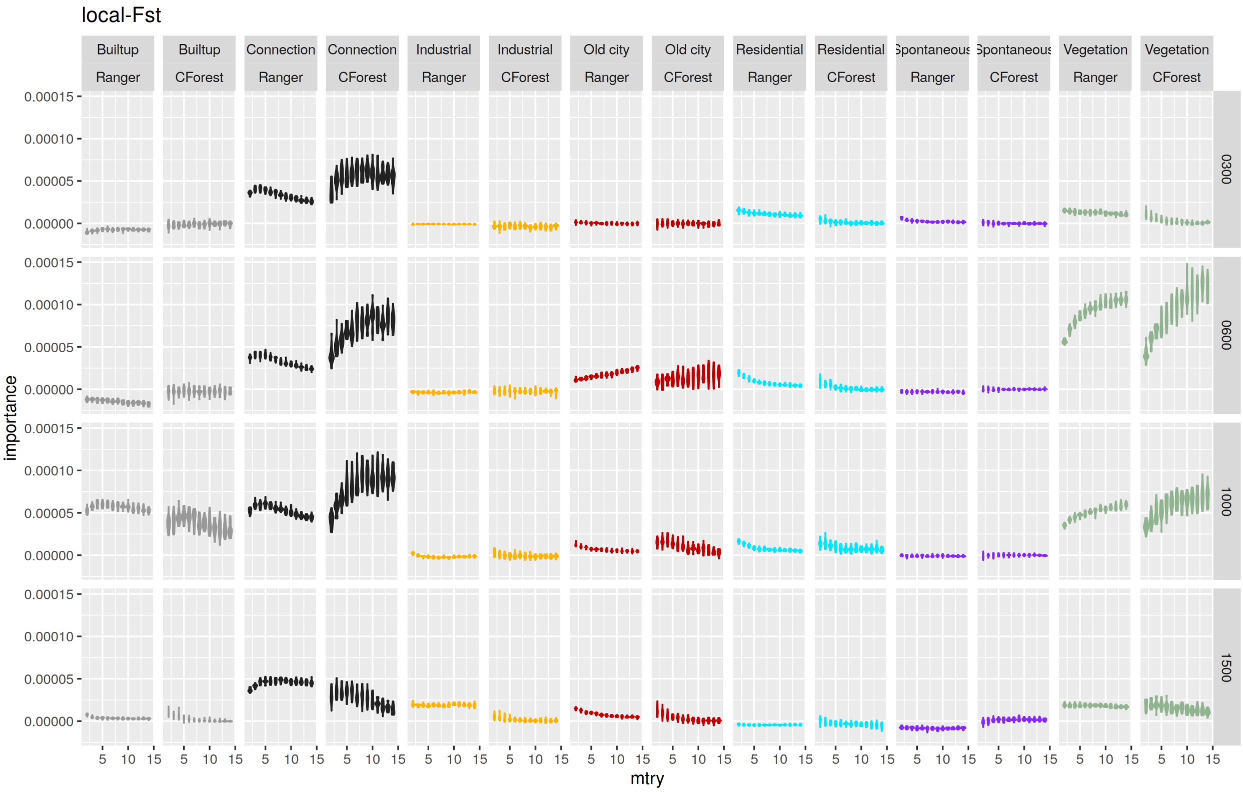
local-Fst
Builtup Ranger
Builtup CForest
Connection Ranger
Connection CForest
Industrial Ranger
Industrial CForest
Old city Ranger
Old city CForest
Residential Ranger
Residential CForest
Spontaneous Ranger
Spontaneous CForest
Vegetation Ranger
Vegetation CForest
0300
0600
1000
1500
importance
0.00015
0.00010
0.00005
0.00000
mtry
5 10 15

C)

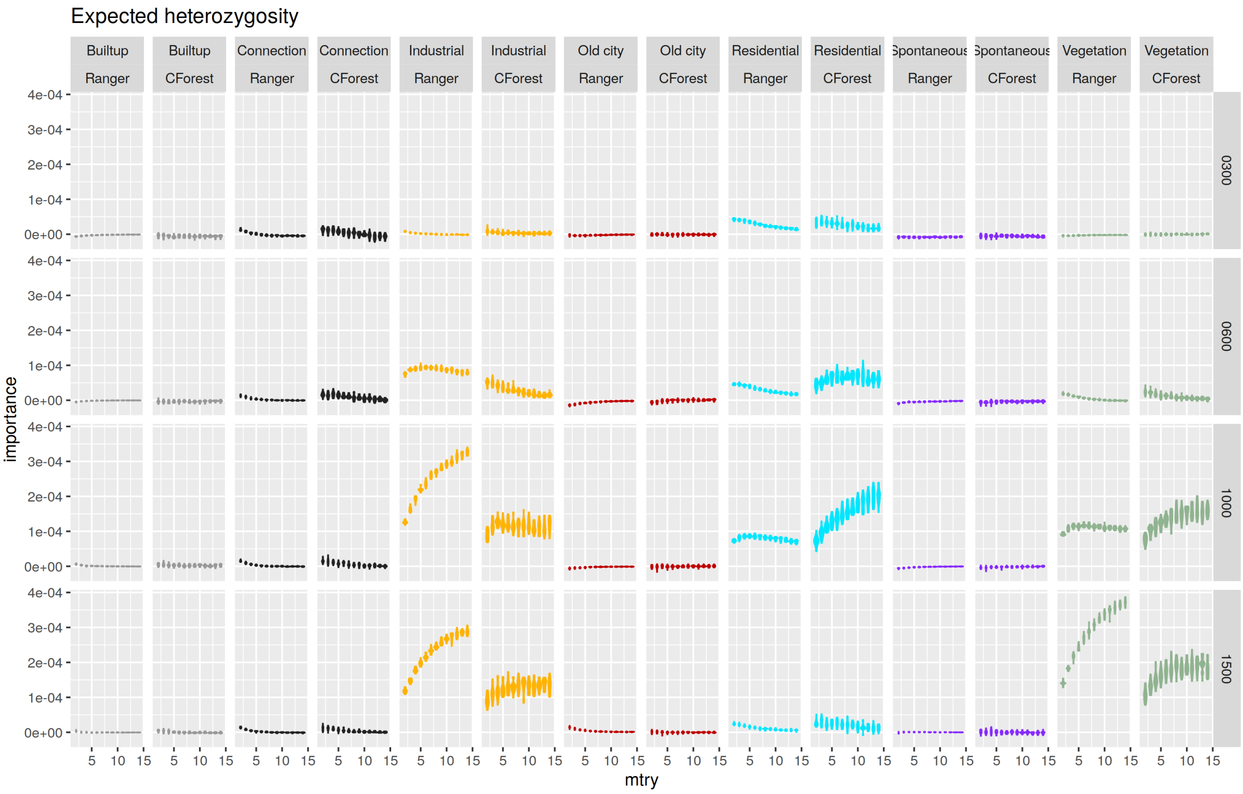
Expected heterozygosity
Builtup Ranger
Builtup CForest
Connection Ranger
Connection CForest
Industrial Ranger
Industrial CForest
Old city Ranger
Old city CForest
Residential Ranger
Residential CForest
Spontaneous Ranger
Spontaneous CForest
Vegetation Ranger
Vegetation CForest
0300
0600
1000
1500
importance
4e-04
3e-04
2e-04
1e-04
0e+00
mtry
5 10 15

#### Figure S3.2 - Relationships between cityscape variables and punctual genetic estimates

Illustration of the shape of the relationships between the most important cityscape features and the punctual genetic estimates of diversity ($a_r$ and $H_s$) and differentiation (local - $F_{ST}$) assessed using A) partial dependence plots for the "cforest" conditional RF method and B) ALE plots for the "ranger" RF method.

**A)**

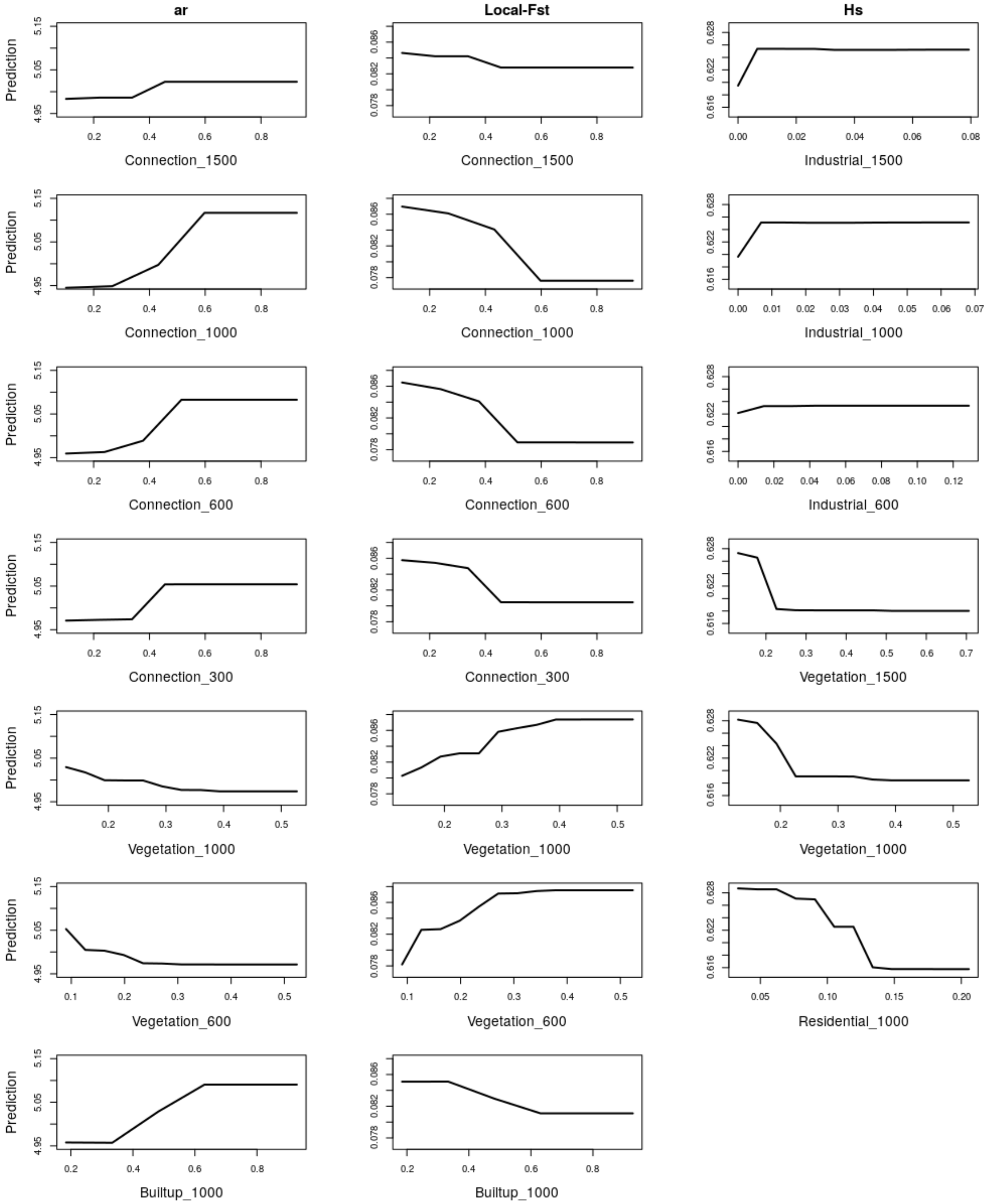

**B)**

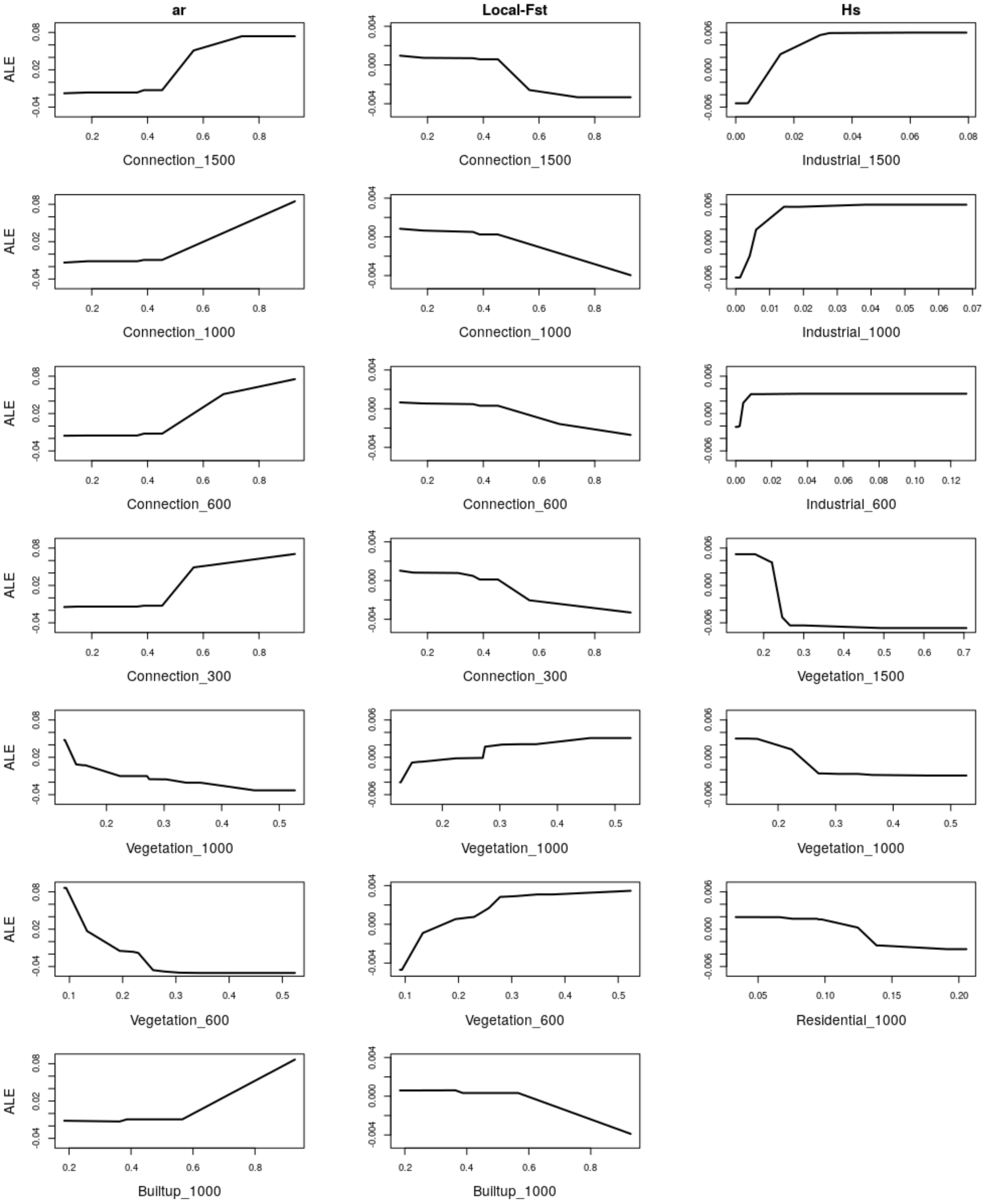
ar
Local-Fst
Hs
ALE
Connection_1500
Industrial_1500
Connection_1000
Industrial_1000
Connection_600
Industrial_600
Connection_300
Vegetation_1500
Vegetation_1000
Vegetation_600
Residential_1000
Builtup_1000

#### Table S3.3 - Spearman correlations for the pairwise approach

Spearman correlation coefficient ($R_s$) and p-values estimated for all pairs of variables computed within the MAPI grid cells.

| Variable 1 | Variable 2 | $R_s$ | p-value |
|---|---|---|---|
| Smoothed-$F_{ST}$ | Old_City | -0.48 | 0.000 |
| Smoothed-$F_{ST}$ | Spontaneous | -0.32 | 0.000 |
| Smoothed-$F_{ST}$ | Residential | 0.16 | 0.015 |
| Smoothed-$F_{ST}$ | Industrial | -0.47 | 0.000 |
| Smoothed-$F_{ST}$ | Vegetation | 0.65 | 0.000 |
| Smoothed-$F_{ST}$ | Built-up | -0.33 | 0.000 |
| Smoothed-$F_{ST}$ | Connection | -0.65 | 0.000 |
| Old_City | Spontaneous | 0.35 | 0.000 |
| Old_City | Residential | 0.27 | 0.000 |
| Spontaneous | Residential | 0.26 | 0.000 |
| Old_City | Industrial | 0.16 | 0.013 |
| Spontaneous | Industrial | -0.13 | 0.044 |
| Residential | Industrial | -0.13 | 0.048 |
| Old_City | Vegetation | -0.69 | 0.000 |
| Spontaneous | Vegetation | -0.56 | 0.000 |
| Residential | Vegetation | 0.00 | 0.964 |
| Industrial | Vegetation | -0.18 | 0.005 |
| Old_City | Built-up | 0.55 | 0.000 |
| Spontaneous | Built-up | -0.15 | 0.016 |
| Residential | Built-up | 0.14 | 0.034 |
| Industrial | Built-up | 0.35 | 0.000 |
| Vegetation | Built-up | -0.26 | 0.000 |
| Old_City | Connection | 0.71 | 0.000 |
| Spontaneous | Connection | 0.06 | 0.388 |
| Residential | Connection | 0.17 | 0.007 |
| Industrial | Connection | 0.40 | 0.000 |
| Vegetation | Connection | -0.51 | 0.000 |
| Built-up | Connection | 0.69 | 0.000 |

##### Figure S3.3 - Importance values of cityscape variables for the pairwise genetic estimate

Boxplots of the conditional permutation importance of each cityscape feature resulting from the fit of the ”cforest” RF algorithm (10,000 trees) on 100 train datasets build by randomly resampling 50% of the MAPI smoothed-$F_{ST}$. Each train dataset was analyzed using different seeds and ”mtry” values ranging from 2 to 5.

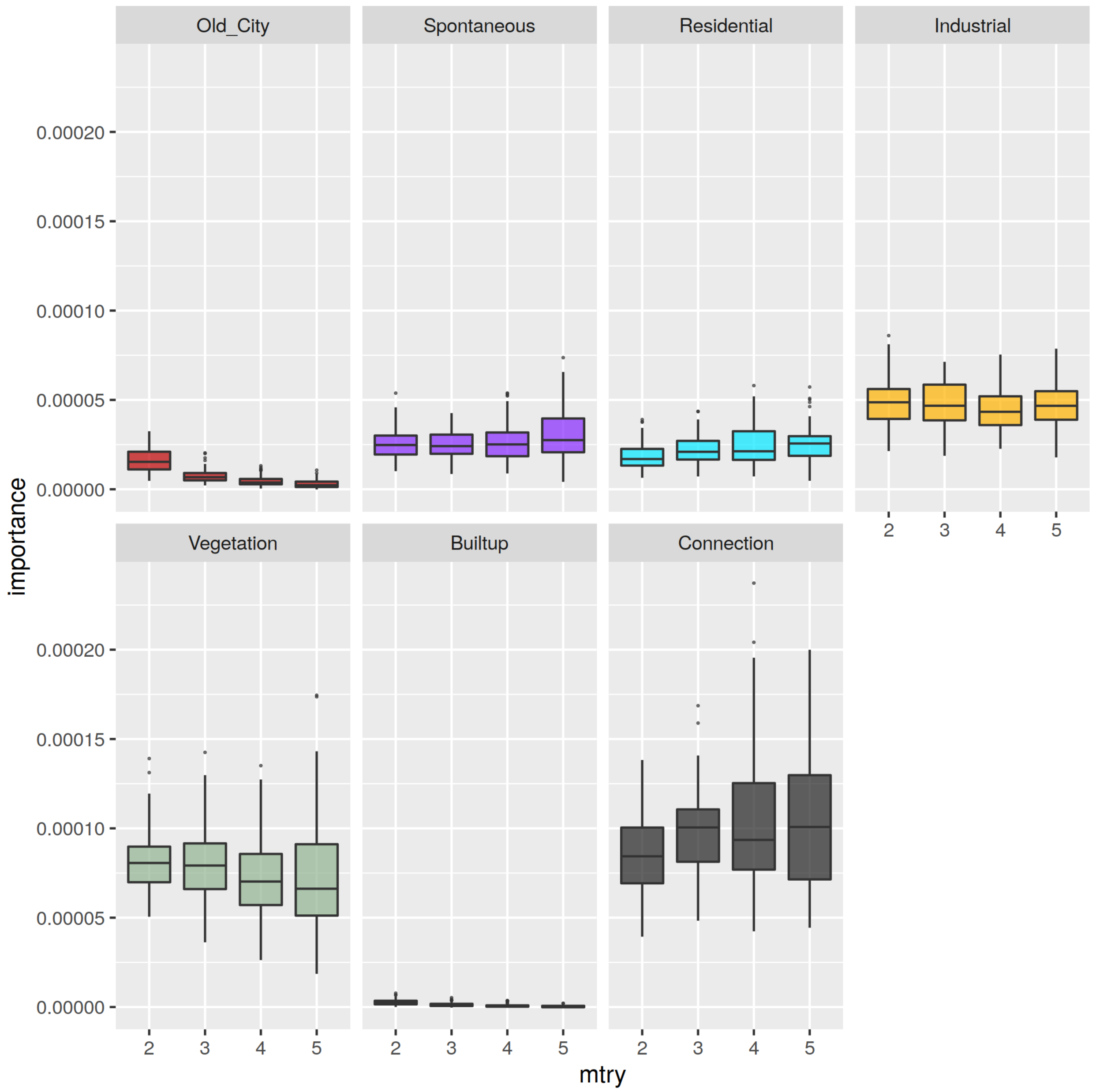

#### Figure S3.4 - Relationships between cityscape variables and the pairwise genetic estimate

Illustration of the shape of the relationships between the most important cityscape features and MAPI smoothed-$F_{ST}$ as assessed using partial dependence plots on "cforest" conditional RF results.

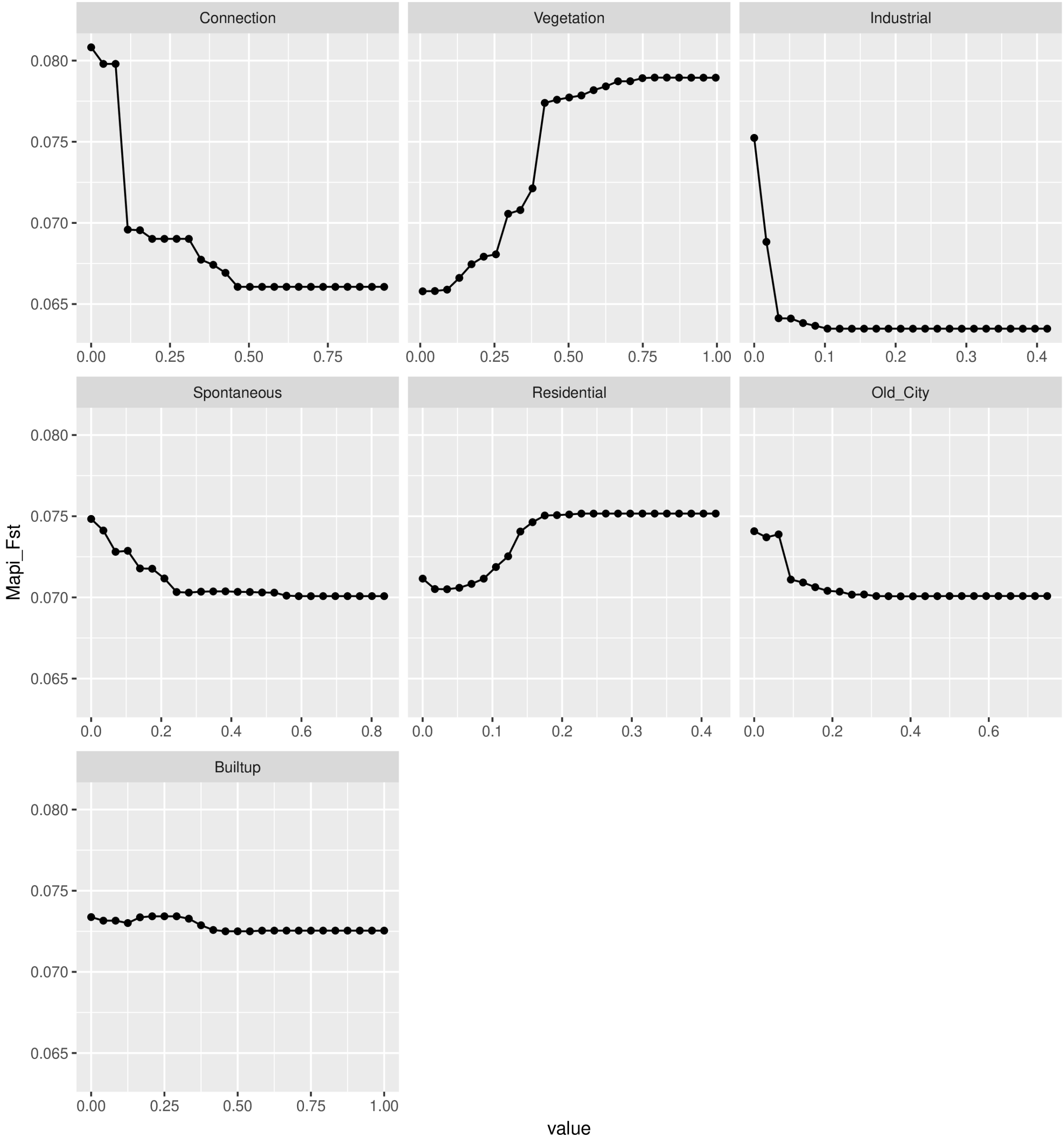

##### General specification of the CAR model for R-INLA

```
form <- Mapi_Fst ~ Old_City + Spontaneous + Residential + Industrial +
     Vegetation + Builtup + Connection +
     f(sites, model = "besag", graph=H, scale.model=TRUE,
          hyper=list(prec=list(initial=log(10^4),
                    prior="pc.prec",
                    param=c(0.025,0.5)
               )
          )
     )

ajust <- inla(form, family="gaussian",
     control.family=list(hyper=list(
               prec=list(initial=log(10^4), prior="pc.prec", param=c(0.025,0.5))
          )),
     data=DON,
     control.predictor=list(compute=TRUE),
     control.compute=list(dic=TRUE, cpo=TRUE, waic=TRUE, config=TRUE),
     verbose=FALSE
)
```

**Figure S3.5 - Spatial residuals of the Bayesian CAR model**

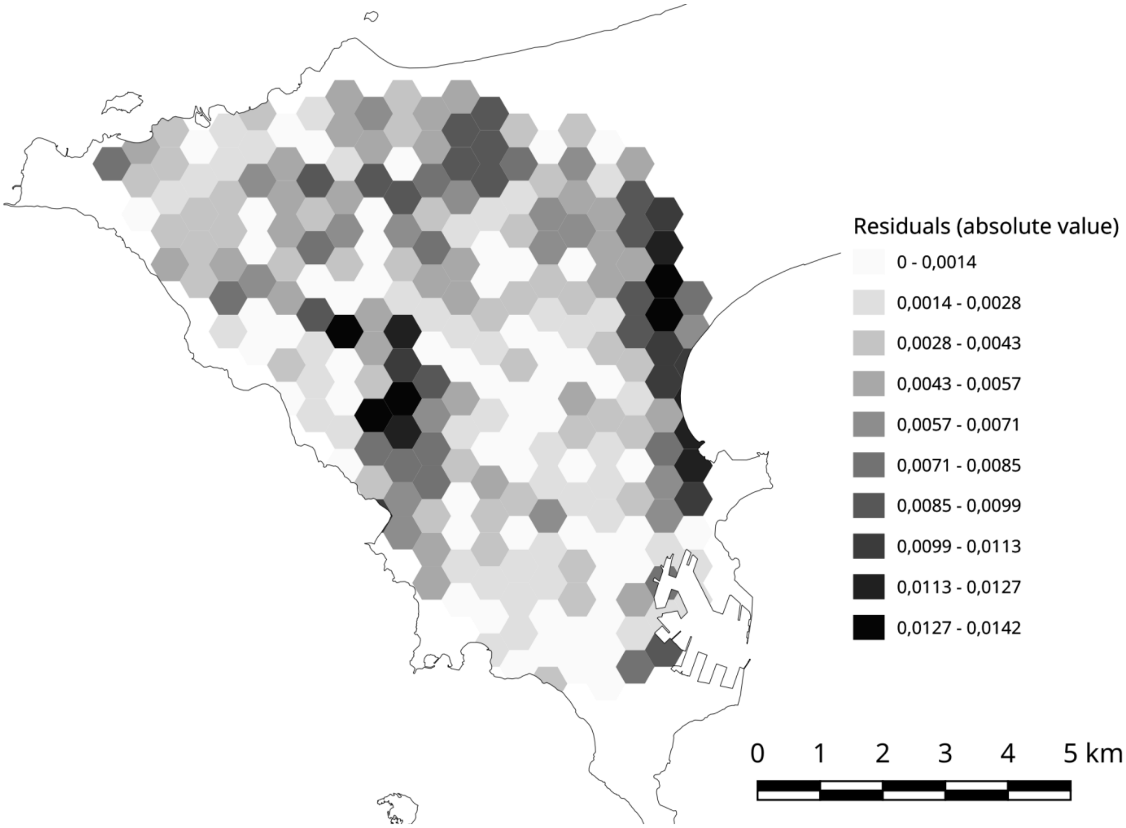

### 4 Analysis of simulated and subsampled datasets

#### Simulation framework

We used simulated datasets as described in Piry et al., 2016 to assess the reliability of the combination of the MAPI and Bayesian CAR methods to test the effect of landscape features on genetic patterns. Data were simulated using the forward-time generation-by-generation algorithm implemented in the software SimAdapt (Rebaudo et al., 2013) to simulate individual genotypes at 10 microsatellite markers within a landscape raster of 58x52 cells mimicking a transition from a favourable to an unfavourable (carrying capacity = 20 and 2; landscape value of 0 and 1, respectively) habitat from north to south with a high level of interpenetration between the two habitats (figure S4.1 on the next page; see Piry et al., 2016 and ”Simulations processed” on MAPI website for details on the simulation study). A total of 20 simulated replicates were analyzed. For each replicate, we divided the landscape raster in twelve cells and randomly picked the location of one individual within each cell. A thirteenth location was then selected anywhere within the raster (figure S4.1, on the following page). This resampling was carried out 10 times for each replicate. Once the 13 spatial locations were selected, we built population-like samples by aggregating the 40 closest individuals to each location (mean number of individual by locality in the house mouse dataset =42). The coordinates of a given population were then determined as being the barycenter of all individuals belonging to this population and the maximum distance between the barycenter and the individuals was used to set the ”error radius” parameter in the MAPI analysis. For each of the 200 datasets (20 replicates $\times$ 10 resampling per replicate), a MAPI analysis was run using pairwise $F_{ST}$ and a grid resolution depending on the sampling locations and according to the Nyquist-Shannon sampling theorem under a situation of random sampling (as for the MAPI analysis performed on the House mouse dataset). The pixel values of the landscape raster (0=favorable and 1=unfavorable) were averaged within each cell of the generated MAPI grids to express the unfavorability of the local habitat. For each of the 200 datasets generated and analyzed with MAPI, the impact of the habitat on the genetic pattern was tested using a Bayesian CAR model approach as for the house mouse dataset. A significant positive effect on the MAPI smoothed-$F_{ST}$ values was expected as the averaged landscape value represented the degree of unfavorability.

When analyzing the 200 simulated datasets (20 independent replicates $\times$ 10 resampling of each replicate) with MAPI and a CAR model, we found that the expected significant positive effect (posterior probability $>$ 0.95) of the proportion of unfavorable habitat on MAPI smoothed-$F_{ST}$ for 95.5% of the simulated datasets. None of the dataset showed an unexpected significant negative effect.

#### Figure S4.1 - Illustration of the analysis process on simulated datasets

A) Landscape raster for the simulation process performed with the software SimAdapt (Rebaudo et al., 2013) - the favourable and unfavourable habitats are represented in black and white, respectively; B) Result of a MAPI analysis run using the individual genetic distance (Rousset, 2000) computed on the output of the simulation that includes 2000 individual genotypes (represented by black dots); C) Result of a MAPI analysis run using pairwise $F_{ST}$ values computed on one resampled dataset with 13 sites sampled by aggregating 40 individual genotypes.

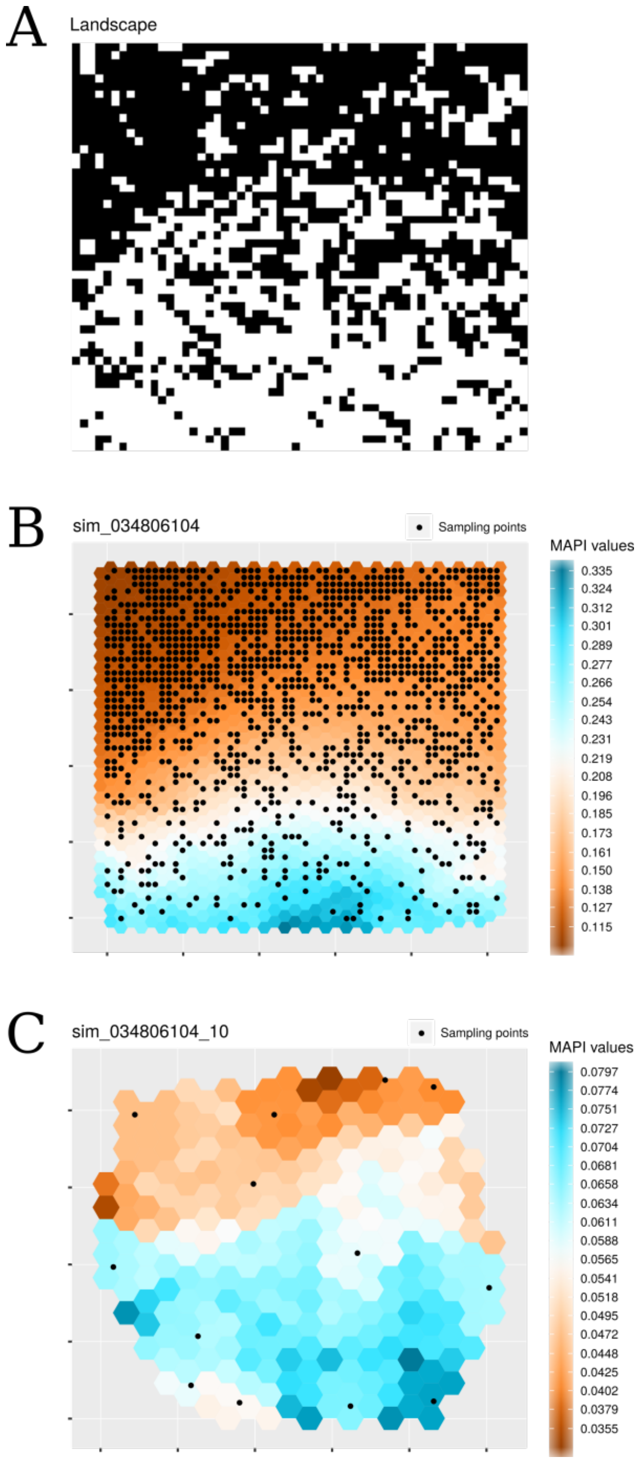

#### Subsampling framework

For N ranging from 6 to 12, we independently subsampled 100 times N sampling sites from the original dataset (13 sites). The approach combining the MAPI analysis and Bayesian CAR model was applied on each replicate of the different subsampling size N. As different sampling sites were subsampled for each replicate of N, the MAPI grid resolution was determined independently for each subsampling scheme by setting the "beta" parameter to 0.25 in respect of the Nyquist-Shannon sampling theorem under a situation of random sampling. Cityscape feature values were computed within the cells of the MAPI grids as described in the main text. The fit of the CAR model was checked using Conditional Predictive Ordinate (CPO) and Probability Integral Transformation (PIT) values. Kolmogorov-Smirnov tests were applied on PIT values to confirm that they did not significantly depart from a uniform distribution as expected if model assumptions are correct.

Graphical outputs of MAPI retrieved a gradient-like pattern in genetic differentiation from lowest smoothed-$F_{ST}$ values in the eastern part of the peninsula to highest values in the western part, whatever the sampling size N considered (varying from 6 to 12). The Bayesian CAR model applied on the MAPI results showed that the main effects retrieved from the complete dataset were observable even with N=6 sampling sites: high posterior probability for a negative effect of "Connection" and "Industrial" and positive effect of "Vegetation", "Residential" and "Built-up" variables. Posterior probability variability decreased rapidly for these variables when the number of sampling site increased (Figure S4.2, on the next page). For the variables "Old_city" and, more especially, "Spontaneous", the number of sampling sites sampled was crucial to obtain clear trends from posterior probabilities across the replicates.

#### Figure S4.2 - Bayesian CAR model on resampled datasets

Boxplots of the posterior probabilities across 100 independent replicates for each subsampling size varying from 6 to 12 sites. The number of replicates for which the CAR model was correctly fitted is indicated in bracket for each sample size on the x-axis. Considering a threshold of 0.05, posterior probabilities > 0.95 indicates a significant positive effect while posterior probabilities < 0.05 indicates a significant negative effect of the variable.

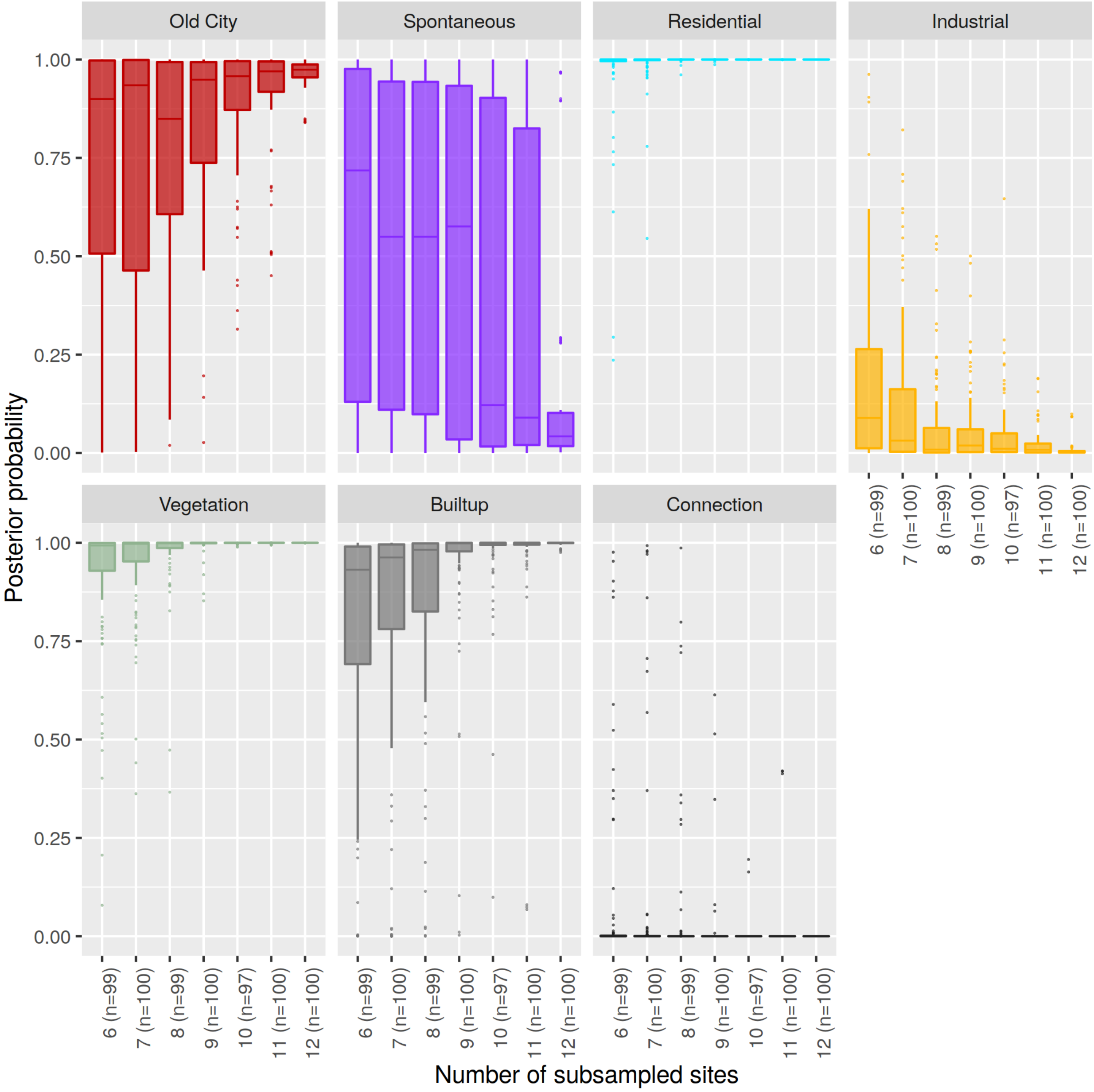
