## Appendix for "Interplay between historical and current features of the cityscape in shaping the genetic structure of the house mouse (*Mus musculus domesticus*) in Dakar (Senegal, West Africa)"

Claire Stragier[1], Sylvain Piry[2], Anne Loiseau[2], Mamadou Kane[1], Aliou Sow[1], Youssoupha Niang[1], Mamoudou Diallo[1], Arame Ndiaye[1], Philippe Gauthier[3], Marion Borderon[4], Laurent Granjon[3], Carine Brouat*[3], and Karine Berthier*[5]

[1] *BIOPASS (IRD-CBGP, ISRA, UCAD), Campus de Bel-Air, BP 1386, CP 18524 Dakar, Senegal*
[2] *CBGP, INRAE, CIRAD, IRD, Montpellier SupAgro, Univ. Montpellier, Montpellier, France*
[3] *CBGP, IRD, CIRAD, INRAE, Montpellier SupAgro, Univ. Montpellier, Montpellier, France*
[4] *Department of Geography and Regional Research, University of Vienna, Austria*
[5] *Pathologie Végétale, INRAE, 84140 Montfavet, France*

**3 Final time series** **18**

*These two authors contributed equally to this work

### 1 Geoprocessing

#### 1.1 Acquisition of historical data

Maps have been downloaded from the web or scanned from paper as ungeoreferenced images. The scanned maps have been georeferenced as ”rasters” using QGis software. Ground control points were used as geo-references. Once georeferenced, buit-up areas have been manually digitized as polygons in QGis as WGS84 (EPSG:4326) latitude/longitude coordinates shapefiles (one per map). Once digitized, shapefiles identified for each year have been imported in a PostgreSQL/PostGIS database using the R pacakge ”sf”.

Geoprocessing has been done using SQL queries. To homogenize map spatial scales, that varied between 1:10,000 and 1:250,000, a smoothing-like procedure was applied by adding a 100m buffer (ST_Buffer) to digitized built-up polygons in order to merge (ST_Union) adjacent or overlapping polygons as a single unit in fine scale maps. New polygons resulting from the merging were then eroded by applying a negative 100m buffer to better respect the original spatial extend of the built-up areas.

For all maps presented below, digitized built-up areas are represented by delineated orange or pink polygons depending on whether they are connected to the first European settlement or not, respectively. The actual coastline is materialized as a blue line. Airport lanes were not considered as built-up areas.

### 2 Georeferenced maps and digitized built-up polygons

#### 2.1 1677, Historical Map (figure 1, not used)

**Source:** Gallica, Bibliothèque Nationale de France (http://gallica.bnf.fr/ark:/12148/btv1b77590868) Despite the high level of distortion, this map illustrates the presence of autochtonous villages and the absence of European settlements in the Cap-Vert peninsula.

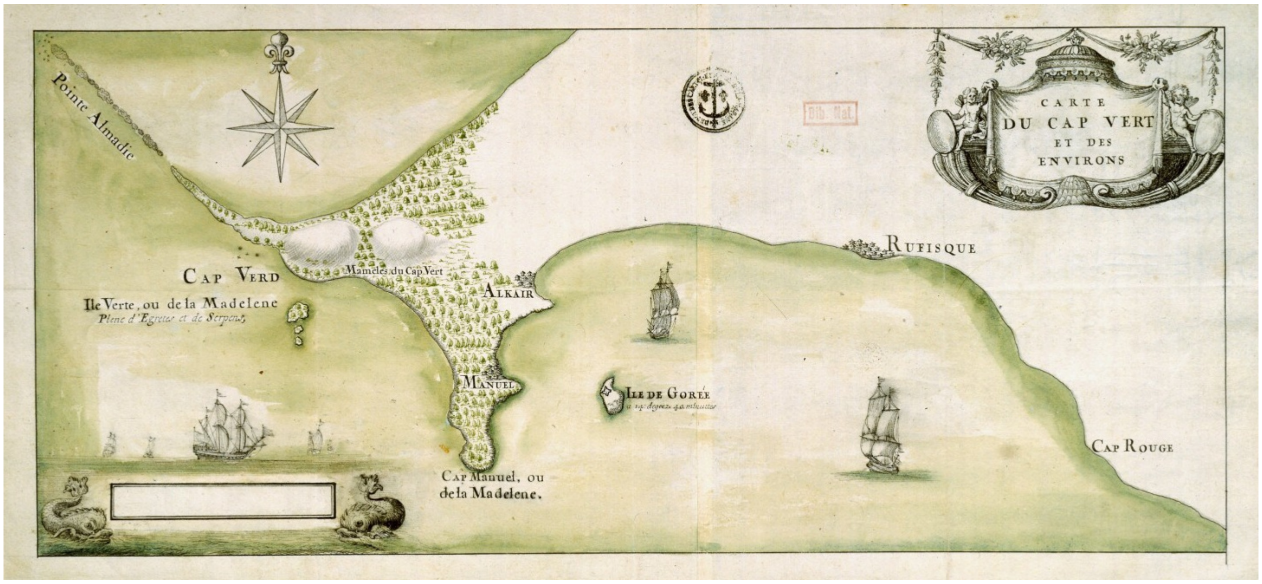

Figure 1: Map of Cap Vert Peninsula and surrounds, 1677

#### 2.2 1850, Historical Map (figure 2)

**Source:** Gallica, Bibliothèque Nationale de France:
http://gallica.bnf.fr/ark:/12148/btv1b53025350r
**Used projection:** EPSG:32628 - WGS 84 / UTM zone 28N - Projected.
This map shows the first European settlement in the Cap-Vert Peninsula.

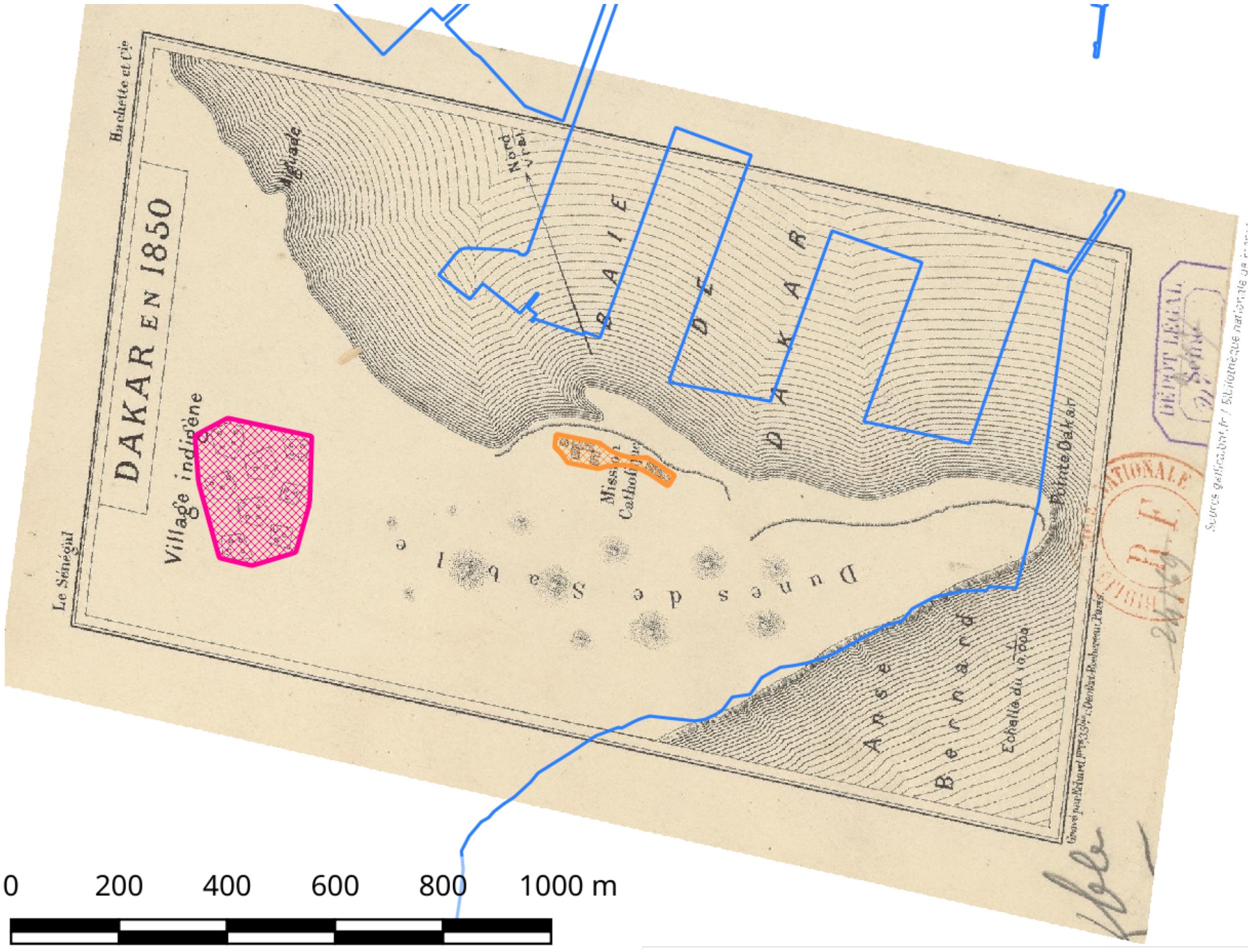

Figure 2: Map of Dakar city in 1850

#### 2.3 1862, French map of Dakar (figure 3)

**Source:** Unknown – J.-M. Duplantier, pers. comm.
**Used projection:** EPSG:32628 - WGS 84 / UTM zone 28N - Projected.

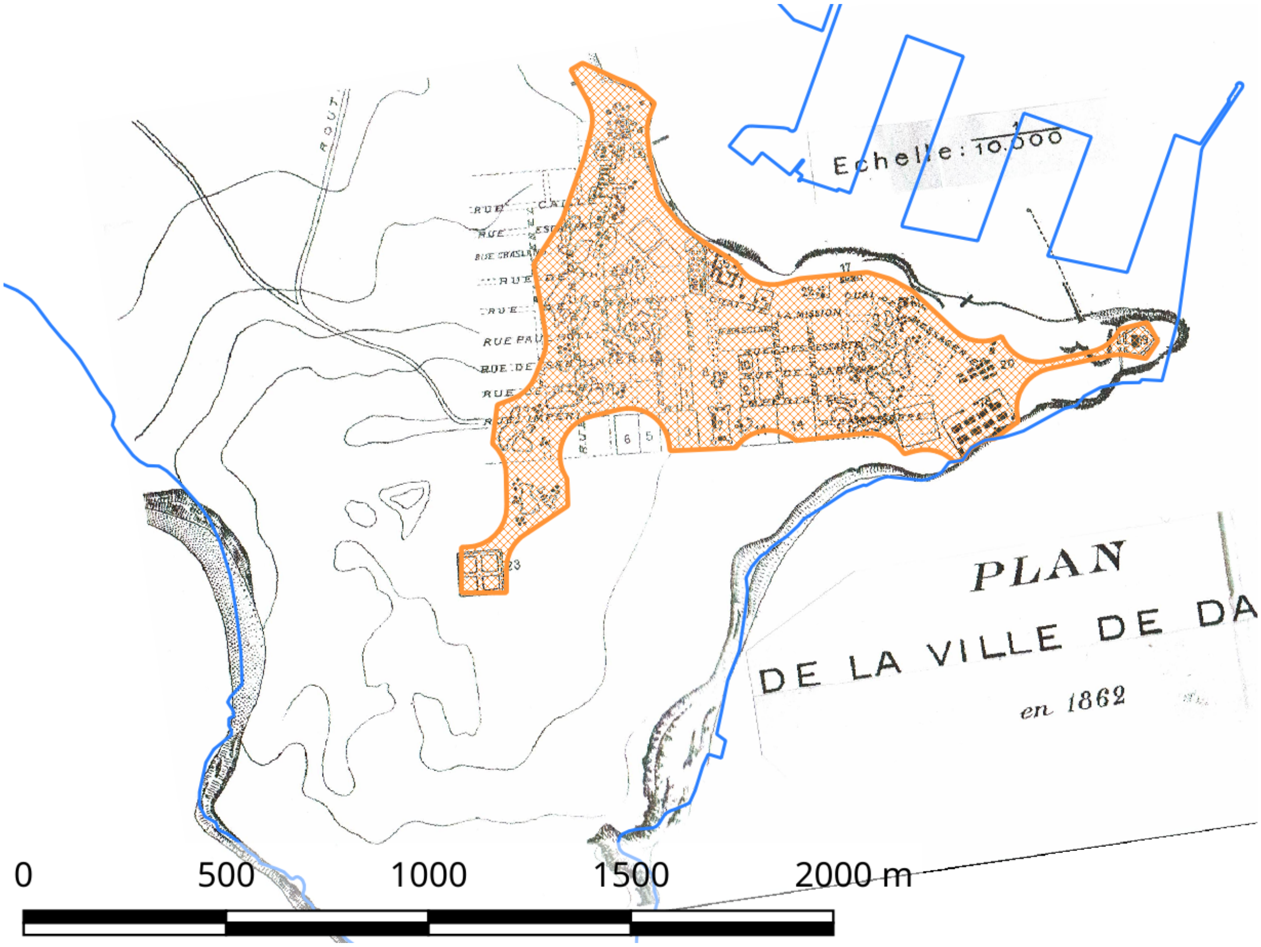

Figure 3: Map of Dakar city in 1862

#### 2.4 1894, French Navigation Map (figure 4)

**Source:** Université Bordeaux Montaigne, France:
http://1886.u-bordeaux-montaigne.fr/items/show/71630
**Processing:** A detailed map of the city was extracted using the Gimp software and georeferenced independantly. Due to high distorsions on the 1894 map, the villages were localized using the 1923 map (figure 7 on page 8).
**Used projection:** EPSG:32628 - WGS 84 / UTM zone 28N - Projected.

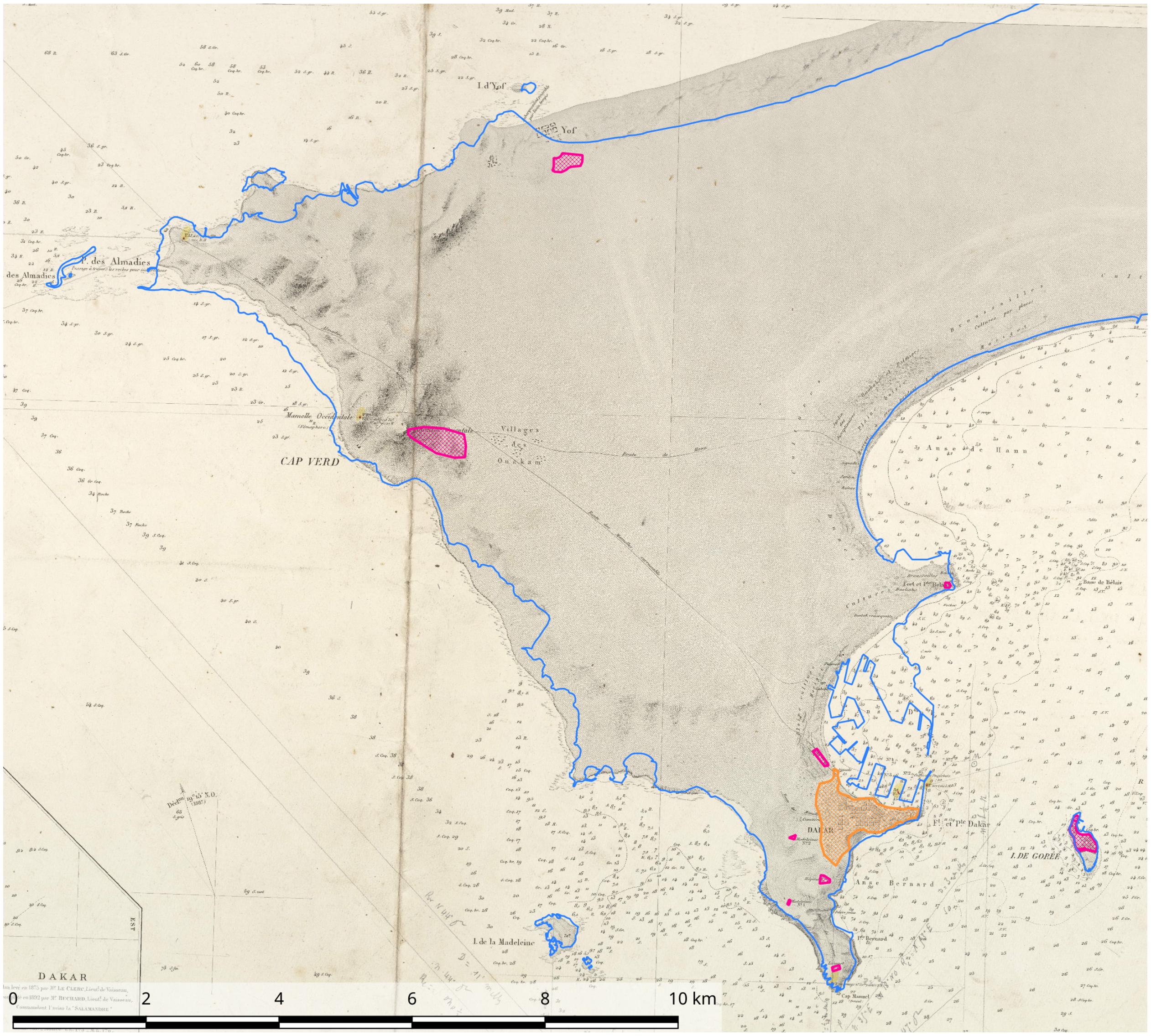

Figure 4: Map of the Cap Vert Peninsula in 1894

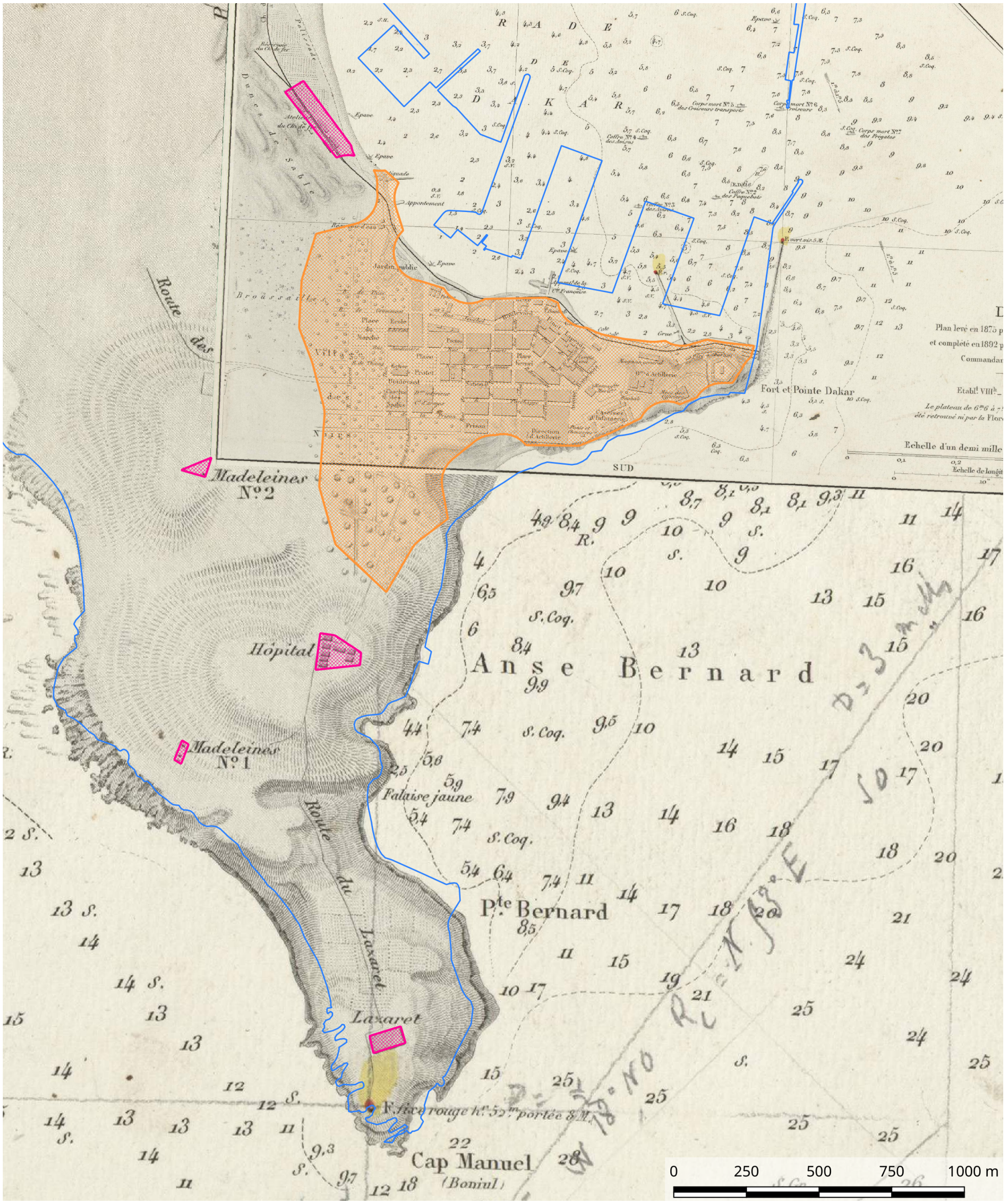

Figure 5: Detail of Dakar city in 1894

##### 2.5 1905, French Military Map (figure 6)

**Source:** Université Bordeaux Montaigne, France:
http://1886.u-bordeaux-montaigne.fr/items/show/9378).
**Used projection:** Ad-hoc (+proj=longlat +a=6378249.2 +b=6356515 +no_defs) - Projected.
**Scale:** 1:100,000.

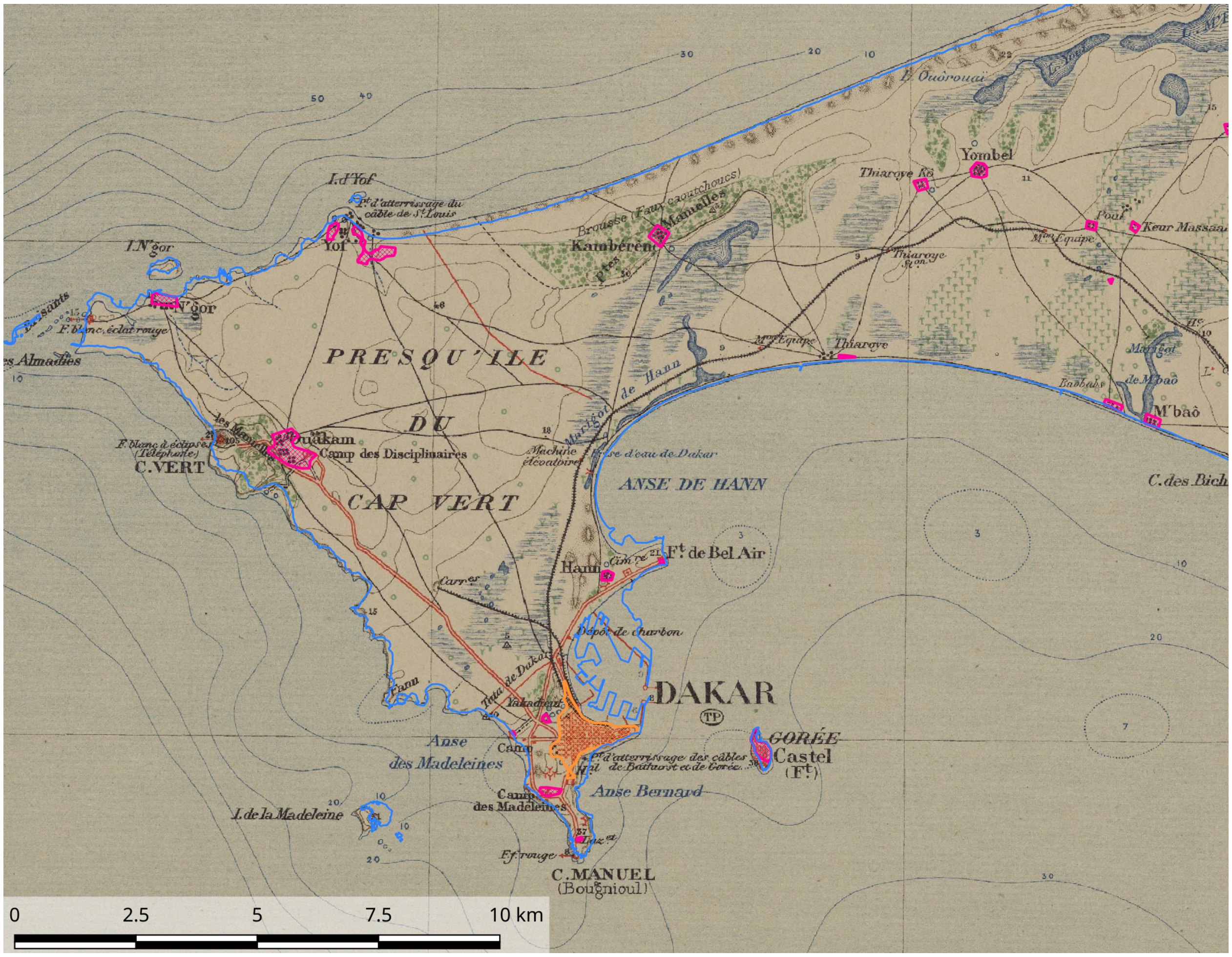

Figure 6: Map of the Cap Vert Peninsula in 1905

#### 2.6 1923, French Military Map (figure 7)

**Sources:** Université Bordeaux Montaigne, France:
http://1886.u-bordeaux-montaigne.fr/items/show/9717,
http://1886.u-bordeaux-montaigne.fr/items/show/9718,
http://1886.u-bordeaux-montaigne.fr/items/show/9719,
http://1886.u-bordeaux-montaigne.fr/items/show/9720.
**Used projection:** Ad-hoc SCR (+proj=longlat +a=6378249.2 +b=6356515 +no_defs) - Projected.
**Scale:** 1:10,000.
**Pre-processing:** Assembling of four tiles.

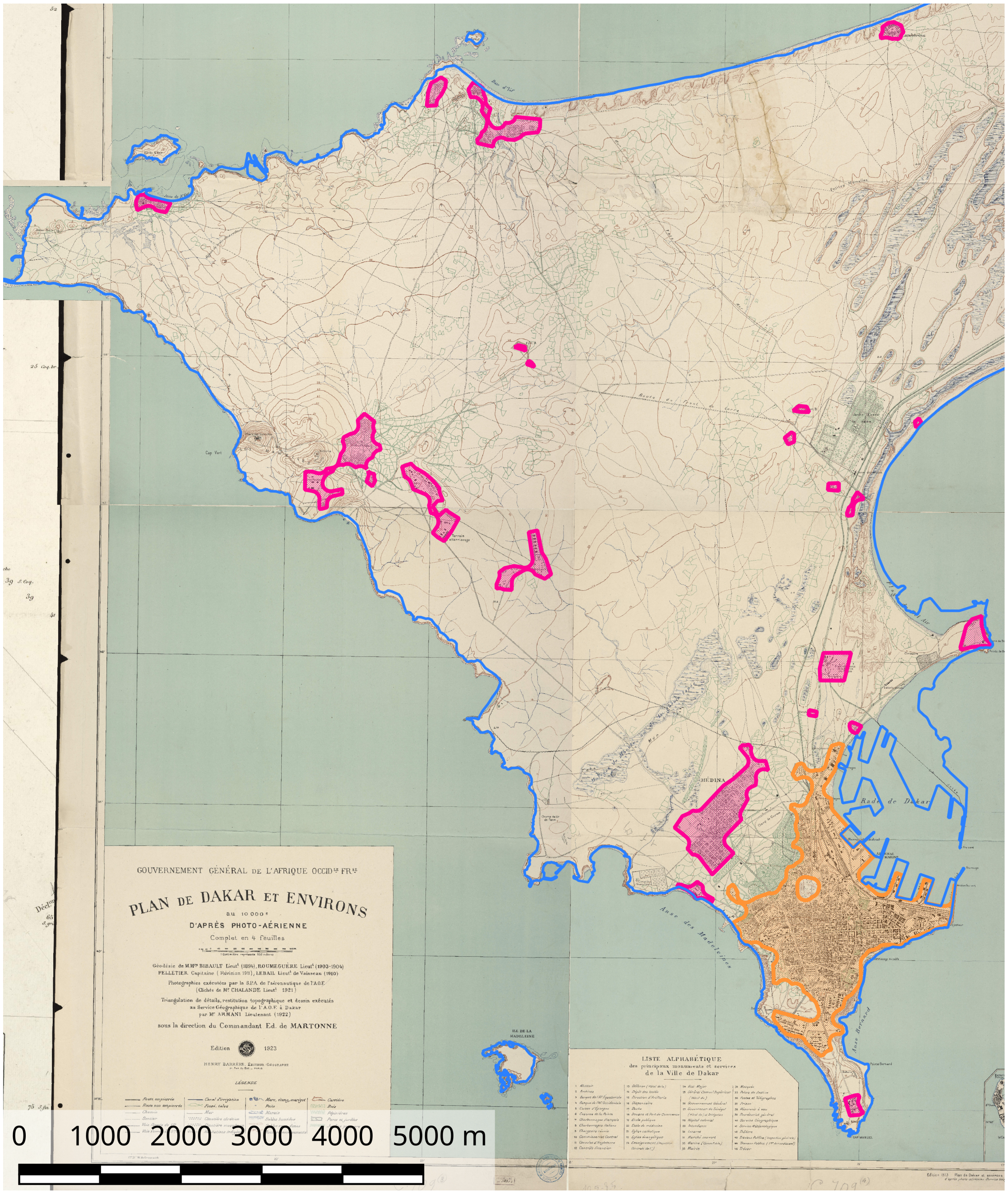

Figure 7: Map of the Cap Vert Peninsula in 1923

##### 2.7 1942, American Military Plan of Dakar (figure 8)

**Source:** Perry-Castañeda Library Map Collection:
http://legacy.lib.utexas.edu/maps/ams/txu-pclmaps-oclc-6595993-dakar.jpg
**Used projection:** EPSG:4326 - WGS 84 - Geographical.
**Post-processing:** The part of the peninsula outside this map was filled using data from the 1923 map (figure 7 on the preceding page)).

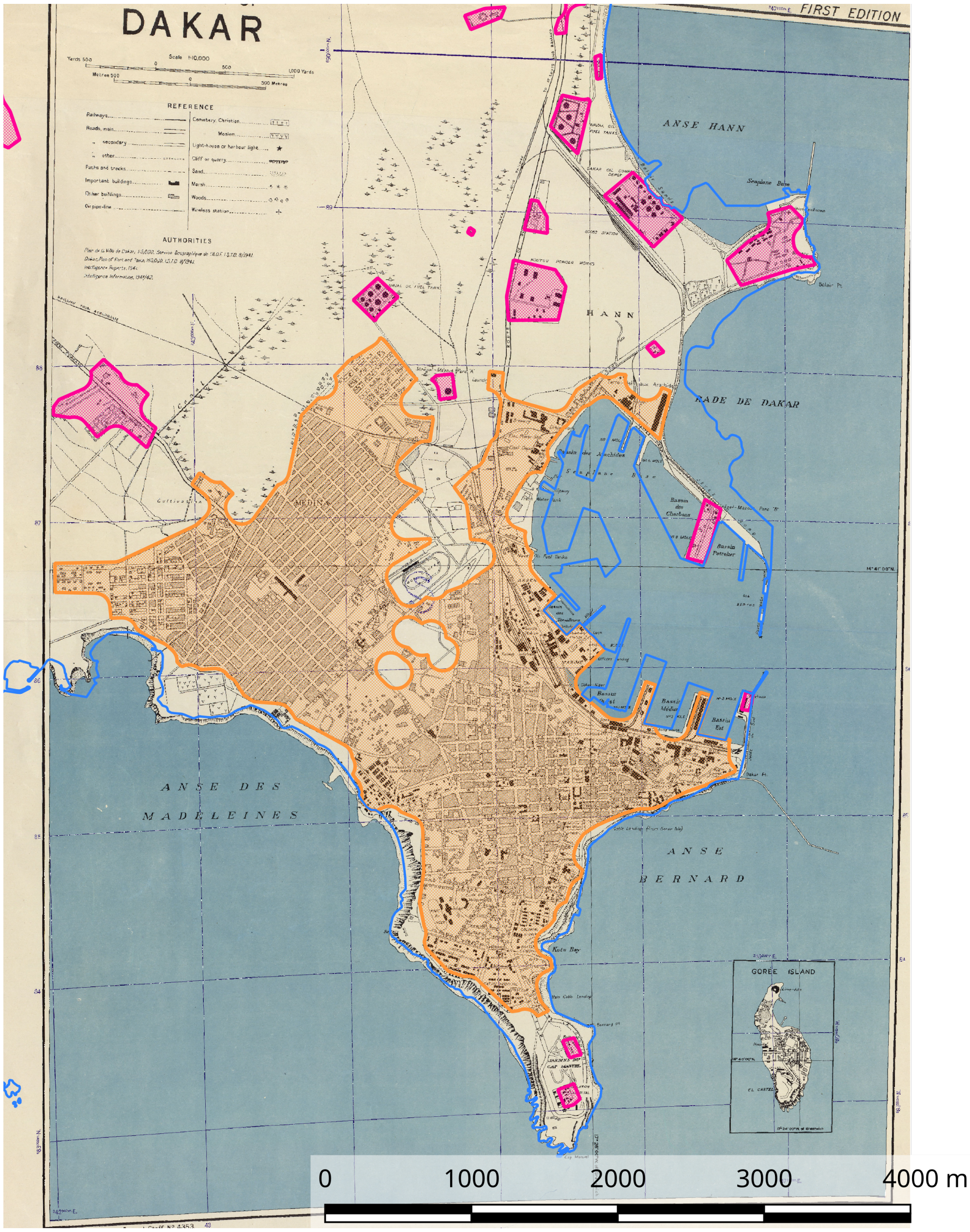

Figure 8: Town Plan of Dakar in 1942

#### 2.8 1953, American Military Maps (figures 9 and 10 on the next page)

**Source:** Perry-Castañeda Library Map Collection:
http://legacy.lib.utexas.edu/maps/ams/west_africa/txu-oclc-6595921-nd28-5.jpg,
http://legacy.lib.utexas.edu/maps/ams/west_africa/txu-oclc-6595921-nd28-5a.jpg.
**Used projection:** EPSG:4326 - WGS 84 - Geographical.
**Scale:** 1:250,000.
**Post-processing:** The polygons of the city on fig. 9 was retrieved from detailed map included in the same source fig. 10 on the next page.

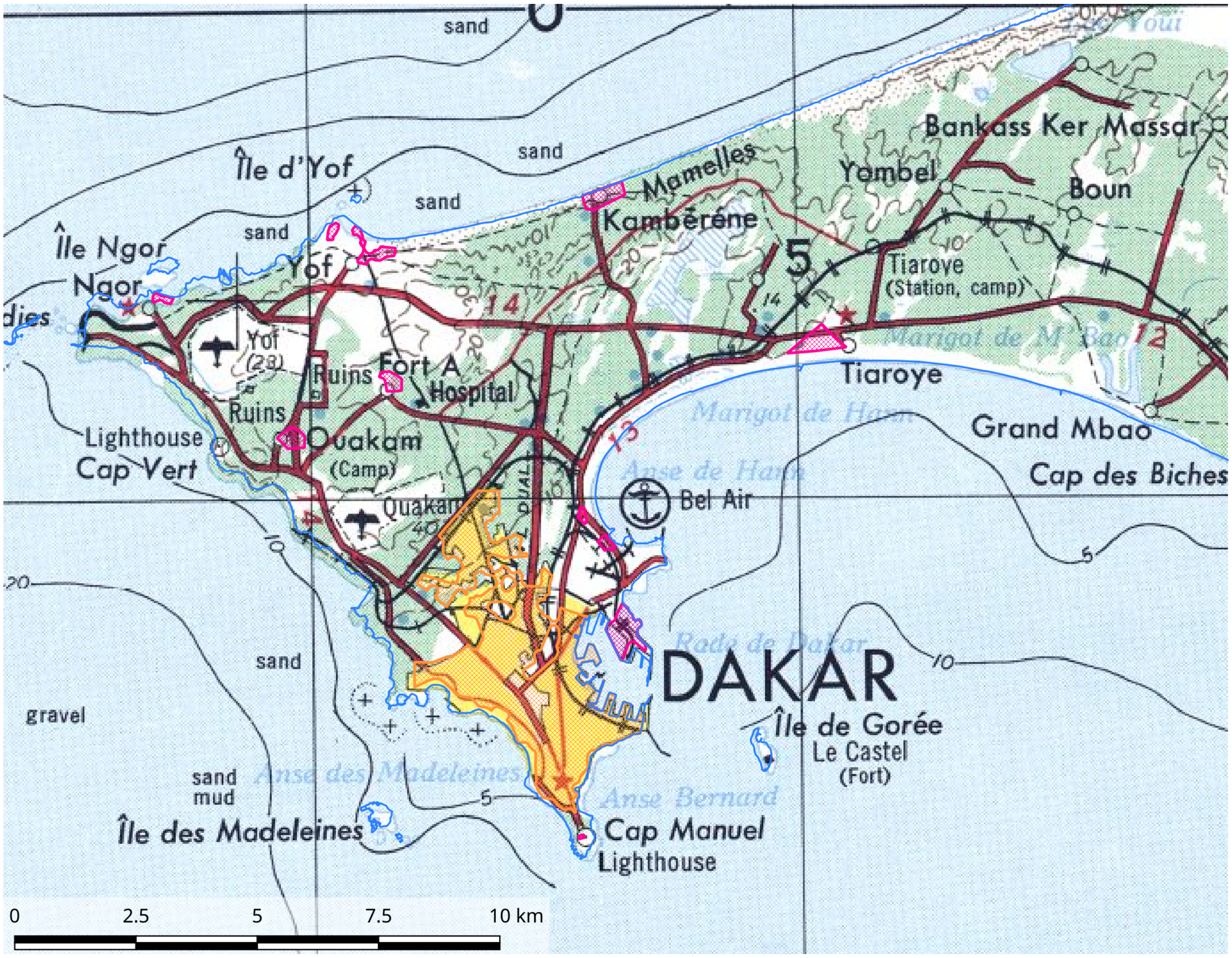

Figure 9: Map of the Cap Vert Peninsula in 1953

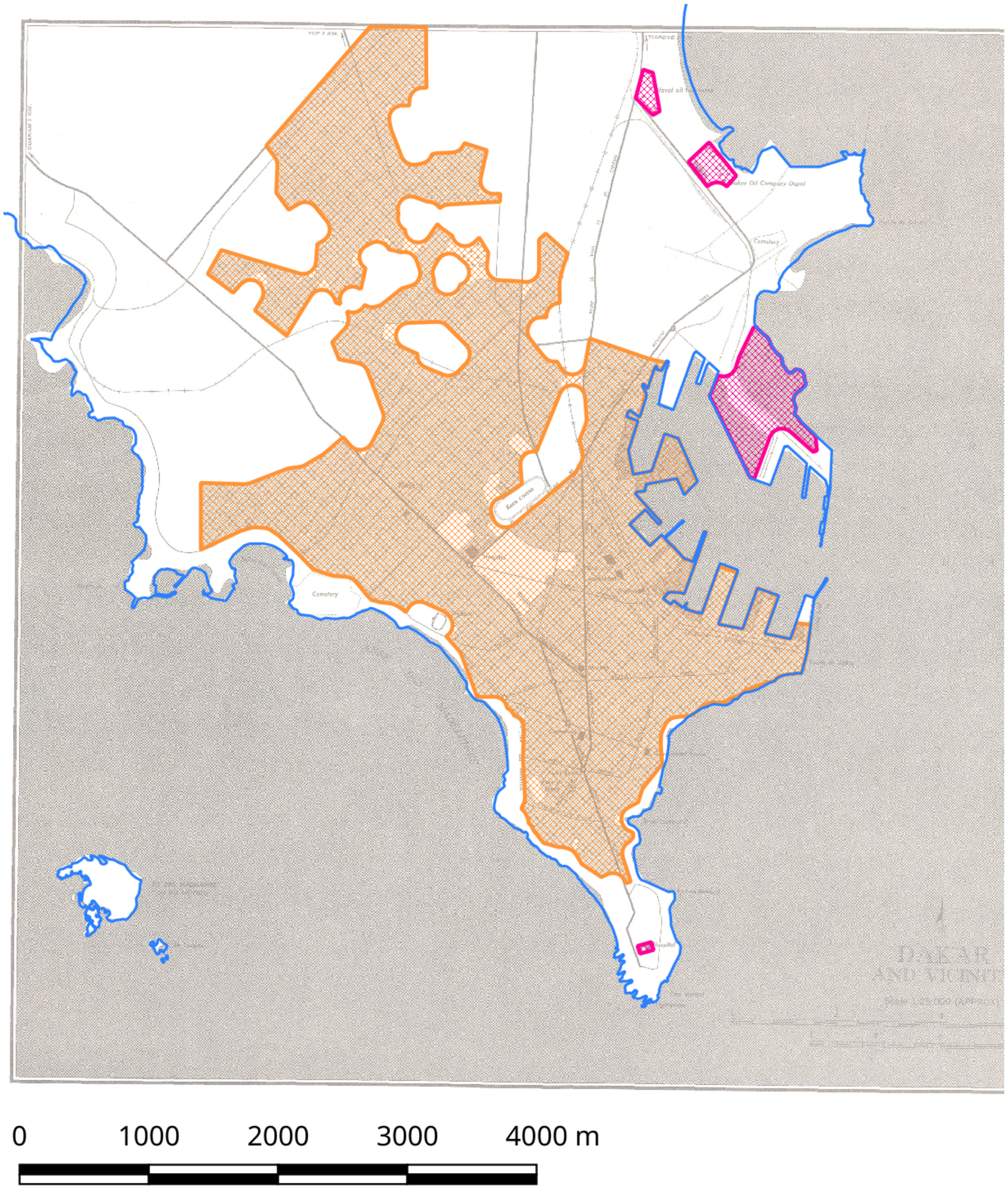

Figure 10: Plan of the City of Dakar in 1953

#### 2.9 1957, Senegalese Geographical Map (figure 11)

**Source:** Scanned by the Senegalese National Geographical Institute.
**Used projection:** EPSG:32628 - WGS 84 / UTM zone 28N - Projected.
**Scale:** 1:200,000.

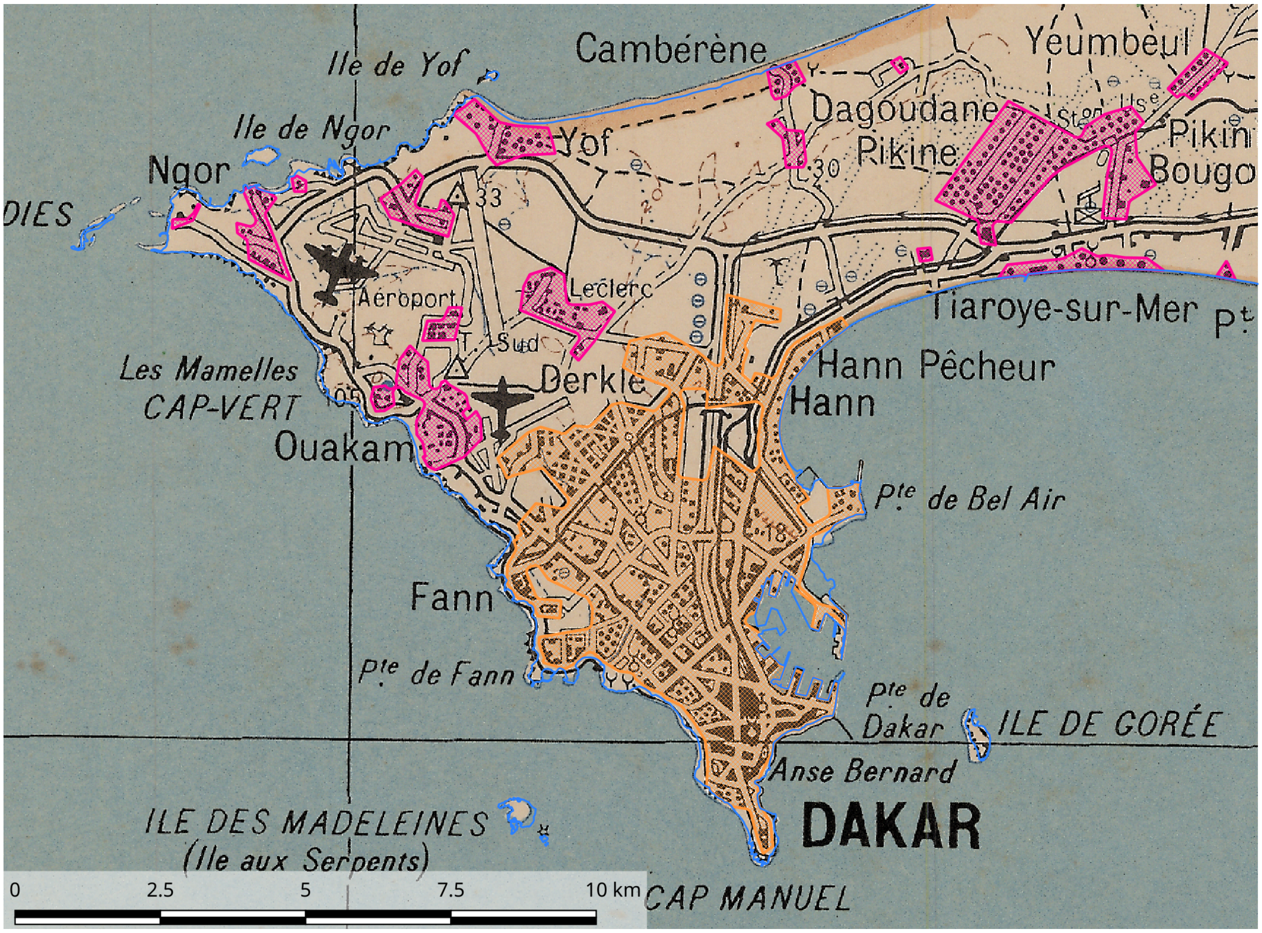

Figure 11: Map of the Cap Vert Peninsula in 1957

#### 2.10 1966, Senegalese Topographical Map (figure 12)

**Source:** Scanned by the Senegalese National Geographical Institute.
**Used projection:** EPSG:32628 - WGS 84 / UTM zone 28N - Projected.
**Scale:** 1:20,000.
**Pre-processing:** The two tiles were georeferenced independantly.

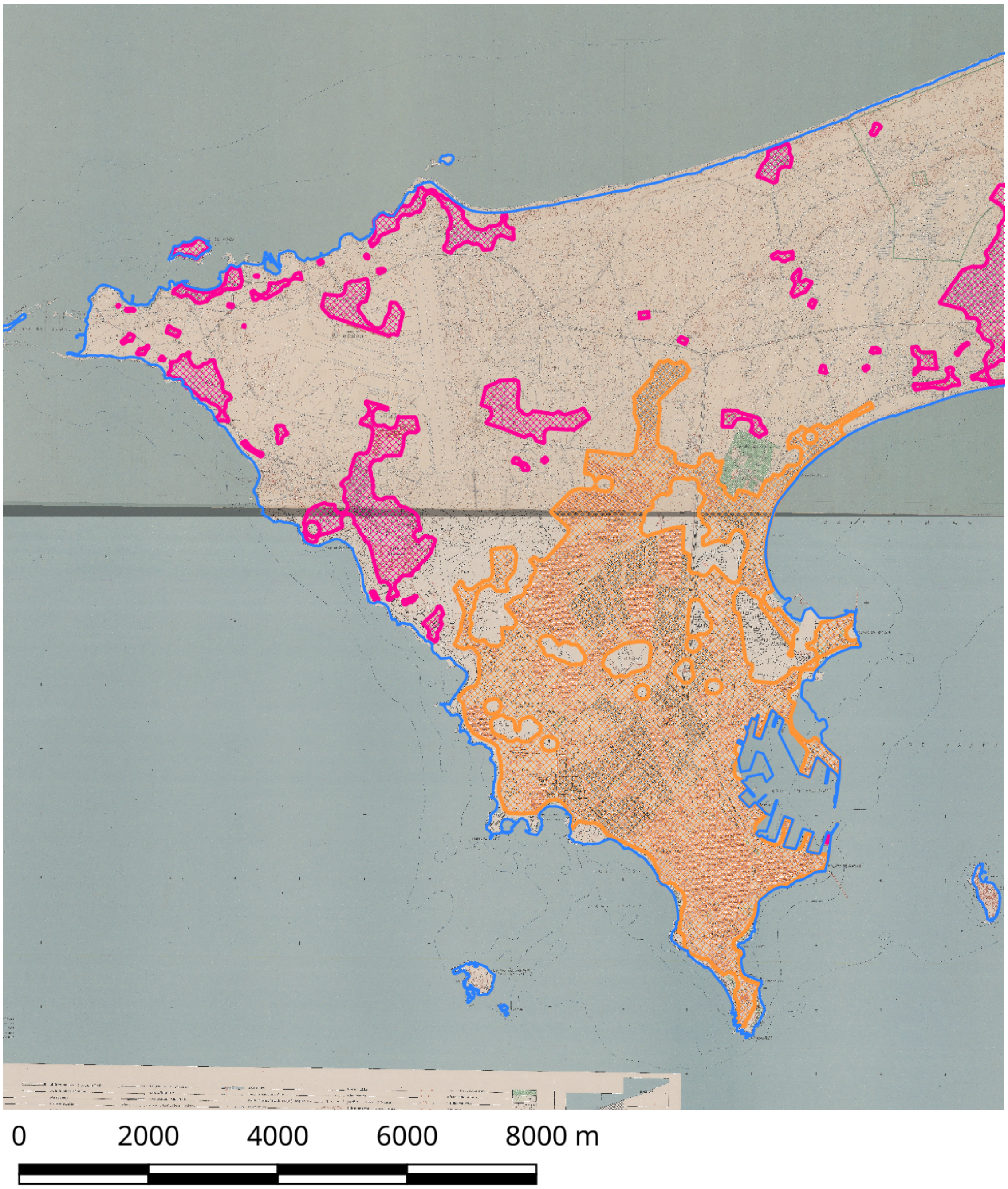

Figure 12: Map of the Cap Vert Peninsula in 1966

#### 2.11 1987, American Aerial Navigation Chart ND 28-05 (figure 13)

**Source:** Perry-Castañeda Library Map Collection (http://legacy.lib.utexas.edu/maps/jog/west_africa/txu-oclc-224327916-nd28-05.jpg).
**Used projection:** EPSG:4326 - WGS 84 - Geographical.
**Scale:** 1:250,000.

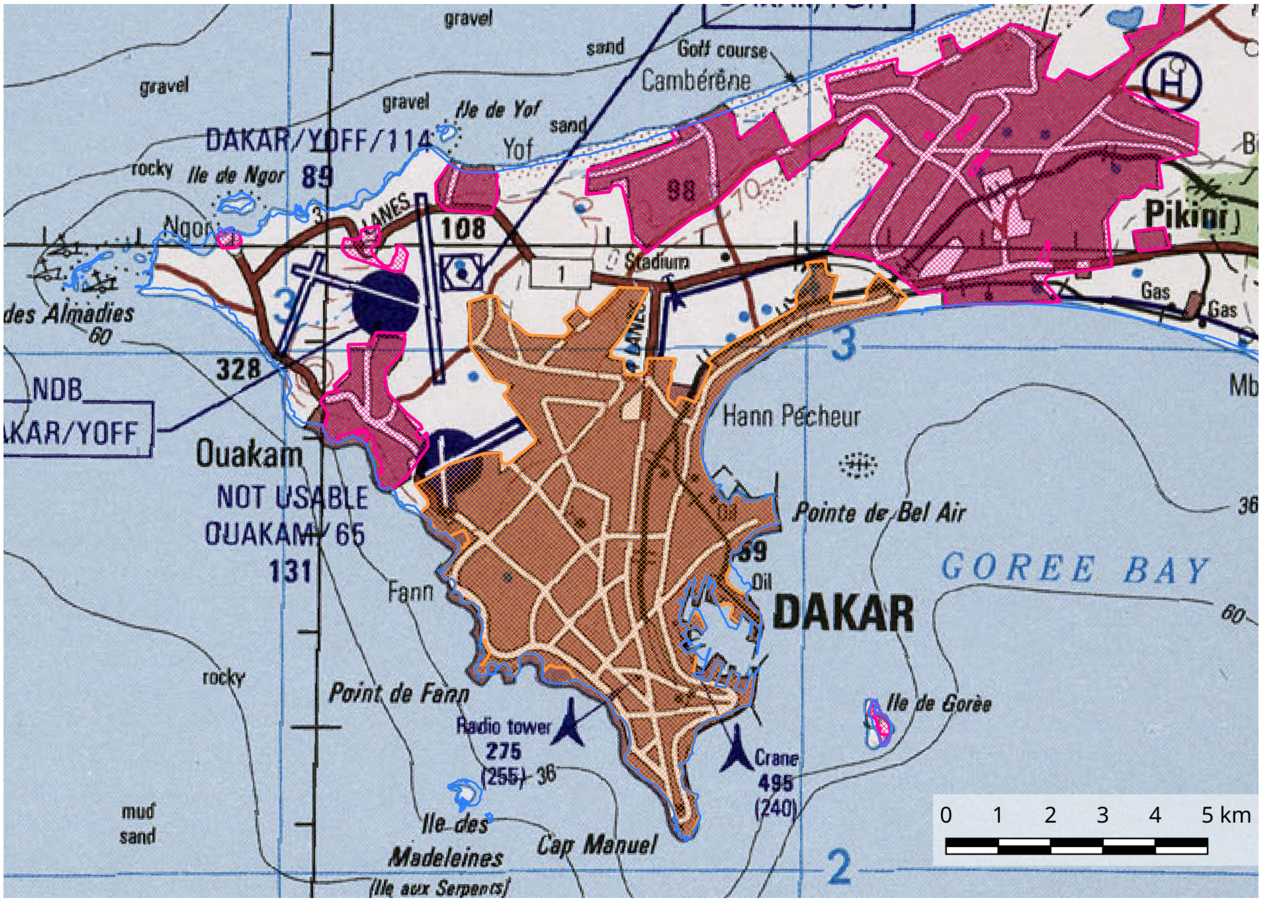

Figure 13: Map of the Cap Vert Peninsula in 1987

#### 2.12 1993, French Road Map (figure 14)

**Source:** French Geographical National Institute, scanned by L. Granjon, pers. comm.
**Used projection:** Ad-hoc (+proj=longlat +a=6378249.2 +b=6356515 +no_defs) - Projected.
**Scale:** unknown.

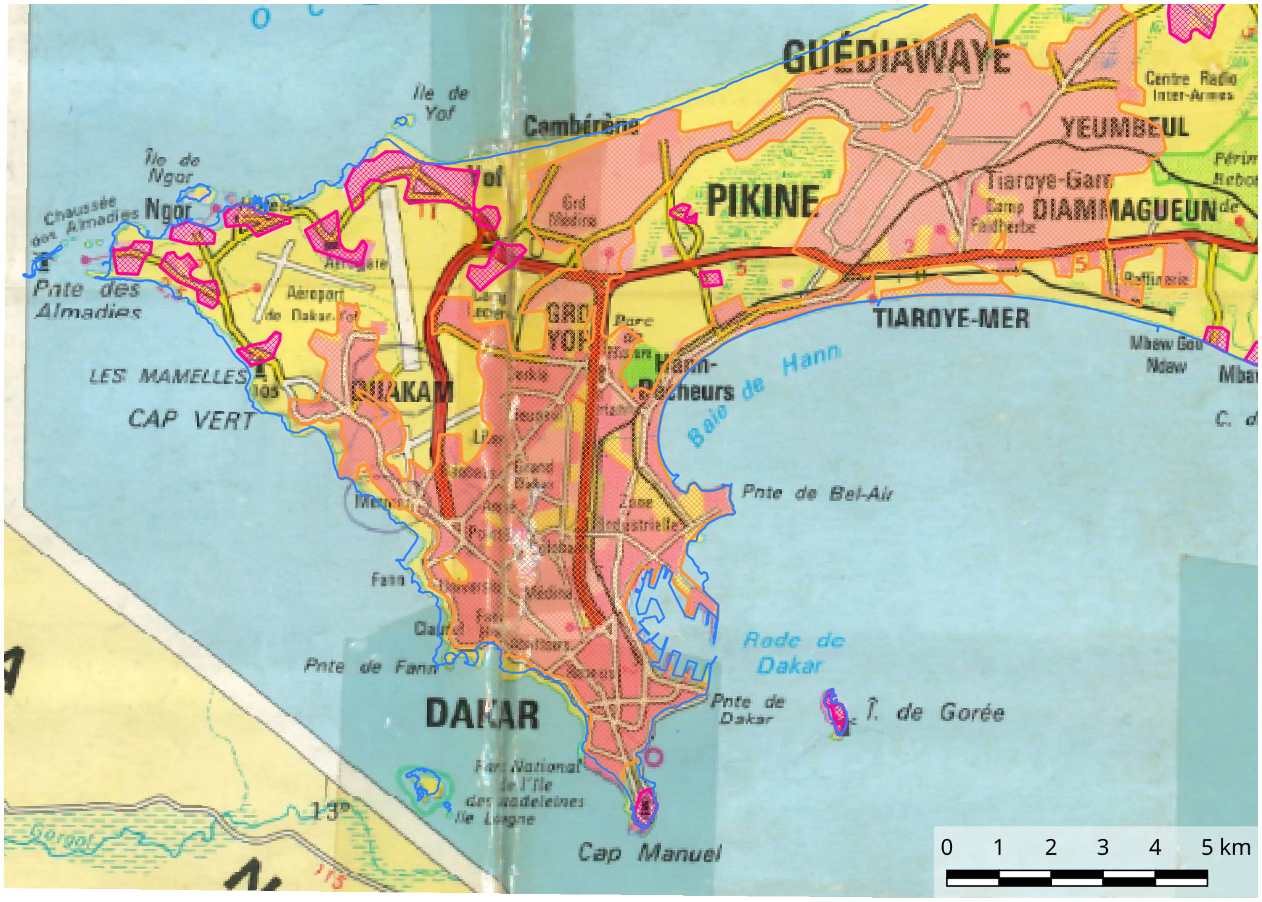

Figure 14: Map of the Cap Vert Peninsula in 1993

#### 2.13 2001, Landsat Image (figure 15)

**Source:** U.S. Geological Survey (https://earthexplorer.usgs.gov/,
Landsat Product Identifier: LE07_L1TP_205050_20010517_20170205_01_T1).
**Pixel size:** 15m (ETM+, band 8)
**Post-processing:** Polygons corresponding to unbuilt areas were delineated from this image and then substracted from the 2018 map (figure 16 on the next page) to generate the 2001 dataset.

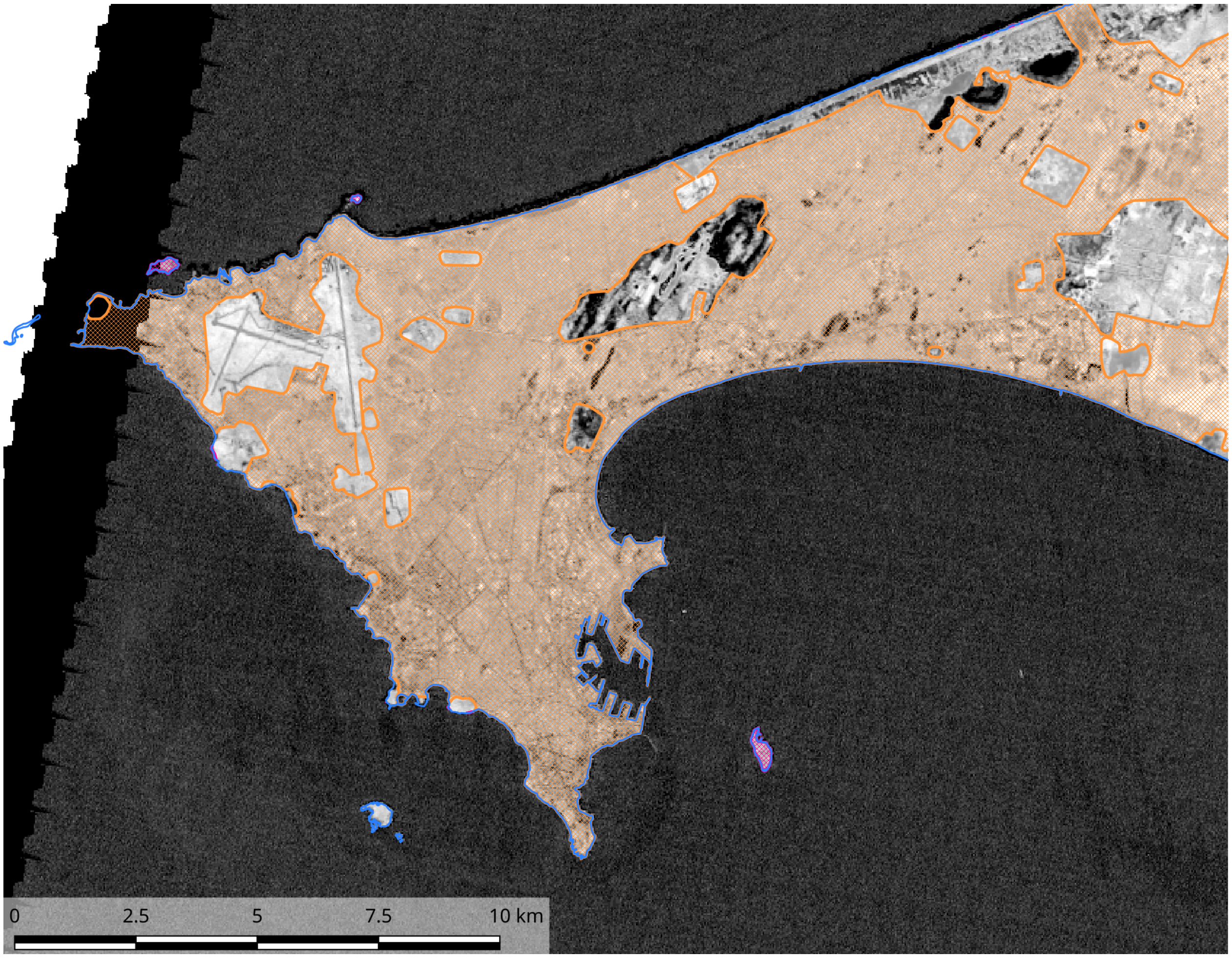

Figure 15: Landsat image (band 8) of the Cap Vert Peninsula in 2001

#### 2.14 2018, OpenStreetMap (figure 16)

**Source:** Open Street Map (XYZ Tiles in QGis with url=http://tile.openstreetmap.org/%7Bz%7D/%7Bx%7D/%7By%7D.png).
**Projection:** EPSG:3857 - WGS 84 / Pseudo Mercator - Projected.
**Processing:** Original polygons were downloaded from OpenStreetMap servers within QGis. The polygons describing coatsline were extracted, along with (Multi)Polygons delineating unbuilt areas. Unbuilt areas were substracted using the QGis function "difference". Visual checking was done using Google maps satellite view within QGis.

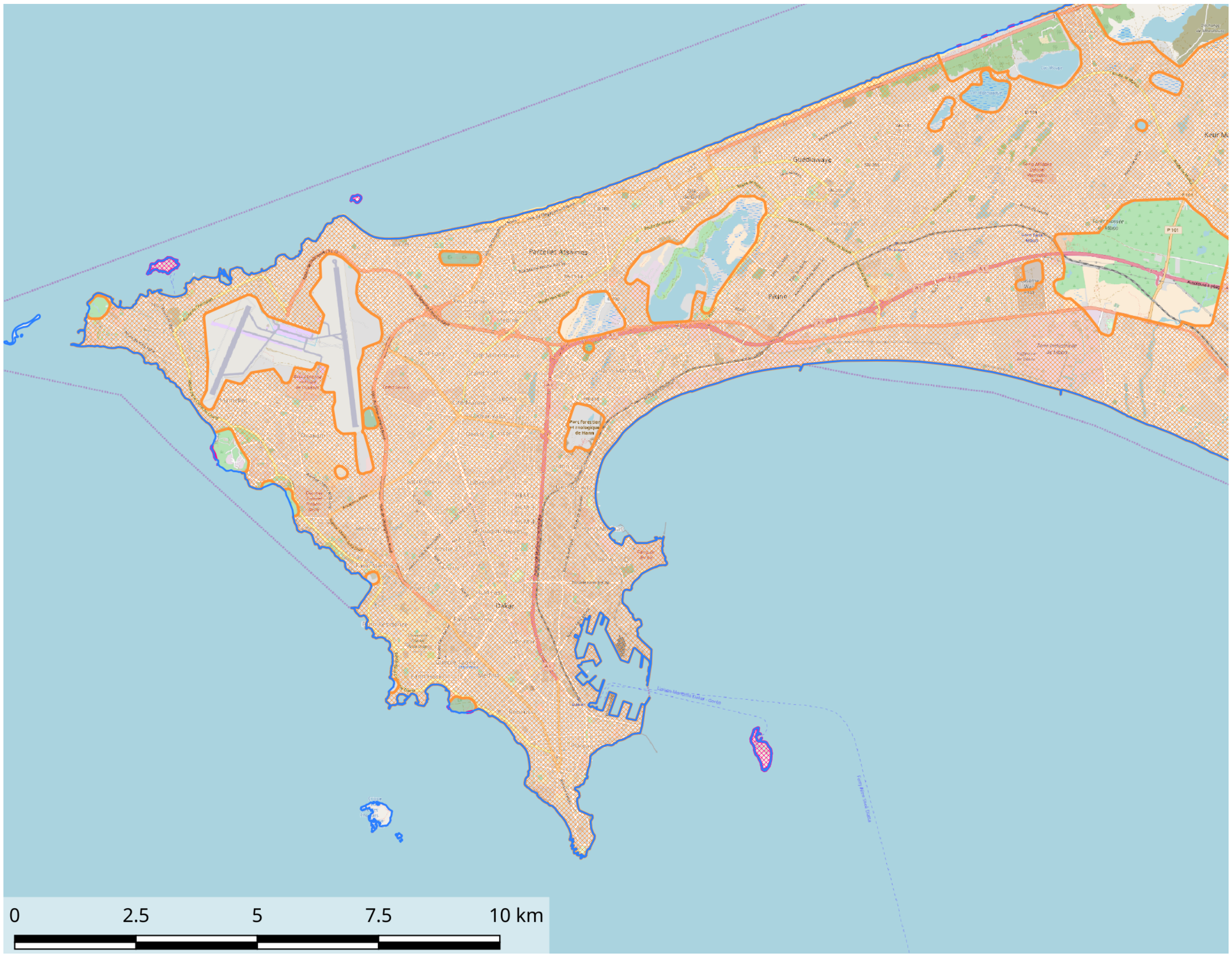

Figure 16: OpenStreetMap map of the Cap Vert Peninsula in 2018

#### 3 Final time series

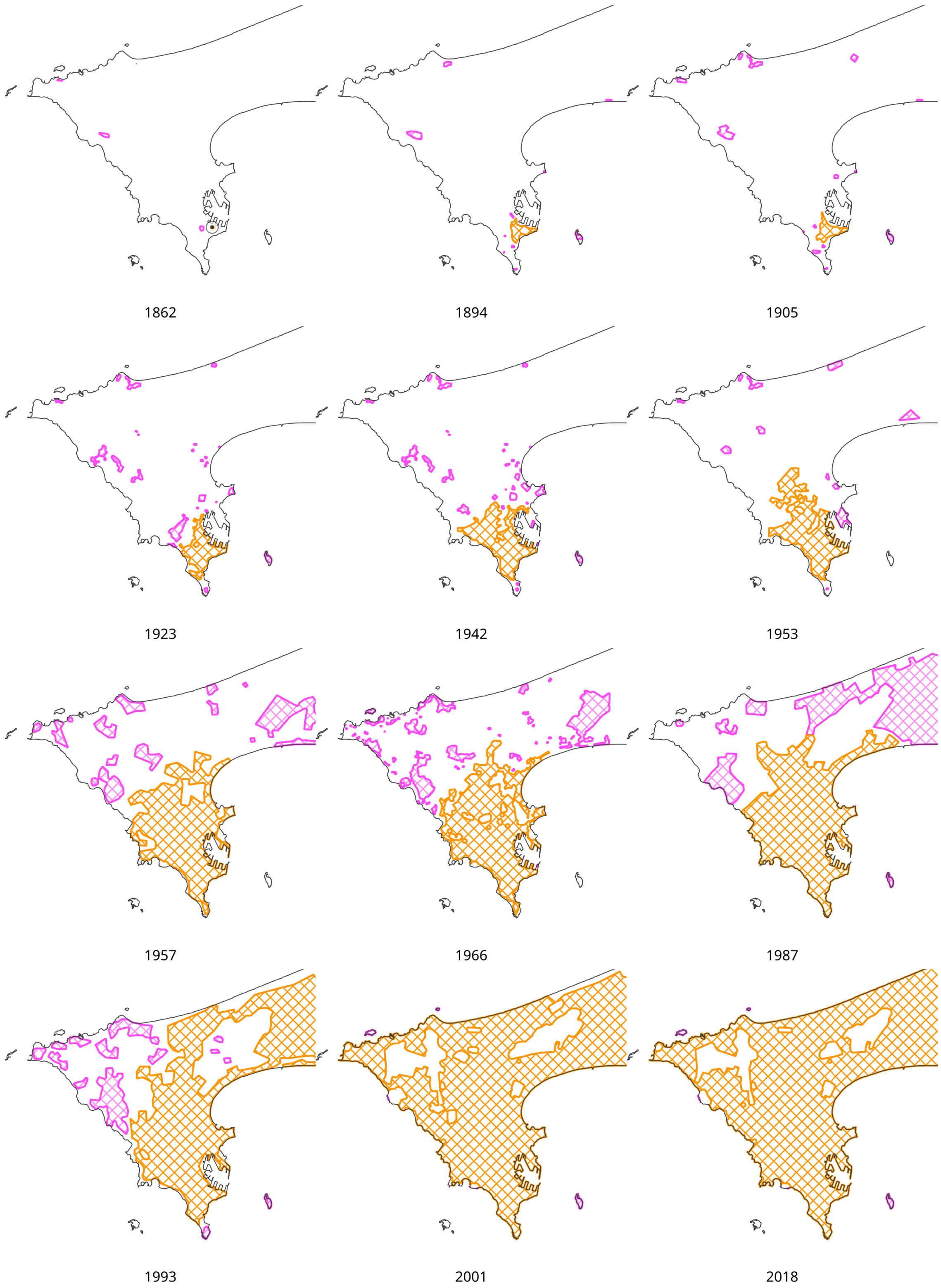

Figure 17: Time series illustrating the urbanization process of Dakar city. For each map, built-up areas are represented by orange or pink polygons depending on whether they are connected to the first European settlement or not, respectively.
